# A comprehensive overview and benchmarking analysis of fast algorithms for genome-wide association studies

**DOI:** 10.1101/2023.12.05.570105

**Authors:** Fang Liu, Jie Zhang, Yusheng Zhao, Renate H. Schmidt, Martin Mascher, Jochen C. Reif, Yong Jiang

**Author notes:** These authors contributed equally to this work (Fang Liu, Jie Zhang).

## Abstract

Genome-wide association studies (GWAS) are a ubiquitous tool for identifying genetic variants associated with complex traits in structured populations. During the past 15 years, many fast GWAS algorithms based on a state-of-the-art model, namely the linear mixed model, have been published to cope with the rapidly growing data size. In this study, we provide a comprehensive overview and benchmarking analysis of 33 commonly used GWAS algorithms. Key mathematical techniques implemented in different algorithms were summarized. Empirical data analysis with 12 selected algorithms showed differences regarding the identification of quantitative trait loci (QTL) in several plant species. The performance of these algorithms evaluated in 10,800 simulated data sets with distinct population size, heritability and genetic architecture revealed the impact of these parameters on the power of QTL identification and false positive rate. Based on these results, a general guide on the choice of algorithms for the research community is proposed.

## Introduction

Genome-wide association studies (GWAS) are an important tool for dissecting the genetic architecture of complex traits in human, animal and plant populations ^1, 2, 3^. Population structure and genetic relatedness affect the accuracy of GWAS and usually cause inflated test statistics resulting in high false positive rate (FPR). Early approaches to reduce inflation include genomic control ^4^, structured association ^5^ and principal component analysis ^6^. The Q+K linear mixed model (LMM) ^7^ has become the gold standard for GWAS because it is assumed to strike a good comprise between FPR and statistical power ^8, 9^. In this model, population structure is controlled by covariates (Q) and the kinship relatedness is accounted for by random polygenetic effects with a covariance matrix (K) derived from pedigree or genomic data.

A standard GWAS algorithm based on the Q+K model involves two steps: 1) solving the LMM and 2) generating the test statistics. These two steps are repeated for each marker (Fig. 1A). The first step is usually implemented by the maximum likelihood (ML) or restricted maximum likelihood (REML) method ^10^, both requiring iterations to estimate the unknown parameters. The most time-consuming parts are matrix multiplications and inverting matrices of size *n* × *n*, where *n* is the number of genotypes. Without specific mathematical techniques to improve the efficiency, the time complexity of this step is about *O*(*tpn*^3^), where *p* is the number of markers and *t* is the average number of iterations. The second step is usually performed by the likelihood ratio (LR) test, the Wald test or the F-test. The calculation of the LR test statistic is straightforward as the likelihood values are obtained in the first step. The Wald test or F-test statistics require extra computations involving multiplications of *n* × *n* matrices with *n*-dimensional vectors for each marker and the complexity is *O*(*pn*^2^). Thus, the complexity of the entire GWAS algorithm is dominated by the first step, namely *O*(*tpn*^3^). As the size of datasets increases rapidly with the advances in genomics technology, computational efficiency has become a bottleneck for the standard algorithm. Therefore, it is indispensable for the research community to improve the efficiency of GWAS algorithms without losing power or inflating FPR.

**Figure 1.**
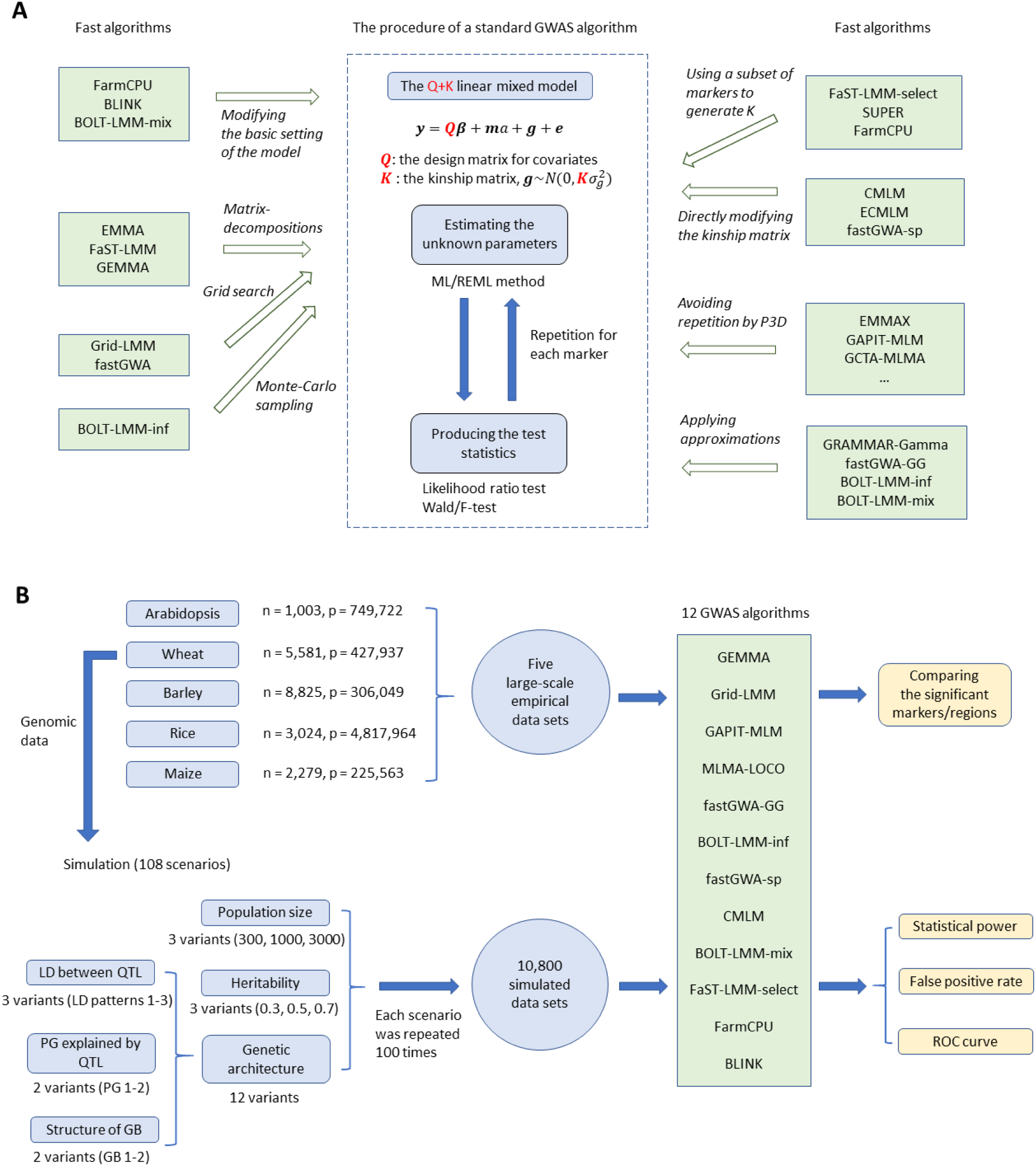
**A.** The principles of a standard GWAS algorithm based on linear mixed models and an illustration of the mathematical techniques applied by the fast algorithms. ML, maximum likelihood; REML, restricted maximum likelihood; P3D, population parameters previously determined. **B.** An outline of the strategy of our benchmarking analysis. LD, linkage disequilibrium; PG, proportion of genetic variance; GB, genetic background; ROC, receiver operating characteristic. n, the number of individuals; p, the number of markers.

In the last decade and a half, many different fast GWAS algorithms have been developed. Some applied elegant mathematical techniques to accelerate the standard algorithm ^11, 12, 13, 14, 15, 16^, others introduced different approximations in solving the LMM and/or generating test statistics ^14, 17, 18, 19, 20, 21, 22^, and still others modified the standard Q+K model with the aim of increasing the statistical power ^23, 24, 25, 26, 27, 28, 29^(Fig. 1A). These algorithms do not necessarily yield consistent results for the same data set. In 2014, a study compared several methods using human family-based data and highly concordant results were found for the different approaches ^30^. However, the underlying algorithms of the methods included in this study were similar. Since then, many new algorithms implementing diverse mathematical techniques have appeared and an up-to-date comprehensive comparison is lacking. Therefore, a deep understanding how the algorithms shape QTL identification is required to interpret the outcome of different GWAS algorithms for the same data set.

A practical problem faced by scientists is how to choose an appropriate algorithm for their research. In this study, we first review GWAS algorithms commonly used by the research community in the past decade and a half. Then, we select 12 representatives to perform a comprehensive benchmarking analysis in which the statistical power and false-discovery rate of the algorithms are compared through a large-scale simulation study with 10,800 data sets. In the end, we provide practical recommendations to scientists on the pros and cons of these algorithms.

## Results

### An overview of fast GWAS algorithms based on LMM

The computational complexity of fast GWAS algorithms is dominated by the step of solving the LMM. Consequently, the first few fast algorithms aimed directly at improving the efficiency of this step. The algorithm EMMA ^11^, also implemented in the software TASSEL ^31^, avoided repeatedly inverting matrices in each iteration by applying spectral decomposition to certain *n* × *n* matrices and reduced the time complexity from *O*(*tpn*^3^) to *O*(*pn*^3^ + *tpn*). However, the decomposition had to be applied for each marker in EMMA. This was further improved by two algorithms FaST-LMM ^12^ and GEMMA ^13^, in which the decomposition was performed only once for the kinship matrix (the *n* × *n* covariance matrix of random polygenic effects in the Q+K model) throughout the whole testing procedure. Then, the complexity was reduced to *O*(*n*^3^ + *pn*^2^ + *tpn*), where the *pn*^2^ term came from generating the marker-derived kinship matrix. These two algorithms produced exact test statistics up to machine precision and possible convergence to local maxima. Hence, they were classified as exact algorithms. Recently, another exact method MM4LMM ^16^ exploiting a minorize-maximization algorithm ^32^ to solve the LMM was published. Instead of solving the LMM by numerical optimization methods, Grid-LMM ^15^ directly searched for solutions in a pre-defined grid spanning all valid values with complexity *O*(*gn*^3^ + *pn*^2^), where *g* is the grid size. It was reported that the test statistics were almost the same as exact methods despite limiting the precision of estimators by the resolution of the defined grid. Thus, we may call it a quasi-exact algorithm.

In order to improve the computational efficiency further, approximations were introduced to the procedure of solving the LMM. The earliest approximated approach “Population Parameters Previously Determined” (P3D) ^17^ was implemented in the software GAPIT ^33, 34, 35^, termed GAPIT-MLM, and was independently invented in EMMAX ^22^. In this approach, the LMM was solved only once for the “null model”, i.e., the model without any marker effects. Then, the estimators of variance components were fixed throughout the whole testing procedure. In this way, it avoided repeatedly solving the LMM for each marker and hence further reduced the complexity to *O*(*n*^3^ + *pn*^2^ + *tn*). It was reported that the P3D approach resulted in similar detection power to the exact methods. Hence, this approach has been implemented either by default or as an option in almost all popular software packages for GWAS (e.g. FaST-LMM, GCTA ^36^, TASSEL).

When P3D is implemented, producing test statistics requires additional computations with complexity *O*(*pn*^2^), no matter which type of statistical test is applied. Hence, producing the test statistics in a more efficient way can also increase the efficiency. The algorithm GRAMMAR ^19^ implemented in GenABEL ^37^ proposed to use the residuals from the null model as the response in a simple linear regression model to test the marker effects. This approach reduced the complexity of calculating test statistics from *O*(*pn*^2^) to *O*(*pn*), but produced biased test statistics with reduced power. This drawback was overcome in GRAMMAR-Gamma ^20^, which introduced an approximation to the original Wald-test statistic and also reduced the complexity to *O*(*pn*). This approach was also included as an option in the algorithm fastGWA ^21^ implemented in GCTA, which is termed fastGWA-GG in our study. Nevertheless, for these algorithms the complexity is still *O*(*n*^3^ + *pn*^2^ + *tn*) since the total complexity is dominated by the step of solving the LMM.

Although applying decompositions to the kinship matrix greatly improves the computational efficiency, the complexity of the technique itself is still high, typically *O*(*n*^3^). Therefore, some algorithms tried to improve the efficiency by modifying the kinship matrix. The algorithms CMLM ^17^ and its enrichment version ECMLM ^18^ in GAPIT compressed the kinship matrix by classifying the genotypes into *c* groups (*c* ≪ *n*), so that the computational efficiency increased by (*n*/*c*)^3^ fold. However, to optimize the detection power, the evaluation of the likelihood function needed to be repeated until the parameter *c* is optimized. The algorithm fastGWA provided the option of setting a threshold to make the kinship matrix sparse, namely all entries below the threshold were set to zero, so that special techniques for sparse matrices can be used to increase efficiency. This variant of fastGWA is termed fastGWA-sp in our study. When both the sparse kinship matrix and the GRAMMAR-Gamma approximation are used, the variant is termed fastGWA-sp-GG.

The first algorithm that avoids decomposing the kinship matrix is BOLT-LMM ^14^, which implemented a Monte Carlo sampling approach to solve the LMM. In this algorithm, it was not necessary to explicitly calculate the marker-derived kinship matrix, and it also avoided inverting or decomposing matrices of size *n* × *n*. Indeed, it transformed the problem to solving systems of linear equations by the conjugate gradient method, in which only products of *n* × *p* matrices with *p*-dimensional vectors are needed (at the cost of a few iterations). Together with P3D, it remarkably reduced the complexity of solving LMM to *O*(*mtpn*), where *m* is the average number of Monte Carlo sampling and *t* is the average number of iterations. BOLT-LMM also invented an approximated approach for calculating the test statistics similar to GRAMMAR-Gamma. Therefore, the total complexity of BOLT-LMM is *O*(*mtpn*). This algorithm has two variants, one follows the standard Q+K model (BOLT-LMM-inf), the other assumed Gaussian mixture priors for the random marker effects which serve as a control of polygenic background, termed BOLT-LMM-mix ^21^. By default, BOLT-LMM combined the two variants and applied cross-validations to determine which variant was used to produce the final test statistics.

Most of the above algorithms aim to improve the computational efficiency of the standard Q+K model, while others modify the model to increase the power and/or to decrease the FPR. The first technique with such a purpose was “Leave-One-Chromosome-Out” (LOCO) ^38^, which has already been implemented by default or as an option in many algorithms (FaST-LMM, MLMA-LOCO in GCTA, BOLT-LMM, REGENIE ^29^). That is, when a marker is tested, all markers on the same chromosome are excluded when calculating the kinship matrix. It has been reported that LOCO can increase the power by avoiding proximal contamination, a phenomenon that the power of detecting QTL is reduced when markers correlated with the QTL are involved in the calculation of kinship matrix ^23, 38^. The algorithm MLMM ^25^ implemented a forward-backward stepwise linear mixed model to include additional marker covariates in the Q+K model. While MLMM still used all markers to build up the kinship matrix, FaST-LMM-select ^23^ used cross-validation to select a subset of markers for deriving the kinship matrix. FaST-LMM-select was reported to have higher power than the standard Q+K model, but in some cases did not sufficiently control the FPR. This was improved in FaST-LMM-all+select ^24^, in which two random polygenic effects were included, one with the kinship matrix derived by all markers and the other by the selected ones. In the software package GAPIT, three different algorithms SUPER ^26^, FarmCPU ^27^ and BLINK ^28^ were implemented, all of which differ from the standard Q+K model. A common feature of these three algorithms is that only a subset of markers was selected to control the population structure. In SUPER and FarmCPU, the selected markers were used to generate the kinship matrix. In BLINK, the selected markers were modeled directly as fixed covariates. The models used to produce the test statistics were also different: SUPER still used LMM, but FarmCPU and BLINK used the multi-variate linear regression (MLR) model. A recently published algorithm REGENIE ^29^ tested the marker effects based on the residuals from the null model, similar to GRAMMAR. But it fitted the null model by a two-step stacked ridge regression approach, which is different from all above algorithms.

The main features of the above algorithms were summarized in Table 1 and the relationship among these algorithms was illustrated in Extended Data Fig. 1. We selected 12 representatives for the subsequent benchmarking analysis: GEMMA, Grid-LMM, GAPIT-MLM, MLMA-LOCO, CMLM, fastGWA-GG, fastGWA-sp, BOLT-LMM-inf, BOLT-LMM-mix, FaST-LMM-select, FarmCPU and BLINK. These algorithms encompass the most important mathematical techniques which were applied to improve the computational efficiency. In case many algorithms implemented essentially the same techniques, only one representative was selected. More details of the selection procedure were described in Methods. An in-depth review of the key mathematical techniques implemented in the 12 selected algorithms was provided in Supplementary Note A.

**Table 1.**
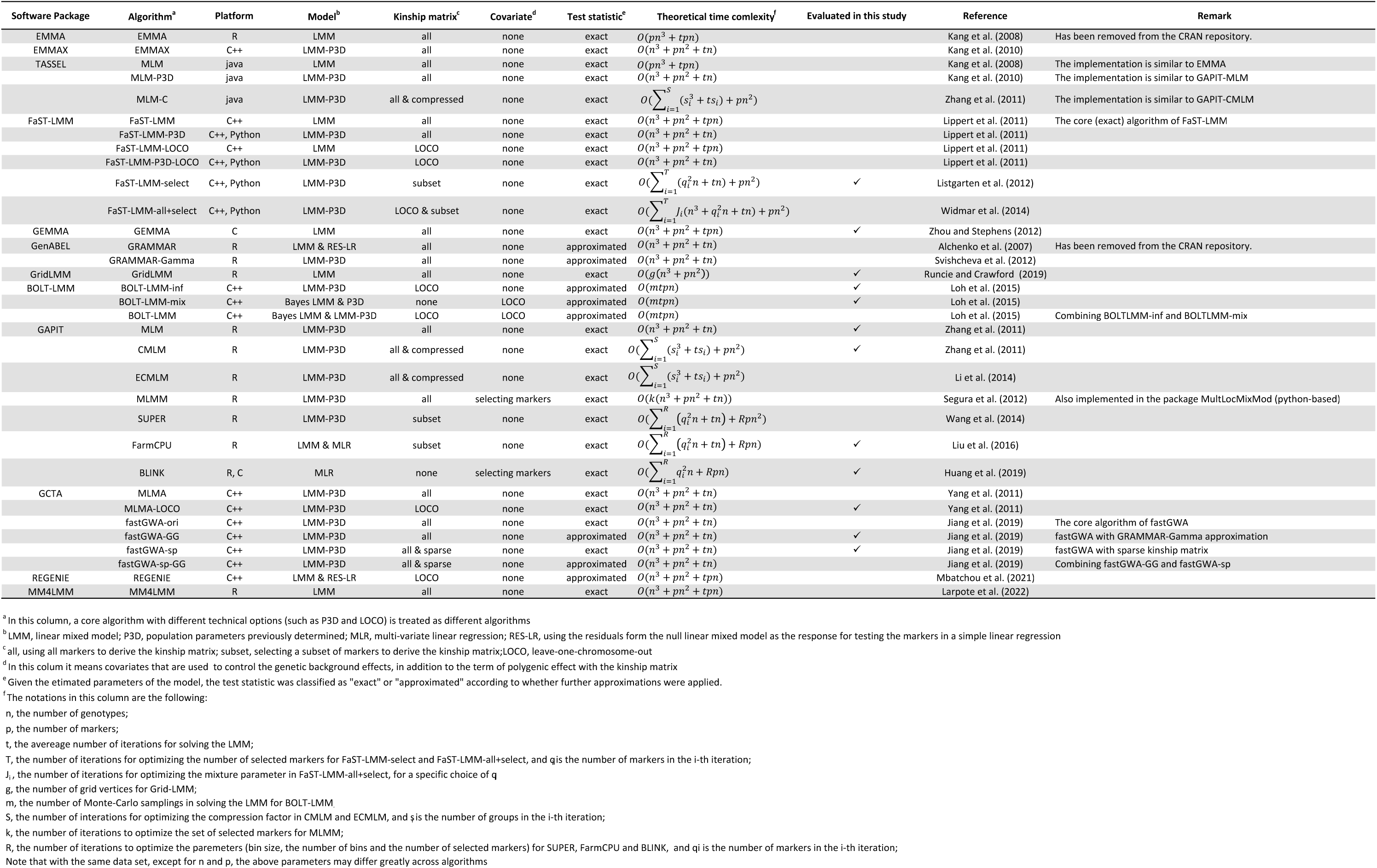
Summary of commonly applied algorithms for genome-wide association studies.

### The strategy of the applied benchmarking analysis

Our benchmarking analysis consisted of two parts based on empirical and simulated data sets, respectively (Fig. 1B). The empirical data sets comprised four large plant populations of inbreeding species: Arabidopsis, wheat, rice, barley as well as one population of maize inbred lines (see Methods for details). The 12 selected algorithms were applied to each of the five datasets, with a few exceptions as the computational load for some algorithms in certain data sets was too high.

The simulated phenotypic data were based on the genomic data from a wheat population consisting of 5,581 accessions with 427,937 single nucleotide polymorphism (SNP) markers. Three population sizes (300, 1000, 3000), three trait heritabilities (0.3, 0.5, 0.7), and 12 different levels of complexity for the genetic architecture were considered in the simulation. Thus, there were in total 3 × 3 × 12 = 108 different scenarios. Each scenario was simulated 100 times, resulting in 10,800 data sets. For the simulated genetic architecture, we considered three factors of complexities. 1) The extent of linkage disequilibrium (LD) between QTL (three levels, denoted by LD pattern 1, 2 and 3). 2) The proportion of genetic variance (PG) explained by the major QTL (two levels, indicated by PG1 and PG2). 3) The number of minor QTL contributed as the genetic background (GB) effects (two levels, designated GB1 and GB2) (Figure 1B). More details of the simulation procedure were described in Methods.

In each of the 108 scenarios, the statistical power in detecting QTL and the FPR of the 12 algorithms were assessed through the 100 replicated datasets, except for CMLM and FaST-LMM-select, which were solely evaluated in the 72 scenarios representing the datasets with population sizes 300 and 1,000 because the computational load was too high for a population size of 3,000.

### Comparing the performance of the algorithms with empirical data

The 12 selected algorithms were compared based on five empirical data sets (Fig. 2; Supplementary Figs. 1-4). We took the result of GEMMA as a benchmark as it is an exact algorithm based on the standard Q+K model. In general, we found that GEMMA detected the least number of regions among all algorithms, except fastGWA, which detected fewer regions than GEMMA in the wheat, Arabidopsis and maize datasets (see Panel D of the five figures). Nevertheless, the regions identified by GEMMA were the most congruent among all algorithms. In particular, the results of Grid-LMM, GAPIT-MLM and CMLM were nearly identical to those of GEMMA (the correlation of -log_10_(*p*) values was close to 1). The regions identified by GEMMA were also detected by BOLT-LMM-inf and BOLT-LMM-mix in most cases, but not always by FaST-LMM-select, fastGWA-sp, FarmCPU or BLINK. Note, that the regions detected by FarmCPU and BLINK were typically represented as scatter points rather than as peaks in the Manhattan plots reflecting the underlying model (Supplementary Note A). Except for the barley dataset, regions commonly identified by other algorithms but not GEMMA were detected by five or fewer algorithms, and many were detected by only one or two algorithms. For example, in the Arabidopsis data set, the 12 algorithms identified in total 20 regions, whereas GEMMA only found three. And there were 14 regions which were identified by just one algorithm (Supplementary Fig. 1). Interestingly, candidate genes whose role in controlling flowering time had been documented in literature were found closely linked to the associated SNPs for 16 out of 20 regions (Supplementary Table 1). Nevertheless, since the positions of true QTL are largely unknown for the empirical data sets, it is unclear which of the QTL represent faithful candidates or false positives. A reliable assessment of power and FPR of the different algorithms requires a simulation study.

**Figure 2.**
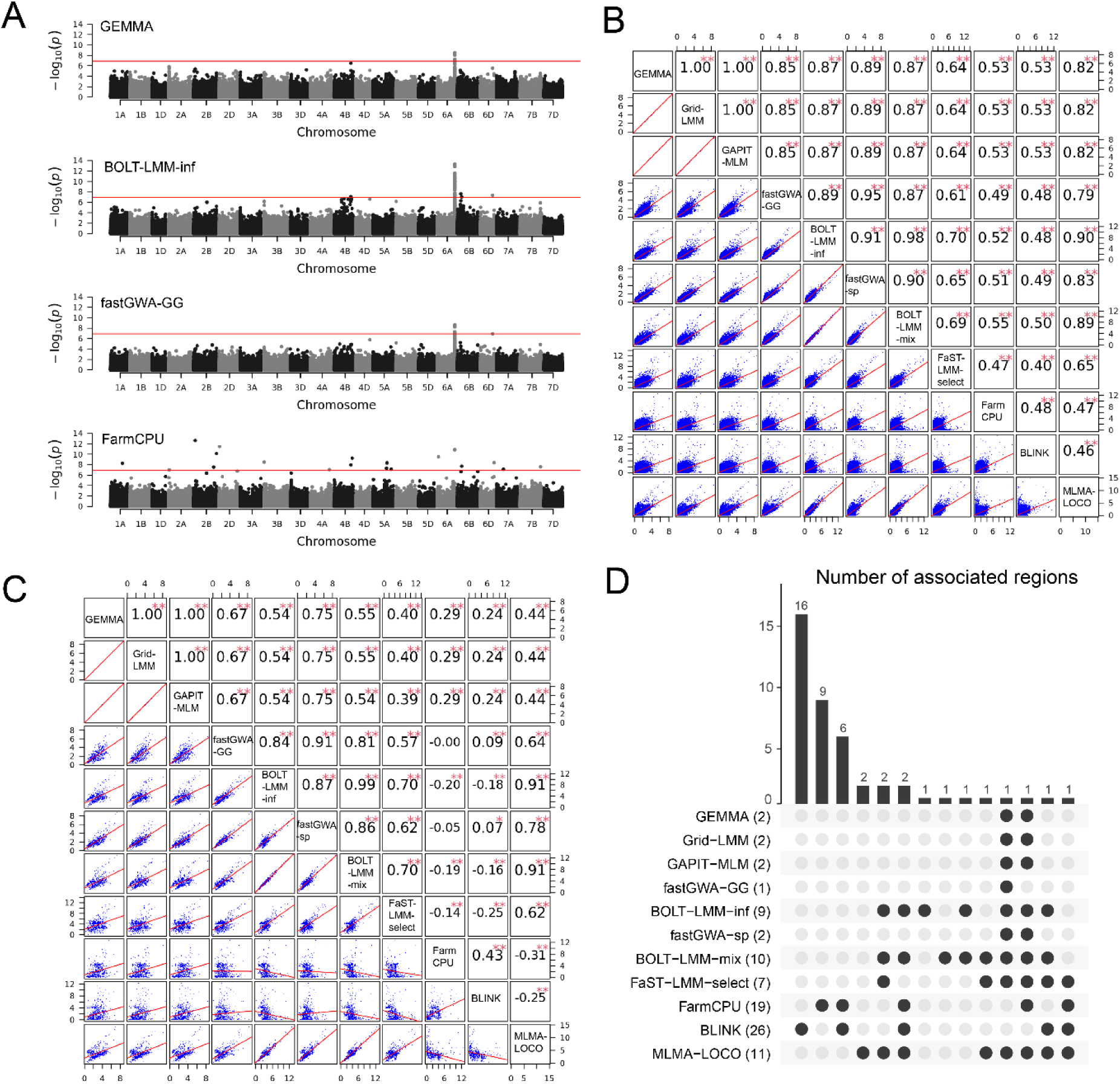
A comparison of the results of 11 GWAS algorithms for the resistance to yellow rust in a wheat data set consisting of 5,581 individuals and 427,937 markers. **A.** The Manhattan plots of 4 selected algorithms. The threshold was *p* < 0.05 after Bonferroni correction. **B.** Correlations between the -log_10_(*p*) values of all markers obtained by each of the indicated pairs of algorithms. The names of the algorithms were indicated in the diagonal blocks. **C.** Pairwise correlations between the -log_10_(*p*) values of markers which were significant under a liberal threshold (-log_10_(*p*) > 4) in at least one algorithm. **D.** A comparison of significant regions identified by the 11 algorithms. The bar plot showed the number of regions commonly identified by the algorithms indicated by the black dots. The number of regions identified by each algorithm was presented in the parentheses next to the names of the algorithms. The algorithm CMLM was not applied to this data set due to the computational load.

### Comparing the statistical power and FPR of the algorithms with simulated data

We compared the power of QTL detection and FPR of all algorithms under the threshold of *p* < 0.05 after Bonferroni correction for multiple testing ^39^. In general, the heritability of the trait, the population size, and the number of markers contributing to the genetic background played a minor role in ranking the algorithms. MLMA-LOCO, FaST-LMM-select and the two variants of BOLT-LMM (BOLT-LMM-inf and BOLT-LMM-mix) achieved the highest power in most scenarios (Fig. 3A, Extended Data Figs. 2A, 3A). However, they also produced the highest FPR among all algorithms, while the FPR of the other 8 algorithms was much lower (Fig. 3B, Extended Data Figs. 2B, 3B).

**Figure 3.**
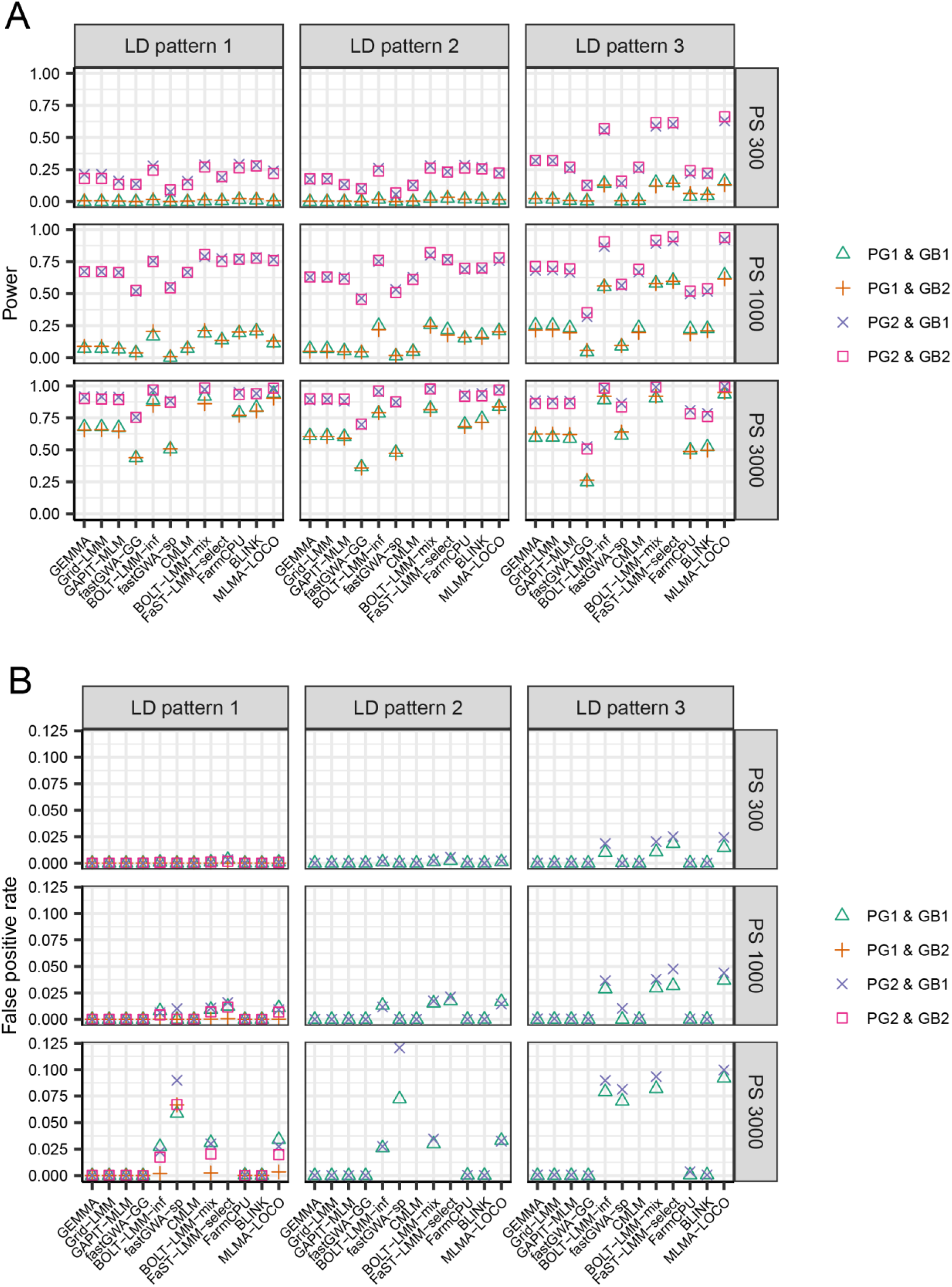
The statistical power (**A**) and false positive rate (**B**) of 12 GWAS algorithms evaluated in simulated data sets with 36 scenarios for trait heritability 0.7, under the threshold of *p* < 0.05 after Bonferroni correction for multiple testing. The 36 scenarios are combinations of three population sizes (PS 300, PS 1000 and PS 3000), three different linkage disequilibrium (LD) patterns among the QTL (LD patterns 1-3), two patterns of QTL effect sizes (PG1 and PG2), and two different genetic backgrounds (GB1 and GB2). In LD pattern 1, there is no LD between any two major QTL or between a major and a minor QTL. In LD pattern 2, there is no LD between any two major QTL, but LD exits between major and minor QTL. In LD pattern 3, there exists LD among the major QTL as well as between major and minor QTL. In PG1, each of the 6 major QTL explained 2% of the genetic variance. In PG2, the 6 major QTL were randomly assigned to explain 2%, 4%, 6%, 8%, 10% and 12% of the genetic variance respectively. In GB1, there were 1,200 markers as minor QTL. In GB2, all markers on the chromosomes (LD patterns 2 and 3) or on half of the chromosomes (LD pattern 1) contributed as minor QTL. Each of the 9 subpanels showed the results of a specific combination of population size and LD pattern. Within each subpanel, the results of four combinations of two PGs and two GBs were indicated by different symbols. For GB2, the FPR was only calculated in LD pattern 1, because for LD patterns 2 and 3, all markers contributed to the simulated trait either as major or as minor QTL. The algorithms CMLM and FaST-LMM-select were not evaluated for PS 3000 because the computational load was too high.

The power of GEMMA and Grid-LMM was very similar across all scenarios, as was the power of FarmCPU and BLINK. Interestingly, the relative performance of the two groups of algorithms depended on the genetic architecture of the datasets. More specifically, the power of FarmCPU and BLINK was higher than that of GEMMA and Grid-LMM when there was no LD between major QTL (LD patterns 1 and 2), especially in PG1 where each of the six QTL explained only 2% of the genetic variance. In contrast, when the QTL were in LD (LD pattern 3), GEMMA and Grid-LMM produced similar power to that of FarmCPU and BLINK in PG1, and substantially outperformed them in PG2, where the PG explained by the six QTL was much higher (from 2% to 12% with a step of 2%). Thus, the results indicated that 1) FarmCPU and BLINK are favored for independent QTL, while GEMMA and Grid-LMM are better for detecting QTL pairs in LD. 2) In the case of independent QTL, the advantage of FarmCPU and BLINK are more pronounced for QTL explaining small PG. These conclusions were supported by evidence from more detailed analyses: For scenarios with PG2, it is very clear that the power of FarmCPU and BLINK was much higher than that of GEMMA and Grid-LMM for the discovery of QTL with PG ≤ 6% in LD patterns 1 and 2, whereas for QTL with PG ≥ 8% their advantage was less evident (Extended Data Fig. 4, Supplementary Figs. 5-6). For scenarios with LD pattern 3, we observed that at low level of LD between QTL (0.16 < *r*^2^ ≤ 0.36), GEMMA and Grid-LMM had almost no advantage (Extended Data Fig. 5, Supplementary Fig. 7). Nevertheless, as LD increased, the power of GEMMA and Grid-LMM exceeded that of FarmCPU and BLINK. This trend was more pronounced in PG2 than in PG1.

Further investigations revealed that the two groups of algorithms also differed in their ability to detect QTL with different MAFs (Extended Data Figs. 6-7, Supplementary Figs. 8-11). For GEMMA and Grid-LMM, the difference between the power of detecting QTL with different ranges of MAF was larger than for FarmCPU and BLINK. For example, with LD patterns 1 and 2, the power of GEMMA and Grid-LMM for detecting QTL with MAF above 0.1 was clearly lower than that of FarmCPU and BLINK. However, for QTL with MAF less than 0.1, the gap was much smaller and in many scenarios GEMMA and Grid-LMM achieved similar power as FarmCPU and BLINK. This trend was even more pronounced with LD pattern 3. In most scenarios, the power of GEMMA and Grid-LMM was similar to or only slightly higher than FarmCPU and BLINK for QTL with MAF above 0.1, but for QTL with MAF below 0.1, FarmCPU and BLINK were clearly outperformed. These results indicate that GEMMA and Grid-LMM are more sensitive to the MAF of QTL and are better at detecting QTL with rare alleles, while FarmCPU and BLINK are more powerful at detecting QTL with common alleles.

For the remaining four algorithms, the power of GAPIT-MLM and CMLM was similar to or slightly lower than GEMMA and Grid-LMM in most scenarios, followed by the two variants of fastGWA (fastGWA-sp and fastGWA-GG). The power of fastGWA-GG was the lowest in most scenarios. Detailed analysis indicated that the low power was likely due to the underlying model of fastGWA which is slightly different from the standard Q+K model (Supplementary Note B). In addition, we found that in some scenarios (e.g., LD patterns 1 and 2, population size 3000 and heritability 0.7, Fig. 3B), the FPR of fastGWA-sp was surprisingly high. Further analysis revealed that for a small fraction of simulated datasets, the *p*-value produced by fastGWA-sp were zero for all markers, and we suspected that this might be caused by cumulated numerical errors during the computation (Supplementary Note B).

The above comparisons of the performance of the 12 algorithms are based on a common threshold. To assess their overall ability to classify true and false positives, we investigated the receiver-operating characteristic (ROC) curves (Fig. 4, Extended Data Figs. 8-9), which is obtained by depicting the power against the FPR under various thresholds. Since fastGWA-sp produced erroneous *p*-values in a small proportion of data sets, it was excluded from this part of analysis. Surprisingly, the ranks of the algorithms from the viewpoint of ROC curves differed from those under a fixed threshold. We found that the ROC curves of GEMMA, Grid-LMM, GAPIT-MLM and CMLM overlapped almost completely and were closest to the upper-left corner, or point (0, 1) in all scenarios. Thus, the area under the curve (AUC) was largest for these four algorithms, implying that they clearly outperformed the other algorithms in the sense that they would have the highest power at any given level of FPR and the lowest FPR at any given power. It should be noted, however, that it is not easy to exploit this theoretical advantage in reality, because different algorithms reach a given FPR or power at different thresholds, and for empirical data sets it is impossible to know the exact relationship between the threshold and power/FPR. For a given threshold, the algorithm favored by the ROC curve does not necessarily produce the highest power. This is exactly what we observed in Fig. 3 with our simulated datasets for which a stringent threshold (*p* < 0.05 after Bonferroni correction) was applied. We also tried a more liberal threshold (*p* < 0.05 after Benjamini-Hochberg correction ^40^) and found that the rankings of the algorithms did not change in most scenarios (Supplementary Figs. 12-14). Therefore, it would be very interesting to know if there is an optimized threshold for the algorithm whose ROC curve has the largest AUC, so that the theoretical advantage can be exploited. This topic is beyond the scope of the current study, but is certainly worth further investigation.

**Figure 4.**
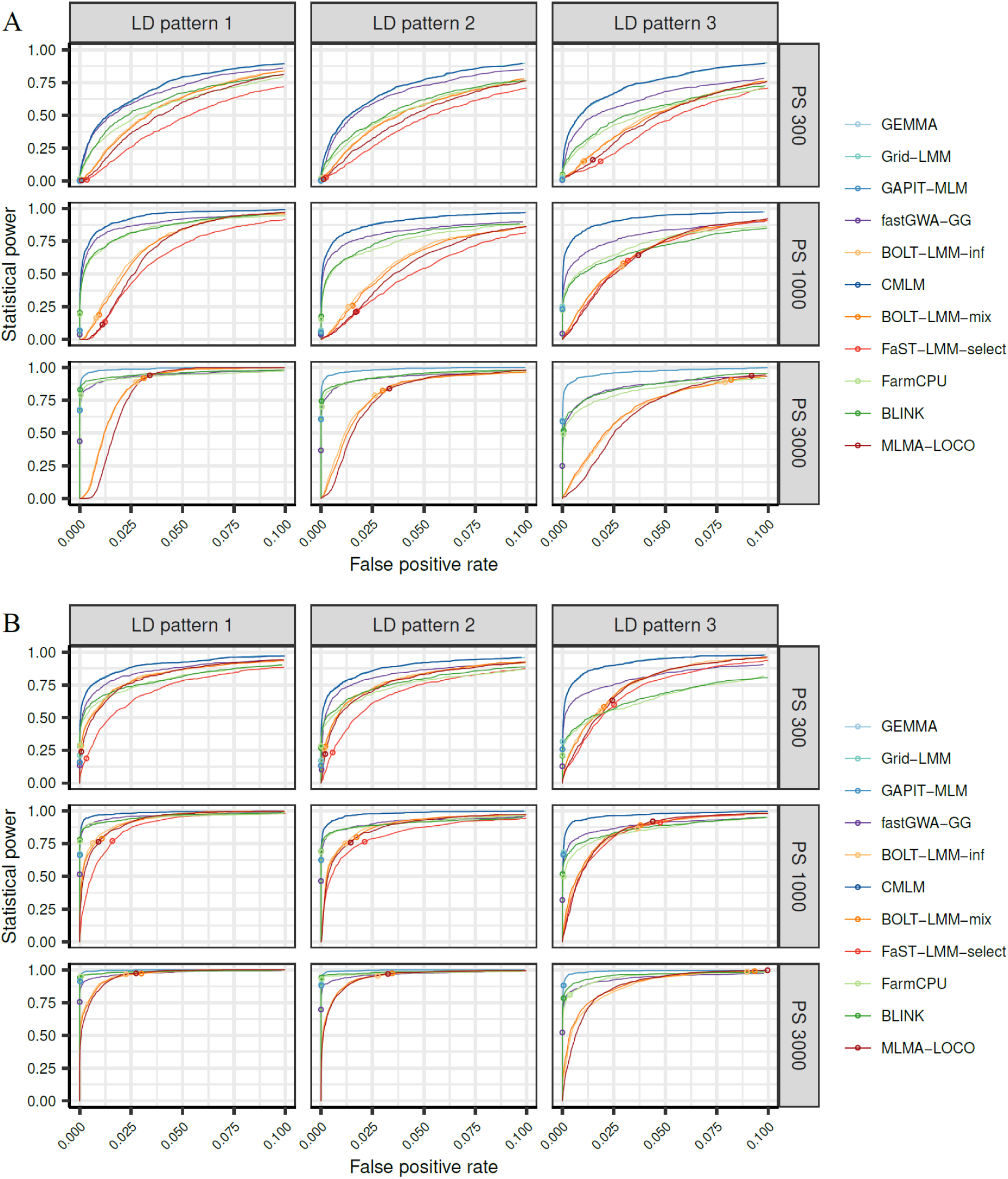
The receiver operating characteristic (ROC) curves of 11 GWAS algorithms evaluated in simulated data sets with 18 scenarios for trait heritability 0.7 with GB1 (1,200 markers contributed as minor QTL to the genetic background effects). The 18 scenarios are combinations of three population sizes (PS 300, PS 1000 and PS 3000), three different linkage disequilibrium (LD) patterns among the QTL (LD patterns 1-3), and two patterns of QTL effect sizes (PG1 and PG2). In LD pattern 1, there is no LD between any two major QTL or between a major and a minor QTL. In LD pattern 2, there is no LD between any two major QTL, but LD exits between major and minor QTL. In LD pattern 3, there exists LD among the major QTL as well as between major and minor QTL. In PG1, each of the 6 major QTL explained 2% of the genetic variance, In PG2, the 6 major QTL were randomly assigned to explain 2%, 4%, 6%, 8%, 10% and 12% of the genetic variance respectively. Results for PG1 and PG2 were shown in panels **A** and **B**, respectively. Each of the 9 subpanels showed the results for a specific combination of population size and data set. Within each subpanel, the ROC curves of different algorithms were shown in different colors. The power and FPR of each algorithm under the threshold of *p* < 0.05 after Bonferroni correction for multiple testing was indicated by a small circle on the curve. The algorithms CMLM and FaST-LMM-select were not evaluated for PS 3000 because the computational load was too high.

### The influence of specific techniques on the power and FPR

The results obtained in the previous section enabled a detailed investigation of the influence of a specific mathematical/statistical technique on the detection power and FPR by comparing the results of two algorithms that differ only in whether the technique is implemented or not (Fig. 5A). In the following, we mainly focused on the techniques inflating FPR, because the four algorithms that boosted power (FaST-LMM-select, BOLT-LMM-inf, BOLT-LMM-mix and MLMA-LOCO) were also accompanied with inflated FPR.

**Figure 5.**
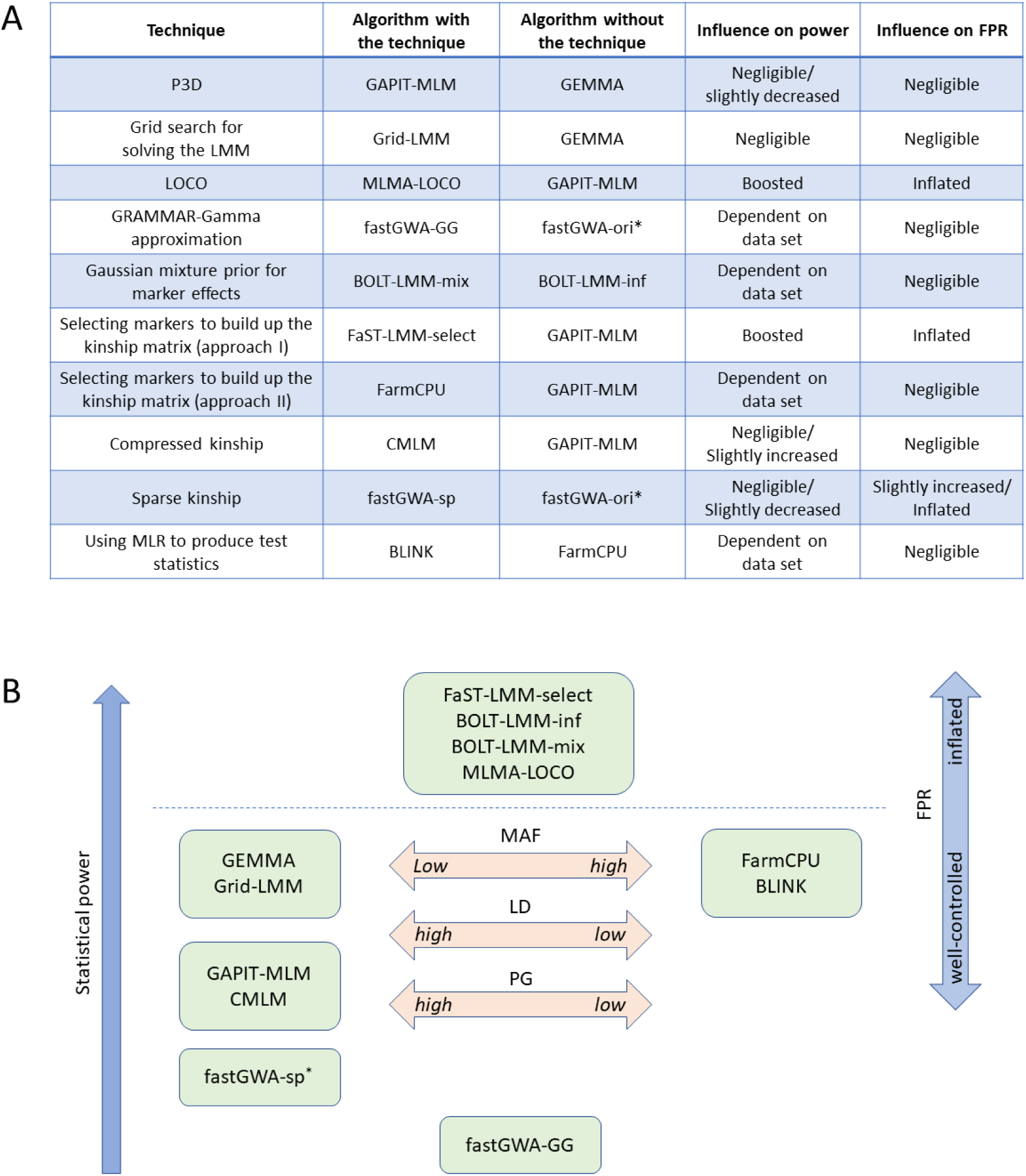
**A.** Summary of key mathematical techniques implemented in the fast GWAS algorithms evaluated in this study and their influence on power and false-positive rate (FPR). For each technique, the results were obtained by comparing two algorithms differing only in having the technique implemented (listed in the second column) or not (listed in the third column). P3D, population parameters previously determined; LMM, linear mixed model; LOCO, leave-one-chromosome-out; MLR, multi-variate linear regression. * fastGWA-ori is the original algorithm of fastGWA without implementing the GRAMMAR-Gramma approximation or the sparse kinship matrix. It was not evaluated in the benchmarking analysis (see Supplementary Note B). **B.** A brief illustration of the results of benchmarking analysis for the 12 GWAS algorithms. Algorithms above the dashed line as well as fastGWA-sp (indicated by the ∗ symbol) had inflated FPR, while others stringently controlled the FPR. In general, the altitude of the algorithms indicated the level of their statistical power. The three arrows in the middle indicate algorithms with power advantage for specific types of QTL. Algorithms next to the left arrow are better at detecting QTL with low minor allele frequency (MAF), high proportion of genetic variance (PG) and high linkage disequilibrium (LD) with each other. Algorithms next to the right arrow are better at detecting QTL with high MAF, low PG and low LD with each other.

FaST-LMM-select specifies a subset of markers whose correlations with the trait are highest to build up the kinship matrix. In this process, the number of markers is determined by cross-validation. We found that this approach boosted the power but also inflated the FPR, which is consistent with previous studies ^24, 38^. It has been reported that FaST-LMM-all+select or adding a few PCs of the SNP matrix as covariates to the FaST-LMM-select model can control FPR ^24^. Nevertheless, we did not evaluate these two approaches as they significantly increase the computational load.

MLMA-LOCO implemented LOCO and P3D, BOLT-LMM-inf and BOLT-LMM-mix implemented LOCO, P3D and introduced certain approximations for computing test statistics. We observed that these three algorithms increased both the power and the FPR. Since there is no evidence that P3D inflates the test statistics, it suggests that LOCO is responsible for the inflated FPR. A previous study also observed inflated FPR for BOLT-LMM-inf and BOLT-LMM-mix ^21^ and claimed that the high FPR was due to a partial LOCO approach implemented in these two algorithms (The LOCO technique was only applied in the calculation of the test statistics, but not in the estimation of the unknown parameters of the LMM). We then re-evaluated the two algorithms by forcing a genuine LOCO procedure (see Methods) with 400 simulated data sets, but could not find essential differences (Supplementary Table 2). Thus, our results indicated that LOCO not only increases the power but also caused inflation of FPR. It is well-known that LOCO can avoid proximal contamination and hence increase the statistical power compared with the algorithms using all markers to build up the kinship matrix ^23, 38^. However, recent studies reported inflated test statistics when LOCO was applied ^41, 42^, thus providing an explanation for the inflated FPR which was observed in this study.

The sparse kinship technique implemented in fastGWA-sp also inflated FPR. Even if we ignore the small proportion of the simulated data sets in which computational errors occurred, the FPR of fastGWA-sp in certain scenarios was still higher than other algorithms (Supplementary Note B). In our analysis, the threshold for sparse kinship was set to 0.05, which was recommended by the algorithm. We also examined the performance of fastGWA-sp with other thresholds (0, 0.1, 0.15, and 0.2) in 100 simulated datasets and observed different levels of inflation of FPR (Supplementary Fig. 15). It should be noted that sparse kinship with the threshold 0 is not the same as exact kinship because negative values exist in the kinship matrix. This result indicated that setting small entries in the kinship matrix to zero may lead to insufficient control of the population structure. Therefore, further studies are needed to find out the applicability of this technique in different populations.

## Discussion

In this study, we provided a comprehensive overview of LMM-based fast GWAS algorithms applied in the last decade and a half and selected 12 representatives for a benchmarking analysis to evaluate their statistical power and FPR using 10,800 simulated datasets. Large plant populations of inbreeding species or inbred lines were the focus as they were often underrepresented in previous studies in which the algorithms were developed and evaluated. For example, some algorithms were developed and tested only with human data sets ^14, 21^ or with small-size plant data sets ^27, 28^. Indeed, we observed some results different from those obtained in simulation studies based on human genomic data. The influence of LOCO and sparse kinship matrix on the FPR are two examples. While we found inflated FPR for algorithms implementing these two techniques, studies based on human populations did not ^38, 43^. Interestingly, it was also reported that LOCO could result in inflated test statistics (the genomic inflation factor *λ* > 1, defined as the median of the observed distribution of test statistics divided by the median of the expected distribution) in some empirical studies with animal populations ^41, 42^ and it was suspected that the stronger population stratification in livestock populations might be the reason why inflation was not observed in studies with human populations ^41^. Therefore, the influence of a specific technique on the power and FPR of GWAS might not be consistent across species, or even populations. Considering this point, we evaluated the 12 algorithms in 1,200 additional simulated data sets based on the genomic data of Arabidopsis, maize and barley populations. The rankings of algorithms in terms of statistical power and FPR were consistent with those observed for wheat genomic data (Extended Data Fig. 10). Nonetheless, the differences between classes of algorithms in terms of power and FPR was less pronounced in the Arabidopsis and maize data sets compared to the results for genomic data of wheat and barley. In future, it should be a priority to assess whether for example differences in LD decay contribute to the observed inconsistencies. For the time being, the selection of GWAS algorithms should consider potential differences between species and populations. If one or more techniques implemented by an algorithm were reported to have inflated test statistics in certain populations, we should be careful to apply it and at least check the genomic inflation factor with the resulting test statistics.

The results of our benchmarking analysis were summarized in Fig. 5B. FaST-LMM-select, BOLT-LMM-inf, BOLT-LMM-mix and MLMA-LOCO had the highest power but also the highest FPR across all scenarios, while fastGWA-sp had inflated FPR in some scenarios. Thus, the additional regions identified by these five algorithms in the empirical datasets could be a mix of true and false positives. In general, we would suggest being cautious when applying the five algorithms. However, for small populations and traits with very low heritability, FaST-LMM-select, BOLT-LMM-inf, BOLT-LMM-mix and MLMA-LOCO might be a good choice because the four algorithms had much higher power than the other algorithms and their FPR was still in an acceptable range.

The other seven algorithms controlled the FPR stringently. While fastGWA-GG produced the lowest power in most scenarios, the remaining six can be divided into two groups: The first group consists of GEMMA, Grid-LMM, GAPIT-MLM and CMLM, and the second is comprised of FarmCPU and BLINK. Note that additional candidates for the first group include FaST-LMM (without LOCO), EMMAX, GCTA-MLMA (without LOCO) and TASSEL-MLM, which were not evaluated in our study but implemented the same technique as GEMMA or GAPIT-MLM. The algorithms in the first group produced higher power for QTL with low MAF, explaining relatively large PG and for QTL pairs with medium to high LD. In contrast, the second group of algorithms was better at detecting independent QTL with medium to high MAF and explaining small PG. The relative behavior of the two groups of algorithms is very interesting because their underlying models differ greatly. While the first group followed the standard Q+K LMM (GEMMA, Grid-LMM and GAPIT-MLM) or introduced only minor modifications (CMLM), the second group employed techniques that differed greatly from the Q+K model. Our results in the simulation studies indicated that most regions identified by FarmCPU and BLINK, but not by GEMMA, in the empirical datasets were unlikely false positives since both groups of algorithms stringently controlled the FPR, but the model differences resulted in a complementary detection power. Based on all results, we recommend a combination of two algorithms, each from one group, as the optimal strategy for performing GWAS.

## Methods

### The procedure of a standard GWAS algorithm based on the Q+K model

In this section, we briefly describe the Q+K model ^7^ for GWAS and the procedure of solving the model as well as producing the test statistics. More details are provided in the Supplementary Note A.

For simplicity, the model is presented in the case that each individual has only one phenotypic observation as follows:

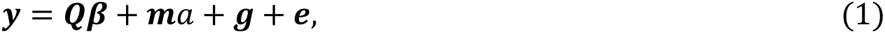

where ***y*** is the *n*-dimensional vector of phenotypic observations, ***β*** is the *k*-dimensional vector of covariates which may include a common intercept, environmental and/or subpopulation effects etc., ***Q*** is the corresponding design matrix of size *n* × *k*, *a* is the effect of the marker being tested, ***m*** is the *n*-dimensional coding vector of the marker, ***g*** is the *n*-dimensional vector of polygenic effects, and ***e*** is the *n*-dimensional vector of residuals.

In the model, ***β*** and *a* are treated as fixed effects, ***g*** and ***e*** are random effects following multi-variate normal distribution: 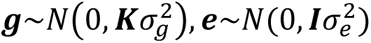, where ***K*** is an *n* × *n* kinship matrix derived from pedigree/marker information, ***I*** is the *n* × *n* identity matrix, 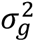 and 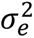 are the corresponding genetic and residual variance components.

The procedure of GWAS can be roughly divided into two steps: 1) solving the model; 2) producing the test statistics. Usually, the model is solved by maximum likelihood (ML) or restricted maximum likelihood (REML) method. Taking the ML method as an example, the log-likelihood function is the following:

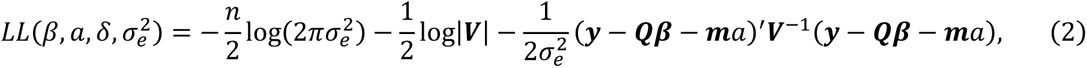

where 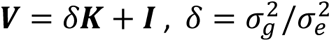, and | · | denotes the determinant of a matrix. The unknown parameters, namely *β*, *a*, *δ* and 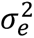, are estimated as the values such that the log-likelihood function reaches its maximum.

The test statistics can be produced with different approaches, e.g. the likelihood ratio test and the Wald test. Taking the Wald test as an example, the test statistic has the following form

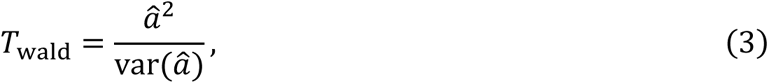

where *â* is the estimated value of *a*. Under the null hypothesis, the test statistic follows a *χ*^2^ - distribution with one-degree of freedom.

### The time complexity of an algorithm

For the convenience of readers, we briefly recall the time complexity of an algorithm ^44^. In computer science, the time complexity describes the amount of computer time it takes to run an algorithm. It is usually estimated by counting the number of elementary operations, i.e. additions and multiplications of numbers, performed by the algorithm. Assuming that each elementary operation takes a fixed amount of time, the amount of time taken and the number of elementary operations performed by the algorithm are related by a constant factor.

In most cases, the running time of an algorithm depends on the size of input data. Thus, the time complexity is generally expressed as a function of the size of the input. Since this function is generally difficult to compute exactly, one commonly focuses on the behavior of the complexity when the input size increases, i.e., the asymptotic behavior of the complexity. Therefore, the time complexity is commonly expressed by the so-called “big O” notation. For example, suppose that the size of input data depends on two variables *m* and *n*, an algorithm with time complexity *O*(*mn*^2^) means that the amount of running time increases linearly as the increase of *m*, and quadratically as the increase of *n*.

Here are useful results about the time complexity of some basic operations in matrix algebra: The time complexity of multiplying an *m* × *n* matrix with an *n* × *k* matrix is *O*(*mnk*). Thus, the product of two *n* × *n* matrices has complexity *O*(*n*^3^), and the complexity of multiplying an *m* × *n* matrix with an *n*-dimensional vector is *O*(*mn*). The inner product of two n-dimensional vectors has complexity *O*(*n*). The complexity of inverting or performing the spectral decomposition of an *n* × *n* matrix is *O*(*n*^3^).

### Fast GWAS algorithms evaluated in this study

The principles of selecting algorithms for benchmarking analysis were the following: 1) Among the same class of algorithms in which similar techniques for improving the computational efficiency were implemented. Only if an algorithm was reported in the literature to be clearly inferior to the others, it was excluded from the analysis. 2) In case several algorithms from different software packages implemented the same techniques, only one representative was selected. The results for the selected algorithm should then be treated as equally working for the entire class of algorithms that it represents. 3) As long as 1) is not violated, we tried to include as many different techniques as possible. Each technique is represented by at least one selected algorithm.

According to the above principles, the following decisions were made:

1. Among the four exact and quasi-exact algorithms (EMMA, FaST-LMM, GEMMA and Grid-LMM), we selected GEMMA and Grid-LMM for our analysis. EMMA is computationally inefficient (complexity *O*(*pn*^3^ + *tpn*)) compared with GEMMA and FaST-LMM (complexity *O*(*n*^3^ + *pn*^2^ + *tpn*)). In fact, the R package EMMA has been removed from the CRAN repository. The new version of FaST-LMM (based on Python) implements P3D and the exact FaST-LMM algorithm is available only in the old C++ version. Thus, we decided to take GEMMA as the representative. Grid-LMM was selected because it solves the LMM by grid search instead of numerical optimization, hence it is different from the other three algorithms.
2. There are several algorithms implementing P3D with the standard Q+K model (but without other techniques), namely GAPIT-MLM, FaST-LMM-P3D, EMMAX, and TASSEL-MLM. There is no essential difference among these algorithms and we selected GAPIT-MLM as the representative.
3. A few algorithms implementing the LOCO technique, including FaST-LMM (as an option), MLMA-LOCO and BOLT-LMM. Since P3D is mandatorily implemented in the new version of FaST-LMM and in MLMA-LOCO, it means that both algorithms implemented P3D and LOCO based on the standard Q+K model, and without other techniques. We selected MLMA-LOCO as a representative. BOLT-LMM was selected as it implements the Monte Carlo sampling approach to solve the LMM which is different from all other algorithms. There were two options in BOLT-LMM for controlling the genetic background effects or the population structure. One follows the standard Q+K model, termed BOLT-LMM-inf. The other implements a Gaussian mixture for the random marker effects, similar to a Bayesian genomic prediction model ^45^, termed BOLT-LMM-mix. The default BOLT-LMM algorithm combined the two variants and performed a cross-validation to determine which variant would be used to produce the final test statistics. In our study, we purposely treated BOLT-LMM-inf and BOLT-LMM-mix as two different algorithms to assess the influence of the different techniques implemented in the two variants.
4. Several algorithms implemented approximations to the test statistics, namely GRAMMAR, GRAMMAR-Gamma, BOLT-LMM, and fastGWA. BOLT-LMM was already selected, while for the remaining three we only included fastGWA. According to the previous studies, the algorithm GRAMMAR produces conservative tests and biased estimates ^19, 20^ and it was improved in GRAMMAR-Gamma, which was also implemented as an option in fastGWA, termed fastGWA-GG. Since the package GenABEL implementing GRAMMAR and GRAMMAR-Gamma has been removed from the CRAN repository, both were excluded in our analysis. But fastGWA-GG was selected to represent GRAMMAR-Gamma. Besides, fastGWA implemented another option of making the kinship matrix sparse, termed fastGWA-sp. This variant was also selected for our analysis.
5. Among the two algorithms which compress the kinship matrix (CMLM and ECMLM), we selected CMLM because the enriched version ECMLM is computationally much more demanding for large data sets despite it may increase the power of detection ^18^.
6. For the algorithms that select a subset of markers to control the population structure or polygenic background (MLMM, FaST-LMM-select, FaST-LMM-all+select, SUPER, FarmCPU and BLINK), we selected FaST-LMM-select, FarmCPU and BLINK because MLMM, FaST-LMM-all+select and SUPER were much more time-demanding than the others when data size is large. The three selected algorithms differ in the method for selecting markers and/or in the testing procedure (For details see Supplementary Note A).

To summarize, 12 algorithms were selected for our benchmarking analysis: GEMMA, Grid-LMM, GAPIT-MLM, MLMA-LOCO, BOLT-LMM-inf, BOLT-LMM-mix, fastGWA-GG, fastGWA-sp, CMLM, FaST-LMM-select, FarmCPU, and BLINK.

After we had started the benchmarking analysis, two interesting new algorithms MM4LMM ^16^ and REGENIE ^29^ were published. MM4LMM is an exact algorithm which solves the LMM in a different way from GEMMA/FaST-LMM. REGENIE implements P3D, LOCO and a two-step stacked ridge regression approach to solve the null model. We evaluated the two algorithms with 400 simulated data sets (4 out of the 108 scenarios) and the results were summarized in Supplementary Note C.

### Protocols and settings of the algorithms evaluated in this study

The GEMMA package (v0.98.1) was downloaded at https://github.com/genetics-statistics/GEMMA. All parameters were set as default.

The Grid-LMM package (v0.0.0.9000) was downloaded at https://github.com/deruncie/GridLMM. All parameters were set as default.

The BOLT-LMM package (v2.3.5) was downloaded at http://data.broadinstitute.org/alkesgroup/BOLT-LMM/downloads/. The parameters “--ImmInfOnly” and “--ImmForceNonInf” were used to force the algorithm producing the test statistics of BOLT-LMM-inf and BOLT-LMM-mix, respectively. Other parameters were set as default. Note that by default, the two algorithms implemented LOCO when calculating the test statistics, but estimated variance components only once using all markers. We kept this setting for our benchmarking analysis, but investigated the influence of the genuine LOCO using 400 simulated data sets. To force a genuine LOCO procedure, we modified the parameter “--modelSnps” to select markers on all chromosomes except the one to which the marker being tested belonged.

The GAPIT package (v3.1.0) was downloaded at https://zzlab.net/GAPIT/. Four algorithms implemented in this package were evaluated in our study, namely MLM, CMLM, FarmCPU and BLINK. All parameters were set as default. Note that for CMLM, the default setting is to optimize the compression level by evaluating the null model with a series of different compression levels and choosing the one maximizing the log-likelihood. We also tested alternative settings, namely fixing the compression level to 5 or 10 as suggested, with 200 simulated data sets. Although it could greatly improve the computational efficiency, inflated FPR were observed (Supplementary Table 3). Thus, we decided to keep the default setting.

The GCTA package (v1.93.2beta) was downloaded at https://yanglab.westlake.edu.cn/software/gcta/#Overview. Three algorithms implemented in this package were evaluated in this study: MLMA-LOCO, fastGWA-GG and fastGWA-sp. The parameter --mlma-loco” was used for MLMA-LOCO and “--fastGWA-mlm” was used for fastGWA-GG and fastGWA-sp. For fastGWA-sp, an additional parameter “--fastGWA-mlm-exact” was set to exclude the GRAMMA-Gamma approximation and the sparse kinship was done by setting “--make-bK-sparse” to the recommended value 0.05, except for the analysis related to Supplementary Fig. 15, for which five different thresholds were used. Other parameters were set as default.

FaST-LMM-select was implemented in the package FaST-LMM (Python platform, v0.6.1), which was downloaded at https://pypi.org/project/fastlmm/. The function “fastlmm.association.single_snp_select()” was used to run FaST-LMM-select. All parameters were set as default.

### Empirical data sets

#### Arabidopsis

The Arabidopsis data set was from the 1001 Genomes Consortium ^46^, which is comprised of 1,134 genotypes and 11,458,975 SNPs. The flowering time recorded for plants grown at a temperature of 10℃ (abbreviated as FT10) was selected as the phenotypic data in this study. After filtering with missing rate (≤ 0.1) and MAF (≥ 0.05), 1,003 genotypes with 749,722 SNPs were used for the current study. The remaining missing values were imputed using IMPUTE2 ^47^.

#### Wheat

The wheat data set consisted of 5,581 winter wheat accessions from the Federal ex situ Genebank for Agricultural and Horticultural Crop Species of Germany hosted at the Leibniz Institute of Plant Genetics and Crop Plant Research (IPK) ^48^. The accessions were fingerprinted using genotyping-by-sequencing (GBS). After quality control and filtering, IMPUTE2 was applied to impute the remaining missing sites, resulting in 427,937 SNP markers. The phenotypic trait considered in this study was yellow rust (YR) resistance based on natural infections in replicated field experiments over years 2015-2020 at two locations in Germany ^48^.

#### Rice

The rice data set was from the 3,000 Rice Genomes Project (https://snpseek.irri.org/_download.zul) ^49^. The genotypic data of 3,024 genotypes were filtered with missing rate (< 0.2) and MAF (> 0.01), resulting in 4,817,964 bi-allelic SNPs. Then, the remaining missing sites were imputed by Beagle 5.2 ^50^. In total, 2,013 genotypes with the phenotypic trait grain length were used for the analysis (https://www.rmbreeding.cn/phenotype#ifr2).

#### Maize

The maize data set consisted of 2,815 inbred accessions preserved in the USDA collection ^51^. The growing degree days to silking were investigated in three environments (Ames, IA; Clayton, NC; and Aurora, NY) during summer 2010. The accessions were genotyping by GBS with 681,257 SNP markers. Both phenotypic and genotypic data were obtained from Panzea database (https://www.panzea.org). After filtering the missing phenotypic data, 2,279 accessions remained. The genotypic data were filtered with MAF > 0.05 and imputed with Beagle. In total, 225,563 high-quality SNP markers were used for the current study.

#### Barley

The barley data set consisted of 15,557 spring barley accessions fingerprinted by GBS, which were also from the Genebank at IPK ^52^. After quality control, the missing values were imputed using FILLIN ^53^ resulting in 306,049 SNPs. The phenotypic information which was considered in this study were historical data for flowering time (FT, 8,825 accessions) ^54^.

### GWAS for the empirical data sets

Each of the 12 selected GWAS algorithms was applied to the five empirical data sets described above. The *p*-values of all markers were obtained and *p* < 0.05 after Bonferroni correction ^39^ for multiple testing was determined as the genome-wide threshold for significance. The significant markers were merged into QTL by the following criterion: Two markers were merged if the physical distance between them is less than the average distance at which the LD (measured by *r*^2^) decayed to 0.1 (for wheat, barley and rice) or 0.05 (for Arabidopsis and maize), which is determined by non-linear regression ^55^. In the five populations, the resulting distance was 380 kbp (wheat), 452 kbp (rice), 1,400 kbp (barley), 26 kbp (Arabidopsis) and 70 kbp (maize), respectively.

### Candidate gene search for marker-trait associations in Arabidopsis

Genes spanning or flanking significant SNPs were retrieved from TAIR (https://www.arabidopsis.org/index.jsp). FLOR-ID ^56^ (http://www.phytosystems.ulg.ac.be/florid/) was inspected to identify genes for which a role in flowering time control had been documented previously. Regions 50 - 60 kbp upstream and downstream of the SNP were considered. In regions for which no candidate genes had been reported in FLOR-ID a literature search for all genes mapping to these regions was conducted based on the information available in TAIR. Only genes in which mutants and/or overexpressing lines of the genes of interest had shown an effect on flowering time were considered as candidate genes.

### Data simulation

The genomic data of the wheat population described in the last subsection were used to simulate the phenotypic data. We considered three different population sizes (300, 1000, 3000), three different trait heritabilities (0.3, 0.5, 0.7), and 12 different complexities of genetic architecture. Each of the 3 × 3 × 12 = 108 scenarios was repeatedly simulated 100 times, which makes in total 10,800 data sets. To reduce the computational load, we chose 6 chromosomes (1A to 6A) to conduct the simulation, resulting in 126,819 SNPs. In all cases, markers were classified into three classes, namely major QTL, minor QTL and neutral marker. The number of major QTL was fixed to 6, while the number of minor QTL and neutral markers varied across scenarios.

The complexity of simulated genetic architecture is determined as follows: First, we considered three different patterns of linkage disequilibrium (LD) between the major and minor QTL: In LD pattern 1, there was neither LD between any two major QTL nor between any major and minor QTL. In LD pattern 2, the major QTL were still independent of each other, but LD exited between major and minor QTL. In LD pattern 3, there existed LD among the major QTL as well as between major and minor QTL. Then, two cases of the proportion of genetic variance (PG) explained by the major QTL were considered (PG1 and PG2). In PG1, all 6 major QTL contributed equally, each explaining 2% of the genetic variance. Thus, the total PG of all major QTL was 12%. In PG2, the proportions of explained genetic variance of the 6 major QTL were randomly assigned as 2%, 4%, 6%, 8%, 10%, and 12%, with a total PG of 42%. Finally, two cases for the number of minor QTL contributed as genetic background effects were introduced (denoted by GB1 and GB2). With GB1, only a few markers were selected as minor QTL. With GB2, all markers on the chromosomes (LD pattern 2 and 3) or on half of the chromosomes (LD pattern 1) contributed as minor QTL, representing the so-called infinitesimal genetic architecture. In total, it gives 3 × 2 × 2 = 12 different levels of complexities.

Next, we described the detailed procedure of simulation. For LD pattern 1, in each round of simulation, three of the six chromosomes were randomly sampled. On each chromosome, two markers with very low LD (*r*^2^ < 0.01) were randomly selected as major QTL. Minor QTL came from the remaining three chromosomes. Namely, 400 markers from each of the three chromosomes were randomly sampled in GB1, while in GB2 all markers on the three chromosomes were treated as minor QTL. For LD pattern 2, one marker was randomly sampled from each of the six chromosomes as major QTL. For the minor QTL, 200 markers were randomly sampled from each chromosome in GB 1, and all remaining markers were treated as minor QTL in GB 2. For LD pattern 3, we only took 3 chromosomes (A1-A3) to conduct the simulation. In each round of simulation, three different levels of LD, namely 0.16 < *r*^2^ ≤ 0.36, 0.36 < *r*^2^ ≤ 0.64, 0.64 < *r*^2^ < 1, were randomly assigned to the three chromosomes as the criterion for sampling major QTL. Then, two markers fulfilling the LD criterion were randomly selected from each of the three chromosomes. In addition, we purposely forced the distance between the two markers sampled as major QTL to be larger than 760 kbps, which is the double distance at which the LD decayed to 0.1. This setting is to make sure that the two QTL would not be treated as a single one in the assessment of statistical power (see the next subsection). For the minor QTL, 400 markers were randomly sampled from each chromosome in GB1, and all remaining markers were treated as minor QTL in GB2.

In LD patterns 1 and 2, we additionally controlled the MAF of the markers sampled as major QTL. Three classes of MAF were considered, namely MAF ≤ 0.1, 0.1 < MAF ≤ 0.3, MAF > 0.3. On each chromosome, the number of markers sampled as major QTL across 100 replicates in each class was about 1/3 of the total number. In LD pattern 3, we did not control MAF because setting many criteria may violate the randomness of the sampling procedure, considering that there was already a control of LD between the pair of markers sampled as major QTL.

The simulated phenotypic data were produced by the following formula:

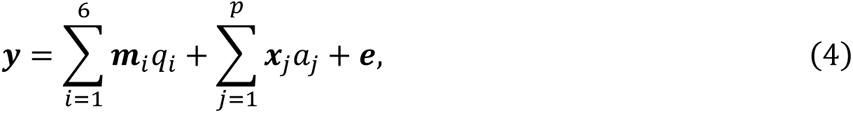

where ***y*** is the vector of simulated phenotypes, *q*_*i*_ is the effect of the *i*-th major QTL, ***m***_*i*_ is the corresponding marker coding vector, *p* is the number of minor QTL, *a*_*j*_ is the effect of the *j*-th minor QTL, ***x***_*j*_ is the corresponding marker coding vector, ***e*** is the vector of residuals.

More precisely, in each round of simulation, the vector ***y*** was produced by the following five steps: 1) For a given population size *N* (300, 1,000 or 3,000), we randomly sampled *N* Genotypes from the entire population and extracted the SNP matrix. Then, we filtered out SNPs whose MAF was below 0.05 (if *N*=300) or below 0.01 (if *N* = 1,000 or 3,000). Subsequently, the filtered SNP matrix was used to generate the simulated phenotype. 2) The effect *a*_*j*_ (for any *j*) was randomly sampled from a normal distribution *N*(0,0.5). Then, we summed up the effects of all minor QTL as 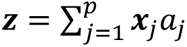 The variance var(***z***) was calculated. For a given case of PG for the major QTL (PG 1 or PG 2), the variance of the contribution of each major QTL ***u***_***i***_ = ***m***_*i*_*q*_*i*_ must satisfy the equation *P*_*i*_⁄(1 − *P*) = var(***u***_***i***_) ⁄var(***z***), where *P*_*i*_ is the proportion of genetic variance explained by the *i*-th major QTL, *P* is the proportion of genetic variance explained by all major QTL. Then, var(***u***_***i***_) = *P*var(***z***)⁄(1 − *P*). Now, the effect of the *i*-th major QTL can be calculated as 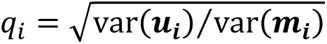. 4) We calculated the total genetic effects as 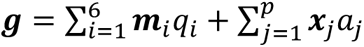. Then, for a given heritability *h*^2^ (0.3, 0.5 or 0.7), the variance of the residuals must satisfy the equation *h*^2^⁄(1 − *h*^2^) = var(***g***) ⁄var(***e***). Thus, var(***e***) = (1 − *h*^2^)var(***g***)⁄*h*^2^. Then, each entry of the residual vector was randomly sampled from a normal distribution 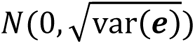. 5) The simulated phenotypic value was generated by Eq. (4).

### Assessing the statistical power and false positive rate

Each of the 12 selected GWAS algorithms was applied to all 10,800 simulated data sets. The *p*-values of all markers were obtained and *p* < 0.05 after Bonferroni correction ^39^ for multiple testing was determined as the genome-wide threshold for significance.

In each of the 108 scenarios, the statistical power and FPR of an algorithm was assessed through its performance across the 100 replicated data sets. More precisely, the power of detecting major QTL was calculated as the number of correctly detected ones divided by the total number of simulated major QTL across 100 data sets, which is 6 × 100 = 600. When the physical distance between a major QTL and a significant marker was within 380 kbp, the QTL was considered as correctly detected. Note that the interval length 380 kbp was the average distance at which the LD decayed to 0.1. This is the same as the criterion of merging significant markers into QTL in the empirical data sets. We did not assess the power of detecting minor QTL because they were considered as contributors to the polygenic background. The FPR was estimated only for scenarios with GB1 (1,200 markers as minor QTL), because in GB2 all markers were either major or minor QTL. The FPR was calculated as the ratio between the number of non-QTL markers wrongly detected as significant and the total number of non-QTL markers, averaged across 100 replicates. A marker was defined as a non-QTL marker if it was not within an interval of 380 kbp flanking a major QTL and nor was it a minor QTL.

In addition, we also divided the simulated major QTL into different classes and investigated the detection power in each specific class. 1) QTL with different MAF. We considered three classes: MAF ≤ 0.1, 0.1 < MAF ≤ 0.3, and MAF > 0.3. Note that in LD patterns 1 and 2, we purposely controlled the MAF of the simulated major QTL such that the proportion in each class was about 1/3. But in LD pattern 3, the MAF of simulated major QTL was random. Nevertheless, this analysis was performed for all data sets. 2) QTL with different proportions of explained genetic variance. This was only for scenarios with PG2 in which six different PGs from 2% to 12 % were assigned to the simulated major QTL. 3) QTL pairs with different LD. This was only applied to LD pattern 3 in which three levels of LD were assigned to the QTL pairs: 0.16 < *r*^2^ ≤ 0.36, 0.36 < *r*^2^ ≤ 0.64, and 0.64 < *r*^2^ < 1.

### Complementary simulation studies

Additional data were simulated based on the genomic data of the Arabidopsis, maize and barley populations. The rice data set was excluded due to the computational load of GWAS later on. We chose four scenarios resulting from the combination of two LD patterns (LD pattern 2 and 3) and two cases of PG (PG1 and PG2). The other parameters were fixed as follows: population size 1,000, trait heritability 0.7 and GB1. For each scenario and with each population, 100 simulated data sets were generated by the same procedure as simulating data sets based on the wheat population. In total, there were 1,200 additional simulated data sets.

Note that in LD pattern 3, the minimal distance between the two simulated QTL on the same chromosome was set to be the double distance at which the LD decayed to 0.1. Thus, this value depended on the population, which was 16.6 kbp (Arabidopsis), 6 kbp (maize) and 2,800 kbp (barley) respectively.

The 12 selected GWAS algorithms were applied to the 1,200 simulated data sets. For each species and in each scenario, the statistical power and FPR were calculated by the same approach as described in the previous subsection. Again, the interval flaking a QTL which was used to determine whether a significant marker is true or false positive depended on the population, namely 8.3 kbp (Arabidopsis), 3 kbp (maize), and 1,400 kbp (barley), respectively.

## Acknowledgements

The work was partly supported by the German Federal Ministry of Education and Research within the GeneBank3.0 Project (grant number: FKZ031B1300A).

## Data availability

The five empirical data sets used in this study were publicly available. All simulated data sets and the results (*p*-values for all markers) of 12 GWAS algorithms on the simulated/empirical data sets were provided in https://osf.io/keam8/.

## Code availability

All computer codes that were used to run the GWAS algorithms in this study were provided in https://osf.io/keam8/.

---

**Extended Data Figure 1.**
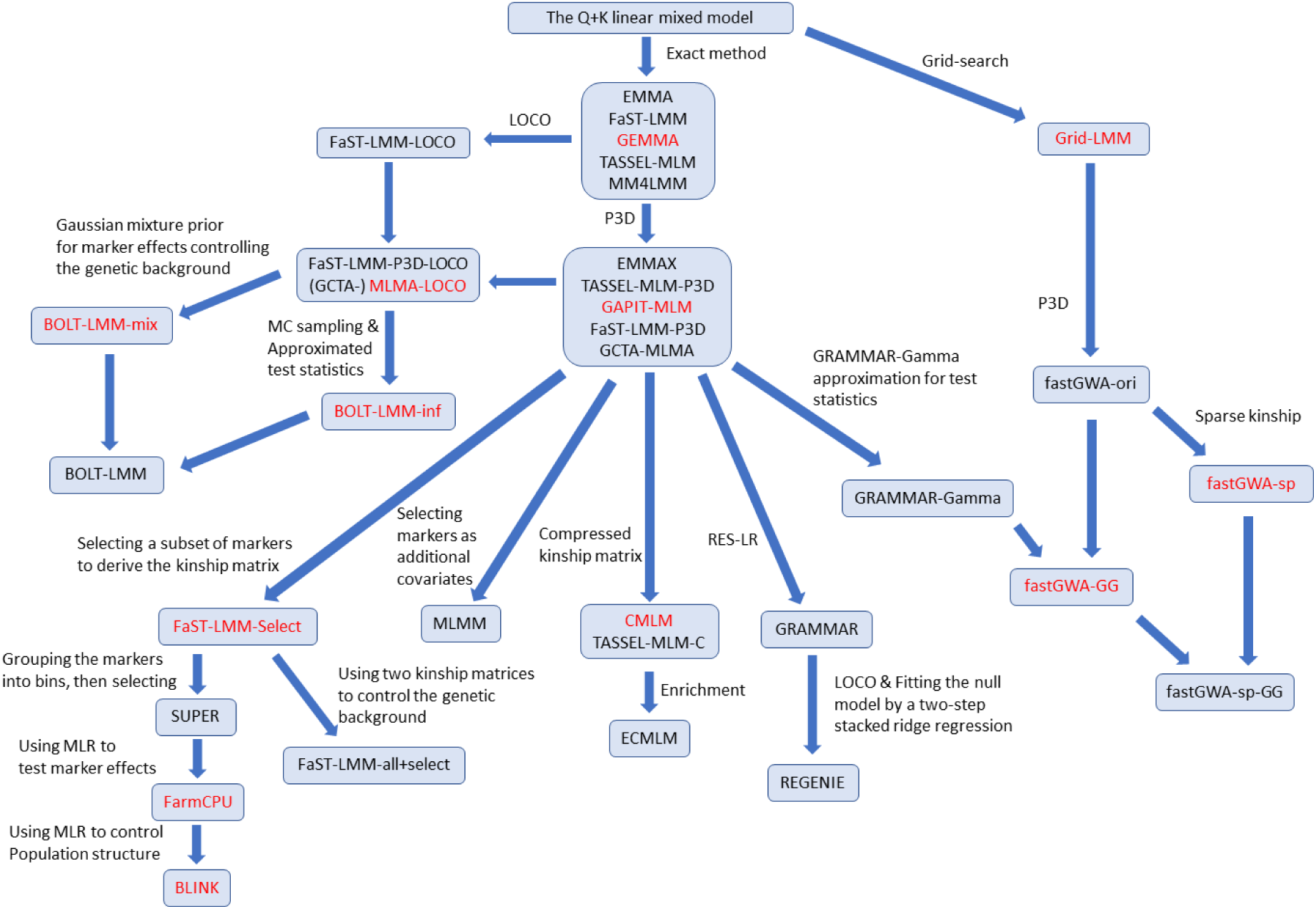
A phylogeny of 33 GWAS algorithms. The 12 algorithms evaluated in the benchmarking analysis are shown in red font. If two algorithms are connected by an arrow, it means that the target is based on the source with additional techniques indicated by the text next to the arrow. If two algorithms each with an arrow targeting to the same algorithm, it means that the target combines the techniques implemented by the two sources (In this case, no text was indicated). P3D, population parameters previously determined; MC, Monte-Carlo; LOCO, leave-one-chromosome-out; MLR, multi-variate linear regression; RES-LR, using the residuals from the null model as the response to test marker effects in a simple linear model.

**Extended Data Figure 2.**
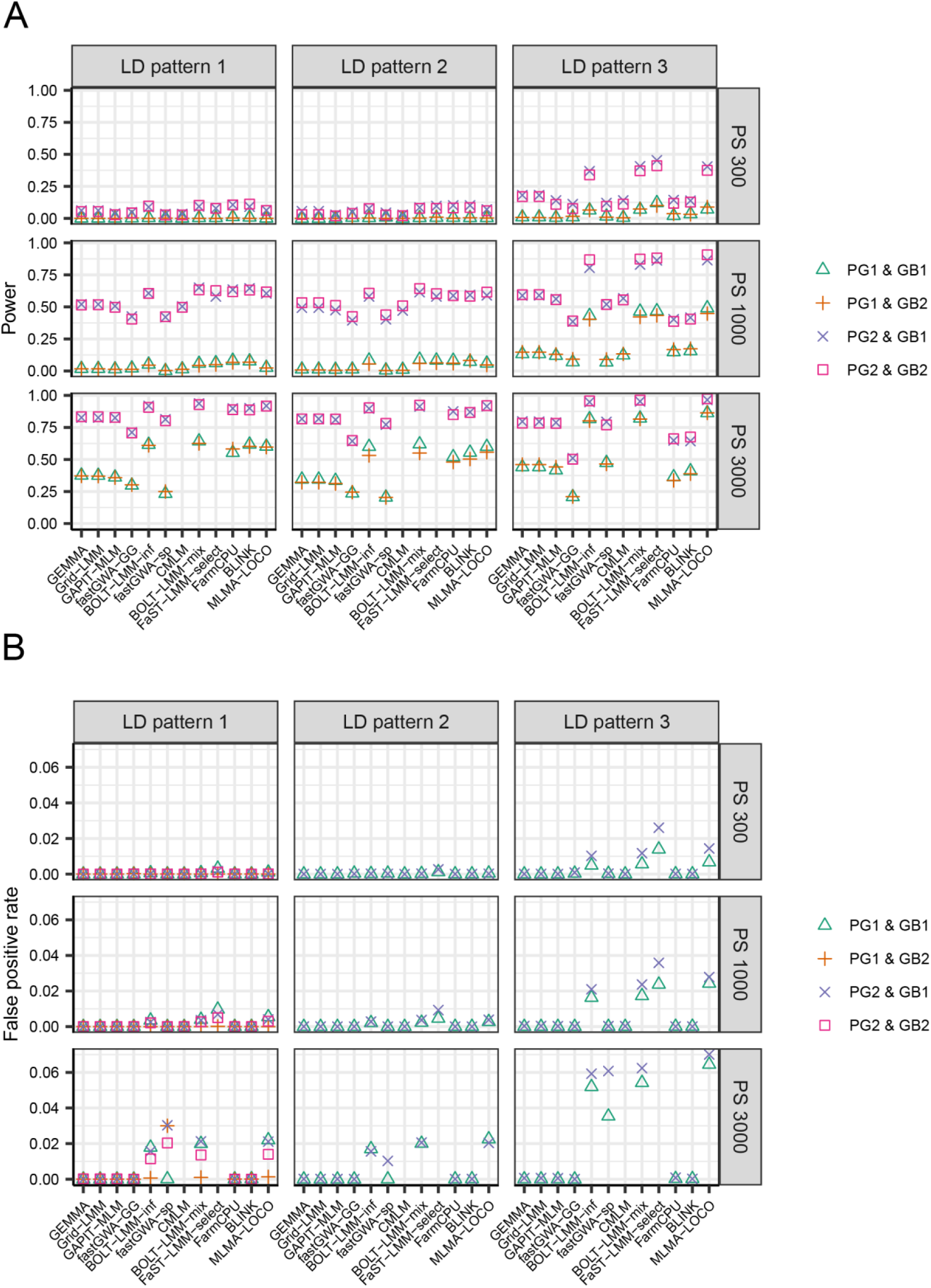
The statistical power (**A**) and false positive rate (**B**) of 12 GWAS algorithms evaluated in simulated data sets with 36 scenarios for trait heritability 0.5, under the threshold of *p* < 0.05 after Bonferroni correction for multiple testing. The 36 scenarios are combinations of three population sizes (PS 300, PS 1000 and PS 3000), three different linkage disequilibrium (LD) patterns among the QTL (LD patterns 1-3), two patterns of QTL effect sizes (PG1 and PG2), and two different genetic backgrounds (GB1 and GB2). In LD pattern 1, there is no LD between any two major QTL or between a major and a minor QTL. In LD pattern 2, there is no LD between any two major QTL, but LD exits between major and minor QTL. In LD pattern 3, there exists LD among the major QTL as well as between major and minor QTL. In PG1, each of the 6 major QTL explained 2% of the genetic variance. In PG2, the 6 major QTL were randomly assigned to explain 2%, 4%, 6%, 8%, 10% and 12% of the genetic variance respectively. In GB1, there were 1,200 markers as minor QTL. In GB2, all markers on the chromosomes (LD patterns 2 and 3) or on half of the chromosomes (LD pattern 1) contributed as minor QTL. Each of the 9 subpanels showed the results of a specific combination of population size and LD pattern. Within each subpanel, the results of four combinations of two PGs and two GBs were indicated by different symbols. For GB2, the FPR was only calculated for LD pattern 1, because for LD patterns 2 and 3, all markers contributed to the simulated trait either as major or as minor QTL. The algorithms CMLM and FaST-LMM-select were not evaluated for PS 3000 because the computational load was too high.

**Extended Data Figure 3.**
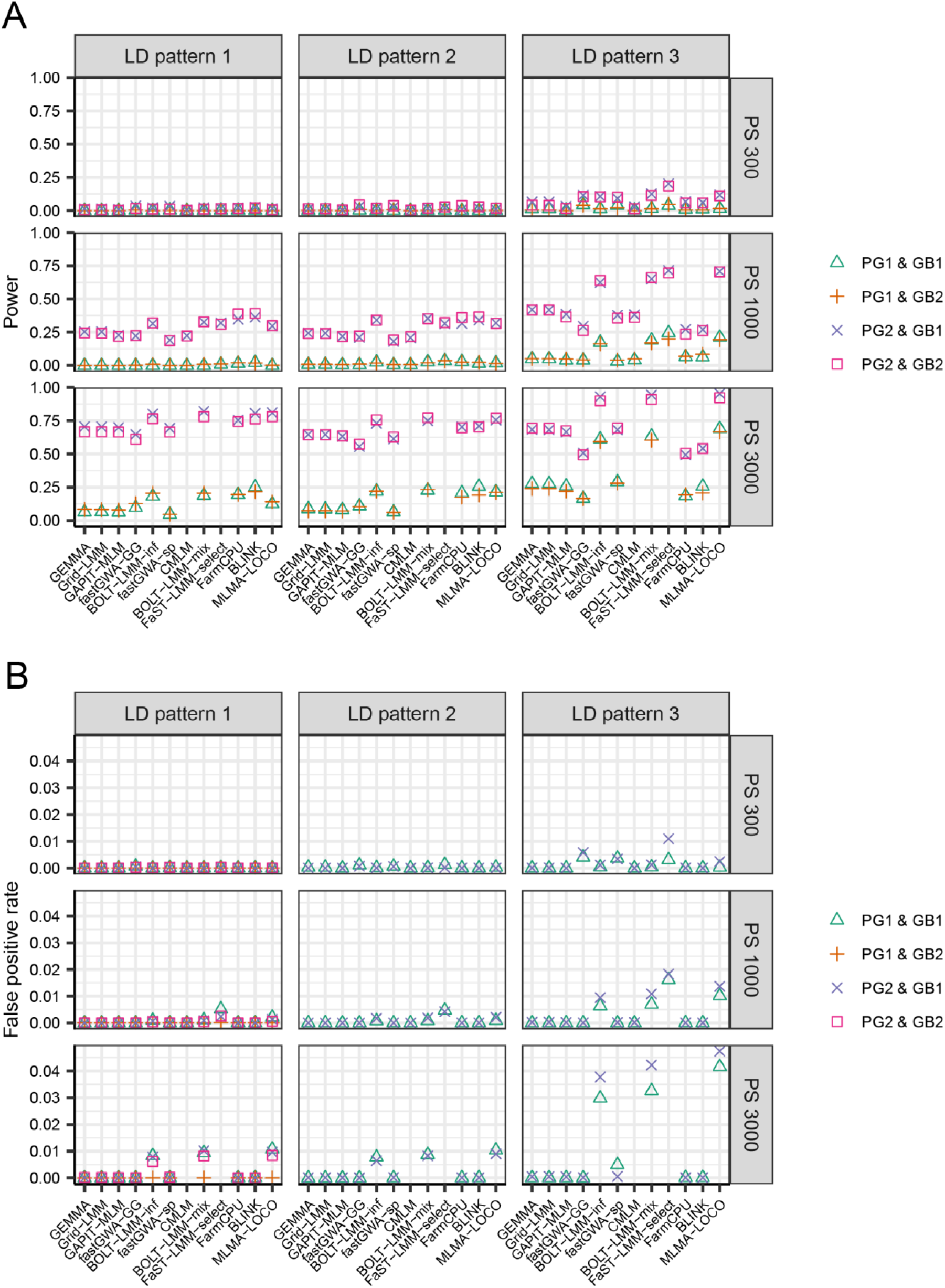
The statistical power (**A**) and false positive rate (**B**) of 12 GWAS algorithms evaluated in simulated data sets with 36 scenarios for trait heritability 0.3, under the threshold of *p* < 0.05 after Bonferroni correction for multiple testing. The 36 scenarios are combinations of three population sizes (PS 300, PS 1000 and PS 3000), three different linkage disequilibrium (LD) patterns among the QTL (LD patterns 1-3), two patterns of QTL effect sizes (PG1 and PG2), and two different genetic backgrounds (GB1 and GB2). In LD pattern 1, there is no LD between any two major QTL or between a major and a minor QTL. In LD pattern 2, there is no LD between any two major QTL, but LD exits between major and minor QTL. In LD pattern 3, there exists LD among the major QTL as well as between major and minor QTL. In PG1, each of the 6 major QTL explained 2% of the genetic variance. In PG2, the 6 major QTL were randomly assigned to explain 2%, 4%, 6%, 8%, 10% and 12% of the genetic variance respectively. In GB1, there were 1,200 markers as minor QTL. In GB2, all markers on the chromosomes (LD patterns 2 and 3) or on half of the chromosomes (LD pattern 1) contributed as minor QTL. Each of the 9 subpanels showed the results of a specific combination of population size and LD pattern. Within each subpanel, the results of four combinations of two PGs and two GBs were indicated by different symbols. For GB2, the FPR was only calculated for LD pattern 1, because for LD patterns 2 and 3, all markers contributed to the simulated trait either as major or as minor QTL. The algorithms CMLM and FaST-LMM-select were not evaluated for PS 3000 because the computational load was too high.

**Extended Data Figure 4.**
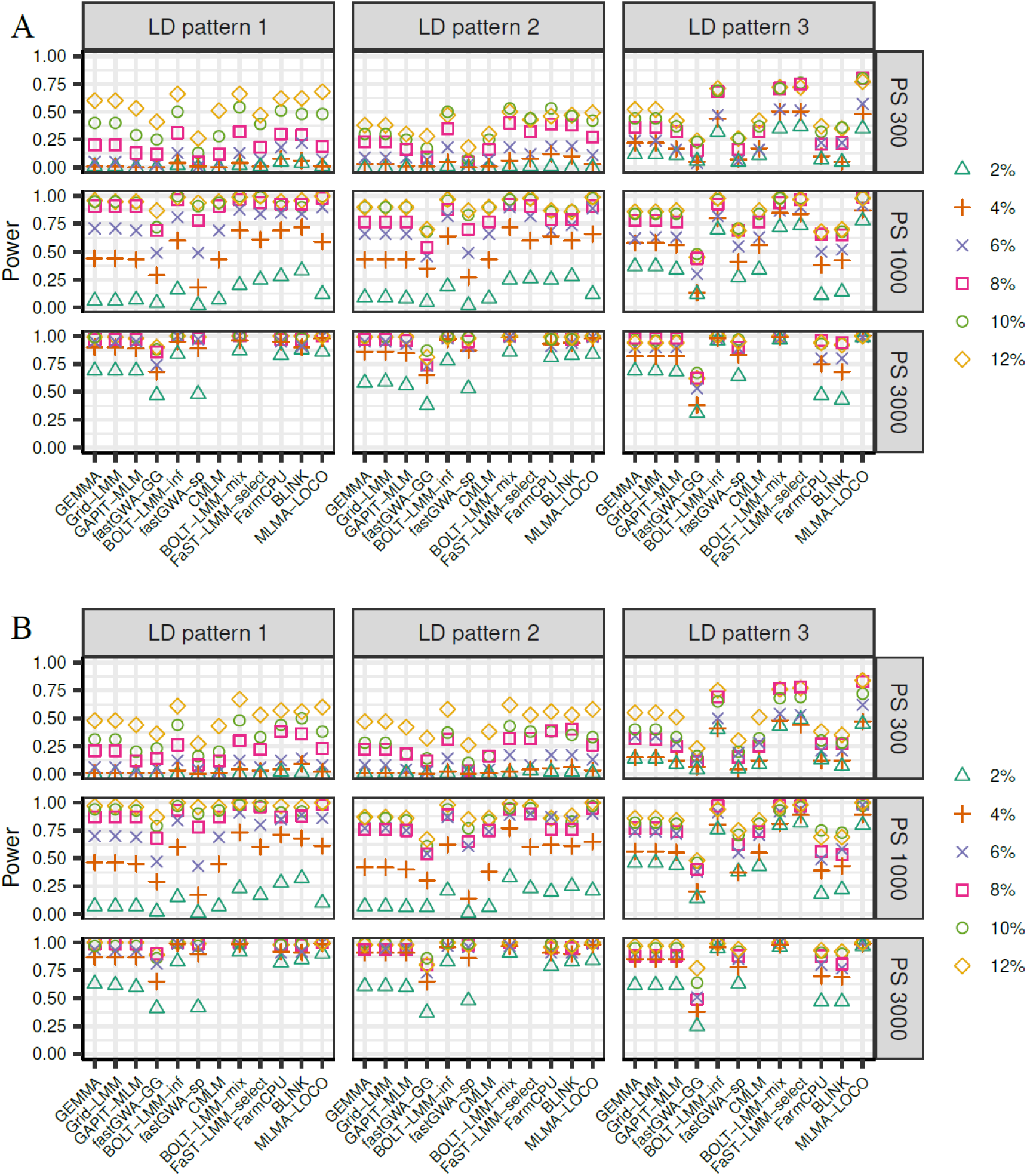
The statistical power of detecting QTL explaining a specific proportion of genetic variance for 12 GWAS algorithms evaluated in simulated data sets with 18 scenarios for trait heritability 0.7. The 18 scenarios are combinations of three population sizes (PS 300, PS 1000 and PS 3000), three different LD patterns among the QTL (LD patterns 1-3), and two different genetic backgrounds (GB1 and GB2). In LD pattern 1, there is no LD between any two major QTL or between a major and a minor QTL. In LD pattern 2, there is no LD between any two major QTL, but LD exits between major and minor QTL. In LD pattern 3, there exists LD among the major QTL as well as between major and minor QTL. In GB1, there were 1,200 markers as minor QTL. In GB2, all markers on the chromosomes (LD patterns 2 and 3) or on half of the chromosomes (LD pattern 1) contributed as minor QTL. The results for GB1 and GB2 were shown in panel **A** and **B**, respectively. Each panel was further divided into 9 subpanels, each showing the results of a specific combination of population size and data set. Within each subpanel, the results for QTL explaining six different PGs (from 2% to 12% with a step of 2%) were indicated by different symbols. The algorithms CMLM and FaST-LMM-select were not evaluated for PS 3000 because the computational load was too high.

**Extended Data Figure 5.**
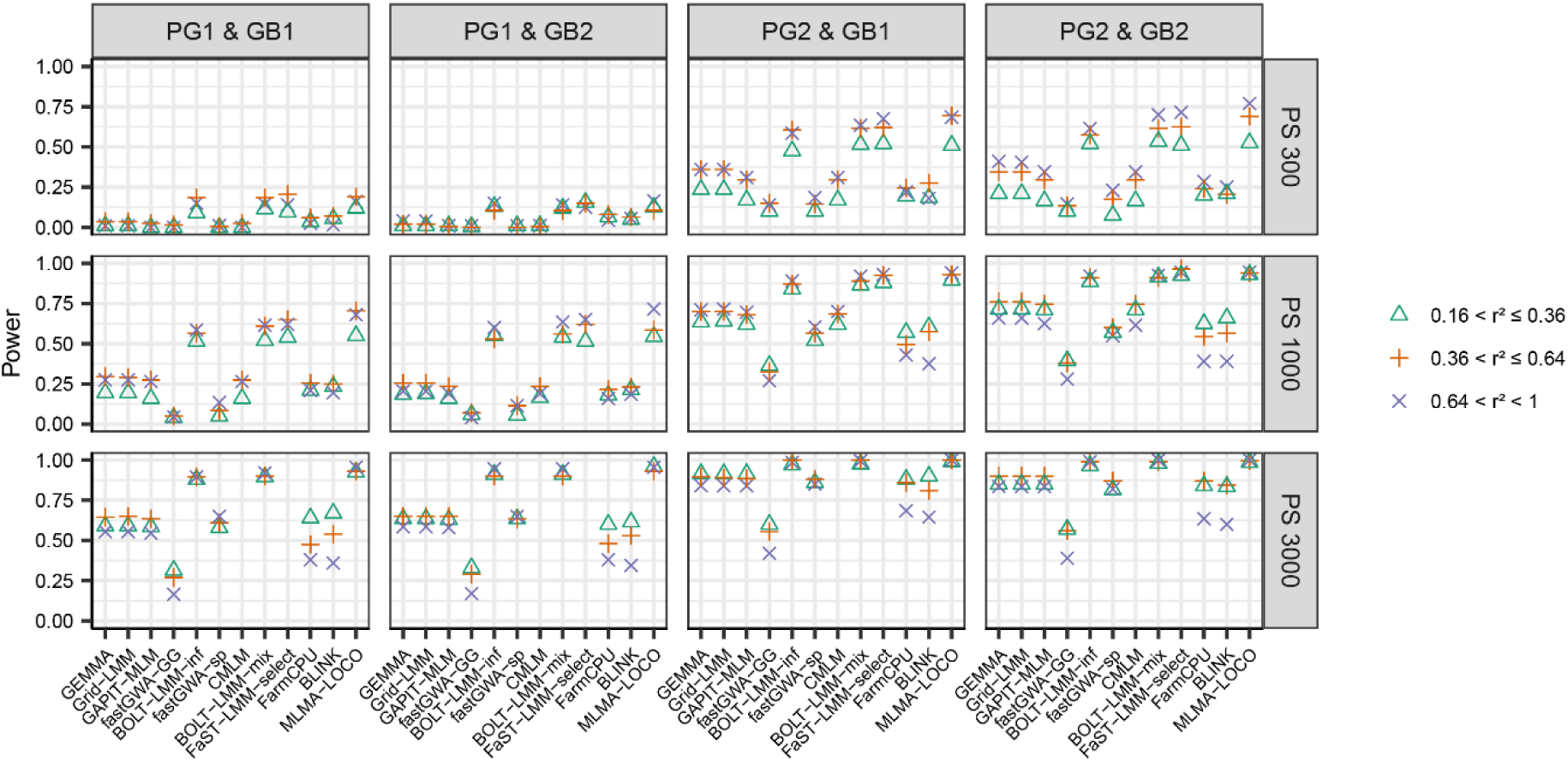
The statistical power of detecting QTL pairs with a particular range of linkage disequilibrium (LD) for 12 GWAS algorithms evaluated in simulated data sets with 12 scenarios for trait heritability 0.7. The 12 scenarios are combinations of three population sizes (PS 300, PS 1000 and PS 3000), two patterns of QTL effect sizes (PG1 and PG2), and two different genetic backgrounds (GB1 and GB2). In PG1, each of the 6 major QTL explained 2% of the genetic variance, In PG2, the 6 major QTL were randomly assigned to explain 2%, 4%, 6%, 8%, 10% and 12% of the genetic variance respectively. In GB1, there were 1,200 markers as minor QTL. In GB2, all markers on the chromosomes (LD patterns 2 and 3) or on half of the chromosomes (LD pattern 1) contributed as minor QTL. Each of the 12 subpanels showed the results of a specific combination of population size, PG and GB. Within each subpanel, the results for QTL pairs with three different ranges of LD (measured by *r*^2^) were indicated by different symbols. The algorithms CMLM and FaST-LMM-select were not evaluated for PS 3000 because the computational load was too high.

**Extended Data Figure 6.**
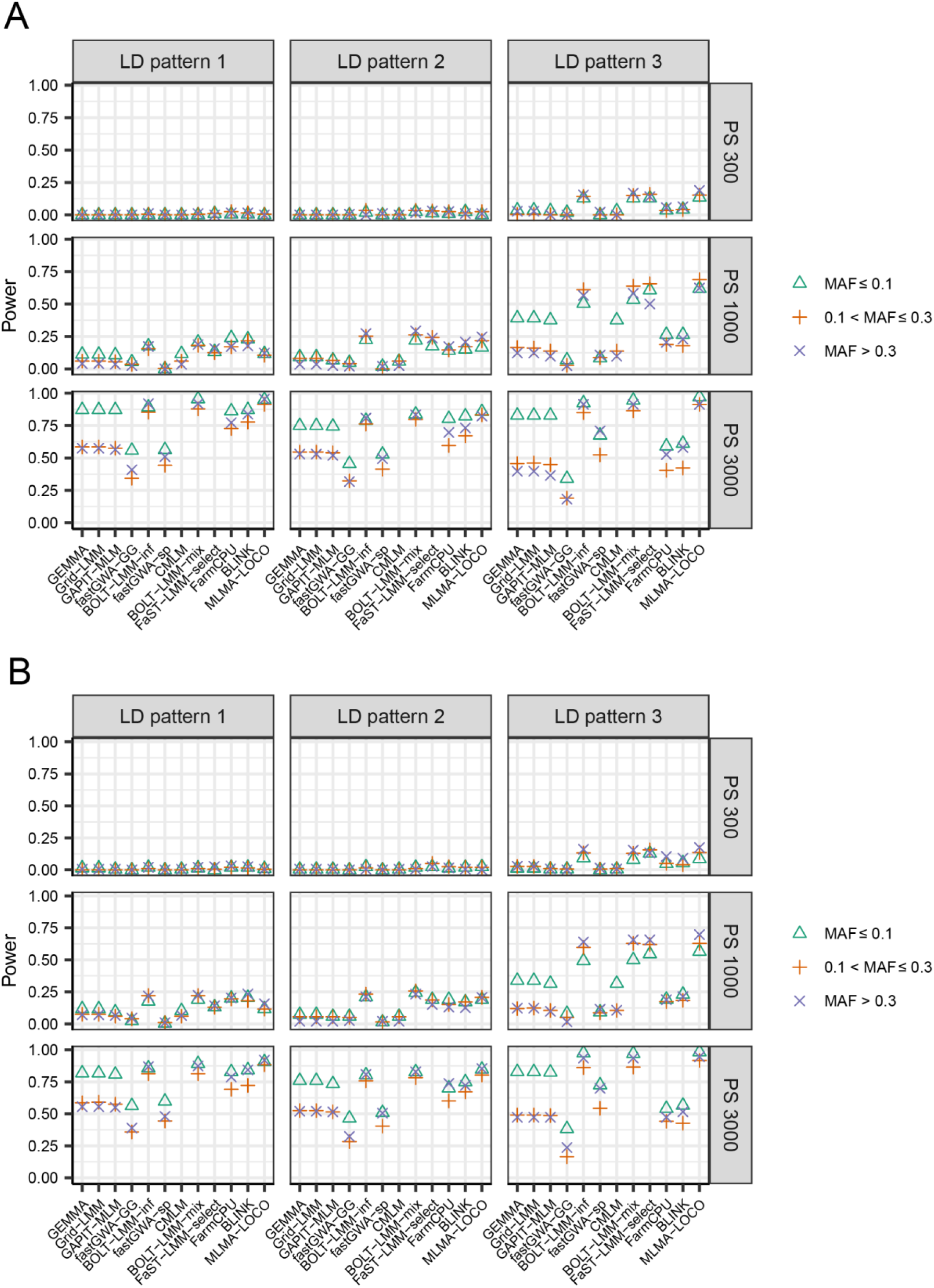
The statistical power of detecting QTL with a specific range of MAF for 12 GWAS algorithms evaluated in simulated data sets with 18 scenarios for trait heritability 0.7 with PG1 (each of the 6 major QTL explained 2% of the genetic variance). The 18 scenarios are combinations of three population sizes (PS 300, PS 1000 and PS 3000), three different LD patterns among the QTL (LD patterns 1-3), and two different genetic backgrounds (GB1 and GB2). In LD pattern 1, there is no LD between any two major QTL or between a major and a minor QTL. In LD pattern 2, there is no LD between any two major QTL, but LD exits between major and minor QTL. In LD pattern 3, there exists LD among the major QTL as well as between major and minor QTL. In GB1, there were 1,200 markers as minor QTL. In GB2, all markers on the chromosomes (LD patterns 2 and 3) or on half of the chromosomes (LD pattern 1) contributed as minor QTL. The results for GB1 and GB2 were shown in panel **A** and **B**, respectively. Each panel was further divided into 9 subpanels, each showing the results of a specific combination of population size and LD pattern. Within each subpanel, the results for QTL with three different ranges of MAF were indicated by different symbols. The algorithms CMLM and FaST-LMM-select were not evaluated for PS 3000 because the computational load was too high.

**Extended Data Figure 7.**
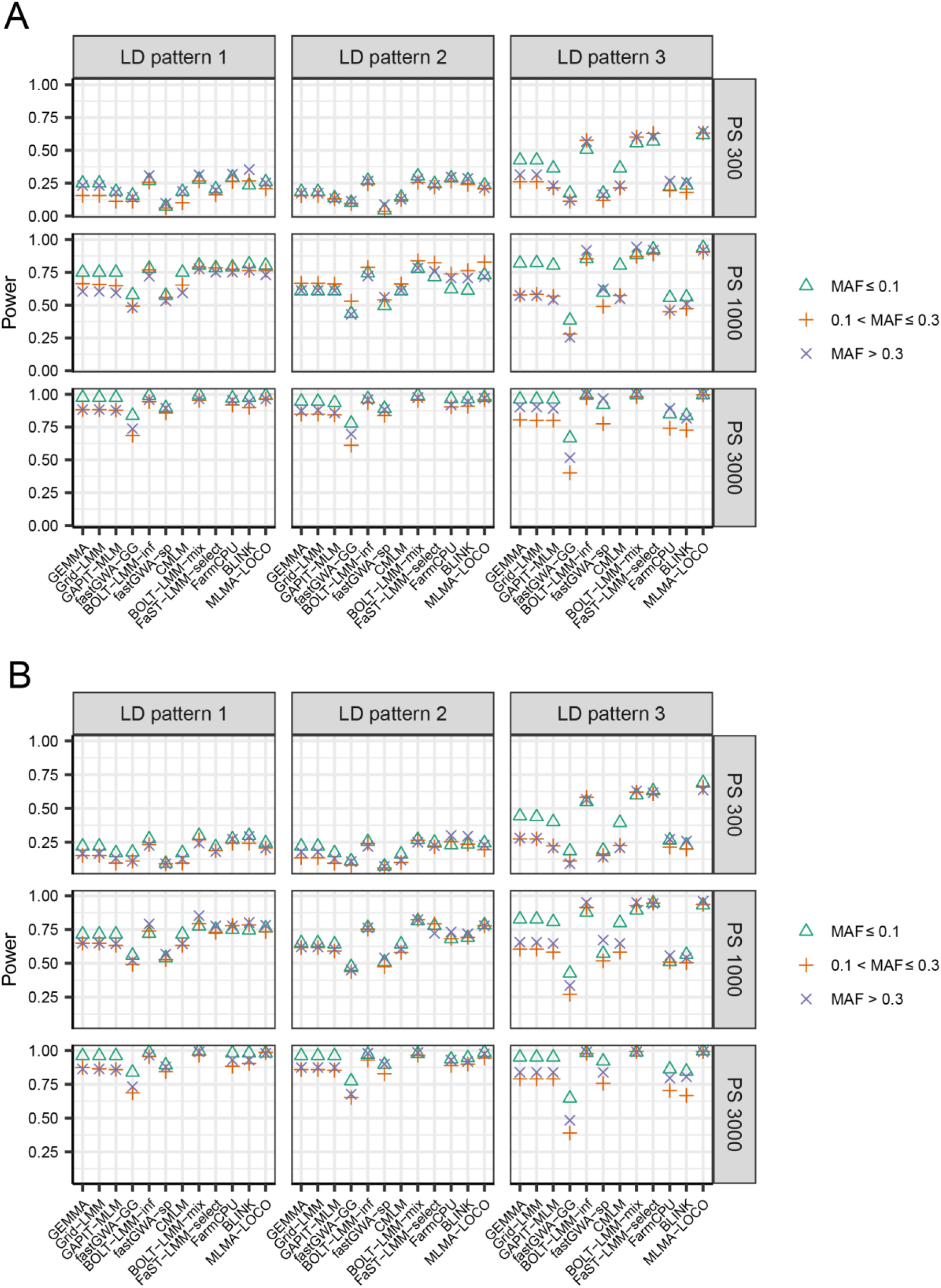
The statistical power of detecting QTL with a specific range of MAF for 12 GWAS algorithms evaluated in simulated data sets with 18 scenarios for trait heritability 0.7 with PG2 (each of the 6 major QTL explained 2% of the genetic variance). The 18 scenarios are combinations of three population sizes (PS 300, PS 1000 and PS 3000), three different LD patterns among the QTL (LD patterns 1-3), and two different genetic backgrounds (GB1 and GB2). In LD pattern 1, there is no LD between any two major QTL or between a major and a minor QTL. In LD pattern 2, there is no LD between any two major QTL, but LD exits between major and minor QTL. In LD pattern 3, there exists LD among the major QTL as well as between major and minor QTL. In GB1, there were 1,200 markers as minor QTL. In GB2, all markers on the chromosomes (LD patterns 2 and 3) or on half of the chromosomes (LD pattern 1) contributed as minor QTL. The results for GB1 and GB2 were shown in panel **A** and **B**, respectively. Each panel was further divided into 9 subpanels, each showing the results of a specific combination of population size and LD pattern. Within each subpanel, the results for QTL with three different ranges of MAF were indicated by different symbols. The algorithms CMLM and FaST-LMM-select were not evaluated for PS 3000 because the computational load was too high.

**Extended Data Figure 8.**
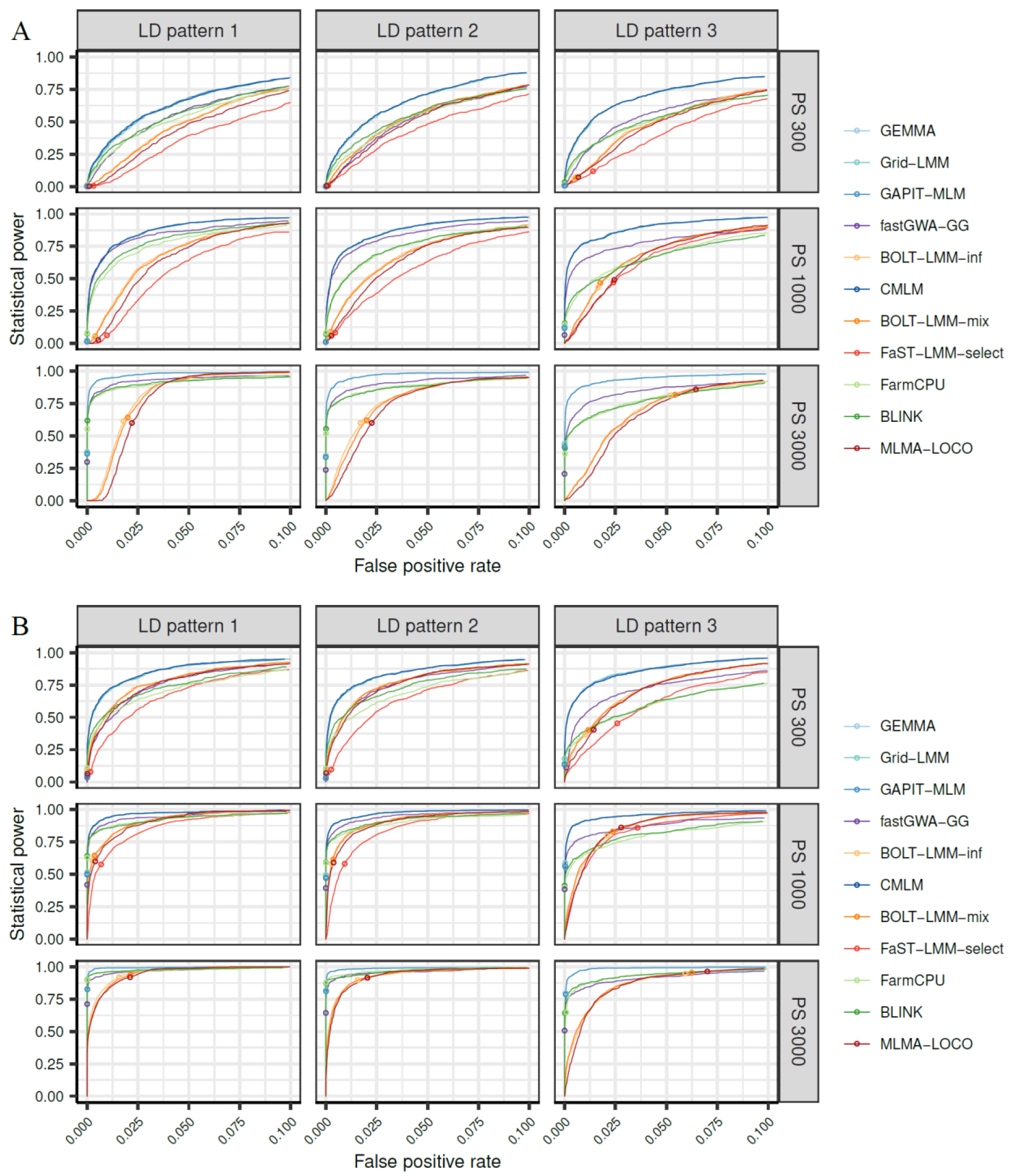
The receiver operating characteristic (ROC) curves of 11 GWAS algorithms evaluated in simulated data sets with 18 scenarios for trait heritability 0.5 with GB1 (1,200 markers contributed as minor QTL to the genetic background effects). The 18 scenarios are combinations of three population sizes (PS 300, PS 1000 and PS 3000), three different linkage disequilibrium (LD) patterns among the QTL (LD patterns 1-3), and two patterns of QTL effect sizes (PG1 and PG2). In LD pattern 1, there is no LD between any two major QTL or between a major and a minor QTL. In LD pattern 2, there is no LD between any two major QTL, but LD exits between major and minor QTL. In LD pattern 3, there exists LD among the major QTL as well as between major and minor QTL. In PG1, each of the 6 major QTL explained 2% of the genetic variance, In PG2, the 6 major QTL were randomly assigned to explain 2%, 4%, 6%, 8%, 10% and 12% of the genetic variance respectively. Results for PG1 and PG2 were shown in panels **A** and **B**, respectively. Each of the 9 subpanels showed the results for a specific combination of population size and data set. Within each subpanel, the ROC curves of different algorithms were shown in different colors. The power and FPR of each algorithm under the threshold of *p* < 0.05 after Bonferroni correction for multiple testing was indicated by a small circle on the curve. The algorithms CMLM and FaST-LMM-select were not evaluated for PS 3000 because the computational load was too high.

**Extended Data Figure 9.**
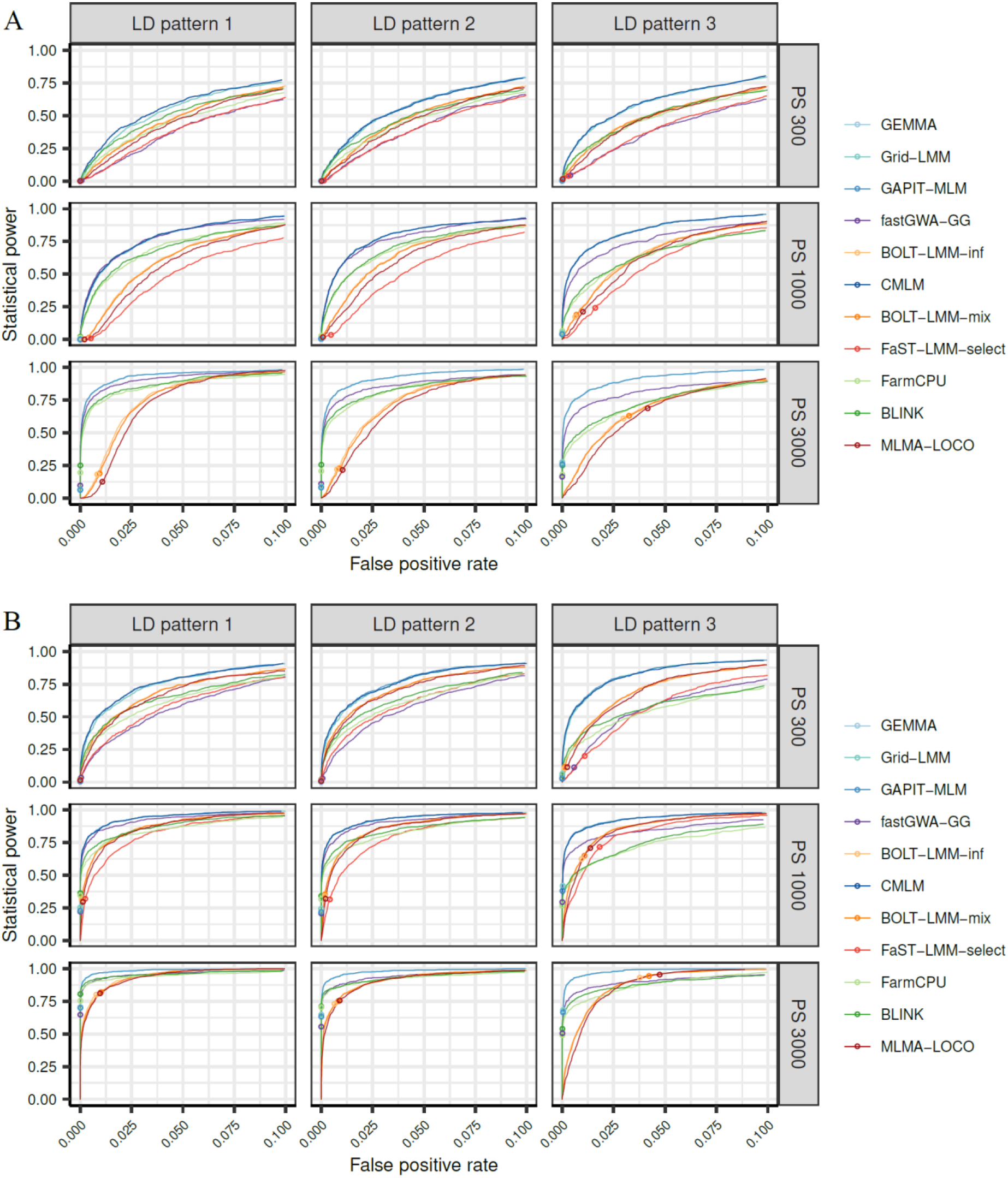
The receiver operating characteristic (ROC) curves of 11 GWAS algorithms evaluated in simulated data sets with 18 scenarios for trait heritability 0.3 with GB1 (1,200 markers contributed as minor QTL to the genetic background effects). The 18 scenarios are combinations of three population sizes (PS 300, PS 1000 and PS 3000), three different linkage disequilibrium (LD) patterns among the QTL (LD patterns 1-3), and two patterns of QTL effect sizes (PG1 and PG2). In LD pattern 1, there is no LD between any two major QTL or between a major and a minor QTL. In LD pattern 2, there is no LD between any two major QTL, but LD exits between major and minor QTL. In LD pattern 3, there exists LD among the major QTL as well as between major and minor QTL. In PG1, each of the 6 major QTL explained 2% of the genetic variance, In PG2, the 6 major QTL were randomly assigned to explain 2%, 4%, 6%, 8%, 10% and 12% of the genetic variance respectively. Results for PG1 and PG2 were shown in panels **A** and **B**, respectively. Each of the 9 subpanels showed the results for a specific combination of population size and data set. Within each subpanel, the ROC curves of different algorithms were shown in different colors. The power and FPR of each algorithm under the threshold of *p* < 0.05 after Bonferroni correction for multiple testing was indicated by a small circle on the curve. The algorithms CMLM and FaST-LMM-select were not evaluated for PS 3000 because the computational load was too high.

**Extended Data Figure 10.**
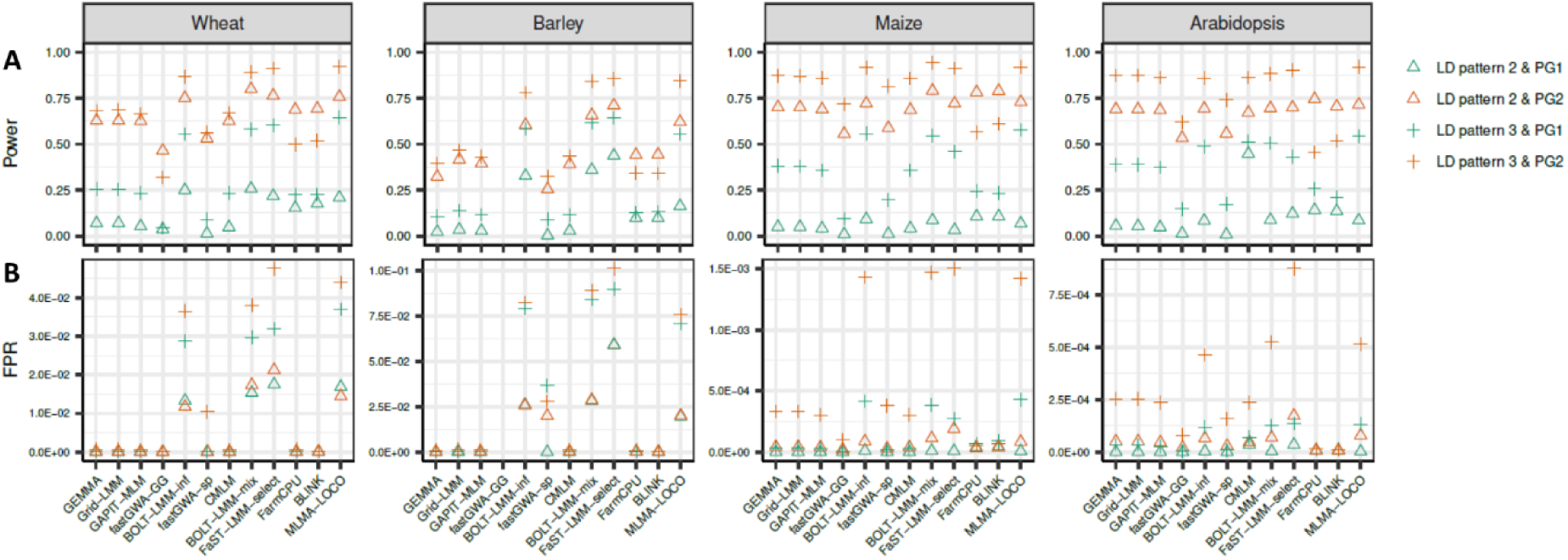
The statistical power (**A**) and false positive rate (**B**) of 12 GWAS algorithms evaluated in simulated data sets based on the genomic data of wheat, barley, maize, and Arabidopsis respectively. In the simulation, four scenarios for trait heritability 0.7 with GB1 (1,200 markers contributed as minor QTL to the genetic background effects) and a population size of 1,000 were considered. The four scenarios are combinations of two linkage disequilibrium (LD) patterns (LD pattern 2 and 3) and two cases of the proportion of genetic variance (PG) explained by the major QTL (PG1 and PG2). In LD pattern 2, there is no LD between any two major QTL, but LD exits between major and minor QTL. In LD pattern 3, there exists LD among the major QTL as well as between major and minor QTL. In PG1, each of the 6 major QTL explained 2% of the genetic variance, In PG2, the 6 major QTL were randomly assigned to explain 2%, 4%, 6%, 8%, 10% and 12% of the genetic variance respectively. Within each subpanel, the results of four scenarios were indicated by different symbols.

## References

1. Donnelly, P., Progress and challenges in genome-wide association studies in humans. Nature 456, 728–731 (2008).

2. Liu, X. et al., GWAS Atlas: an updated knowledgebase integrating more curated associations in plants and animals. Nucleic Acids Res. 51, D969–D976 (2023).

3. Tibbs Cortes, L., Zhang, Z., & Yu, J., Status and prospects of genome-wide association studies in plants. Plant Genome 14, e20077 (2021).

4. Devlin, B., & Roeder, K., Genomic control for association studies. Biometrics 55, 997–1004 (1999).

5. Pritchard, J. K., Stephens, M., Rosenberg, N. A., & Donnelly, P., Association mapping in structured populations. Am. J. Hum. Genet. 67, 170–181 (2000).

6. Price, A. L. e. a., Principal components analysis corrects for stratification in genome-wide association studies. Nat. Genet. 38, 904–909 (2006).

7. Yu, J. et al., A unified mixed-model method for association mapping that accounts for multiple levels of relatedness. Nat. Genet. 38, 203–208 (2006).

8. Zhao, K. et al., An Arabidopsis example of association mapping in structured samples. PLoS Genet. 3, e4 (2007).

9. Sul, J. H., Martin, L. S., & Eskin, E., Population structure in genetic studies: Confounding factors and mixed models. 14(12),. PLoS Genet. 14, e1007309 (2018).

10. McCulloch, C. E., & Searle, S. R., Generalized, linear, and mixed models. (John Wiley & Sons., 2004).

11. Kang, H. M. et al., Efficient control of population structure in model organism association mapping. Genetics 178, 1709–1723 (2008).

12. Lippert, C. et al., FaST linear mixed models for genome-wide association studies. Nat. Methods 8, 833–835 (2011).

13. Zhou, X., & Stephens, M., Genome-wide efficient mixed-model analysis for association studies. Nat. Genet. 44, 821–824 (2012).

14. Loh, P. R. et al., Efficient Bayesian mixed-model analysis increases association power in large cohorts. Nat. Genet. 47, 284–290 (2015).

15. Runcie, D. E., & Crawford, L., Fast and flexible linear mixed models for genome-wide genetics. PLoS Genet. 15, e1007978 (2019).

16. Laporte, F., Charcosset, A., & Mary-Huard, T., Efficient ReML inference in variance component mixed models using a Min-Max algorithm. PLoS Comput. Biol. 18, e1009659 (2022).

17. Zhang, Z. et al., Mixed linear model approach adapted for genome-wide association studies. Nat. Genet. 42, 355–360 (2010).

18. Li, M. et al., Enrichment of statistical power for genome-wide association studies. BMC Biol. 12, 73 (2014).

19. Aulchenko, Y. S. e. a., Genomewide rapid association using mixed model and regression: a fast and simple method for genomewide pedigree-based quantitative trait loci association analysis. Genetics 177, 577–585 (2007).

20. Svishcheva, G. R. et al., Rapid variance components–based method for whole-genome association analysis. Nat. Genet. 44, 1166–1170 (2012).

21. Jiang, L. e. a., A resource-efficient tool for mixed model association analysis of large-scale data. Nature genetics, 51(12), 1749–1755. Nat. Genet. 51, 1749-1755 (2019).

22. Kang, H. M. et al., Variance component model to account for sample structure in genome-wide association studies. Nat. Genet. 42, 348–354 (2010).

23. Listgarten, J. et al., Improved linear mixed models for genome-wide association studies. Nat. Methods 9, 525–526 (2012).

24. Widmer, C. et al., Further improvements to linear mixed models for genome-wide association studies. Sci. Rep. 4, 6874 (2014).

25. Segura, V. et al., An efficient multi-locus mixed-model approach for genome-wide association studies in structured populations. 825. Nat. Genet. 44, 825–830 (2012).

26. Wang, Q. et al., A SUPER powerful method for genome wide association study. PloS one, 9(9), PLoS One 9, e107684 (2014).

27. Liu, X. et al., Iterative usage of fixed and random effect models for powerful and efficient genome-wide association studies. PLoS Genet. 12, e1005767 (2016).

28. Huang, M. et al., BLINK: a package for the next level of genome-wide association studies with both individuals and markers in the millions. GigaScience 8, giy154 (2019).

29. Mbatchou, J. et al., Computationally efficient whole-genome regression for quantitative and binary traits. Nat. Genet. 53, 1097–1103 (2021).

30. Eu-ahsunthornwattana J. et al., Comparison of methods to account for relatedness in genome-wide association studies with family-based data. PLoS Genet. 10, e1004445 (2014).

31. Bradbury, P. J. et al., TASSEL: software for association mapping of complex traits in diverse samples. Bioinformatics 23, 2633–2635 (2007).

32. Zhou, H., Hu, L., Zhou, J., & Lange, K., MM Algorithms for Variance Components Models. J. Comput. Graph. Stat. 28, 350–361 (2019).

33. Lipka, A. E. et al., GAPIT: genome association and prediction integrated tool. Bioinformatics 28, 2397–2399 (2012).

34. Tang, Y. et al., GAPIT version 2: an enhanced integrated tool for genomic association and prediction. Plant Genome 9, plantgenome2015.11.0120 (2016).

35. Wang, J., & Zhang, Z., GAPIT Version 3: boosting power and accuracy for genomic association and prediction.. Genomics Proteomics Bioinformatics 19, 629–640 (2021).

36. Yang, J., Lee, S. H., Goddard, M. E., & Visscher, P. M., GCTA: a tool for genome-wide complex trait analysis. Am. J. Hum. Genet. 88, 76–82 (2011).

37. Aulchenko, Y. S., Ripke, S., Isaacs, A., & Van Duijn, C. M., GenABEL: an R library for genome-wide association analysis. Bioinformatics 23, 1294–1296 (2007).

38. Yang, J., Zaitlen, N. A., Goddard, M. E., Visscher, P. M., & Price, A. L., Advantages and pitfalls in the application of mixed-model association methods. Nat. Genet. 46, 100–106 (2014).

39. Dunn, O. J., Multiple comparisons among means.. J. Am. Stat. Assoc. 56, 52–64 (1961).

40. Benjamini, Y., & Hochberg, Y., Controlling the false discovery rate: a practical and powerful approach to multiple testing. J. R. Stat. Soc. Ser. B Methodol. 57, 289–300 (1995).

41. van den Berg, S., Vandenplas, J., van Eeuwijk, F. A., Lopes, M. S., & Veerkamp, R. F., Significance testing and genomic inflation factor using high-density genotypes or whole-genome sequence data. J. Anim. Breed. Genet. 136, 418–429 (2019).

42. Erickson, P. A. et al., Unique genetic signatures of local adaptation over space and time for diapause, an ecologically relevant complex trait, in Drosophila melanogaster. PLoS Genet. 16, e1009110 (2020).

43. Loh, P. R. et al., Mixed-model association for biobank-scale datasets. Nat Genet. 50, 906–908 (2018).

44. Sipser, M., Introduction to the theory of computation., 3rd ed. (Cengage Learning, 2012).

45. Gianola, D., Priors in whole-genome regression: the Bayesian alphabet returns. Genetics 194, 573–596 (2013).

46. The 1001 Genomes Consortium., 1,135 genomes reveal the global pattern of polymorphism in Arabidopsis thaliana. Cell 166, 481–491 (2016).

47. Howie, B. N., Donnelly, P., & Marchini, J., A flexible and accurate genotype imputation method for the next generation of genome-wide association studies. PLoS Genet. 5, e1000529 (2009).

48. Schulthess, A. W. et al., Genomics-informed prebreeding unlocks the diversity in genebanks for wheat improvement.. Nat. Genet. 54, 1544–1552 (2022).

49. Wang, W. et al., Genomic variation in 3,010 diverse accessions of Asian cultivated rice.. Nature 557, 43–49 (2018).

50. Browning, B. L., Zhou, Y., & Browning, S. R., A one-penny imputed genome from next generation reference panels. Am. J. Hum. Genet. 103, 338–348 (2018).

51. Romay, M. C. et al., Comprehensive genotyping of the USA national maize inbred seed bank. Genome Biol. 14, R55 (2013).

52. Milner, S. G. et al., Genebank genomics highlights the diversity of a global barley collection. Nat. Genet. 51, 319–326 (2019).

53. Swarts, K. et al., Novel methods to optimize genotypic imputation for low-coverage, next-generation sequence data in crop plants. Plant Genome 7, plantgenome2014.05.0023 (2014).

54. Gonzalez, M. Y. et al., Unbalanced historical phenotypic data from seed regeneration of a barley ex situ collection. Sci. Data 5, 180278 (2018).

55. Remington, D. L. et al., Structure of linkage disequilibrium and phenotypic associations in the maize genome. Proc. Nat. Acad. Sci. U.S.A. 98, 11479–11484 (2001).

56. Bouché, F., Lobet, G., Tocquin, P., & Périlleux, C., FLOR-ID: an interactive database of flowering-time gene networks in Arabidopsis thaliana.. Nucleic Acids Res. 44, D1167–D1171 (2016).

