## Supplementary Information for "A comprehensive overview and benchmarking analysis of fast algorithms for genome-wide association studies"

### Supplementary Information for the manuscript "A comprehensive overview and benchmarking analysis of fast algorithms for genome-wide association studies"

Fang Liu[1,2,4#], Jie Zhang[3,4#], Renate H. Schmidt[4], Martin Mascher[4], Yusheng Zhao[4], Jochen C. Reif[4,*], Yong Jiang[4,*]

[1] Key Laboratory of Plant Germplasm Enhancement and Specialty Agriculture, Wuhan Botanical Garden, Chinese Academy of Sciences, Wuhan 430074, China

[2] Center of Economic Botany, Core Botanical Gardens, Chinese Academy of Sciences, Wuhan 430074, China

[3] Key Laboratory of Molecular Genetics, Guizhou Institute of Tobacco Science, Guiyang 550081, China

[4] Leibniz Institute of Plant Genetics and Crop Plant Research (IPK), Corrensstr. 3, 06466 Stadt Seeland OT Gatersleben, Germany

#### SUPPLEMENTARY NOTE A

##### 1. Linear mixed model for genome-wide association study

1.1. **The Q+K linear mixed model.** For simplicity, we presented the model in the case that each individual has only one phenotypic observation. The Q+K linear mixed model is as follows:

$$\boldsymbol{y} = \boldsymbol{X}\boldsymbol{\beta} + \boldsymbol{m}a + \boldsymbol{g} + \boldsymbol{e}, \tag{1}$$

where $\boldsymbol{y}$ is the $n$-dimensional vector of observation phenotypic values, $\boldsymbol{\beta}$ is the $k$-dimensional vector of covariates which may include a common intercept, environmental and/or subpopulation effects etc., $\boldsymbol{X}$ is the corresponding $n \times k$ design matrix, $a$ is the effect of the marker being tested, $\boldsymbol{m}$ is the marker coding vector, $\boldsymbol{g}$ is the $n$-dimensional vector of polygenic effects, and $\boldsymbol{e}$ is the n-dimensional vector of residuals.

In the model, $\boldsymbol{\beta}$ and $a$ are treated as fixed effects, $\boldsymbol{g}$ and $\boldsymbol{e}$ are random effects with multi-variate normal distribution: $\boldsymbol{g} \sim \mathcal{N}(0, \boldsymbol{K}\sigma_g^2)$, $\boldsymbol{e} \sim \mathcal{N}(0, \boldsymbol{I}\sigma_e^2)$, where $\boldsymbol{K}$ is a kinship matrix derived from pedigree/marker information, $\boldsymbol{I}$ is the identity matrix, $\sigma_g^2$ and $\sigma_e^2$ are the corresponding variance components. In most GWAS algorithm, $\boldsymbol{K}$ is calculated as $\boldsymbol{M}\boldsymbol{M}'/c$, where $\boldsymbol{M}$ is the $n \times p$ matrix of markers. Usually, the entries in $\boldsymbol{M}$ is coded as 0, 1, 2 (the number of copies of a reference allele) and it is centered or normalized with respect to the column.

1.2. **The general procedure of GWAS.** For convenience, we could also put the fixed effects in Eq. (1) together. Namely let $\boldsymbol{W} = (\boldsymbol{X}, \boldsymbol{m})$ and $\boldsymbol{\alpha} = (\boldsymbol{\beta}', a)'$. Then Eq. (1) can be rewritten as

$$\boldsymbol{y} = \boldsymbol{W}\boldsymbol{\alpha} + \boldsymbol{g} + \boldsymbol{e}. \tag{2}$$

The procedure of GWAS can be roughly divided into two steps: 1) solving the linear mixed model; 2) producing the test statistics. In the first step, the unknown parameters are estimated via maximum likelihood (ML) or restricted maximum likelihood (REML) method. Here we take the ML method as an example. Assuming $\delta = \sigma_g^2/\sigma_e^2$, the log-likelihood function is as follows:

$$LL(\boldsymbol{\alpha}, \delta, \sigma_e^2) = -\frac{n}{2}\log(2\pi\sigma_e^2) - \frac{1}{2}\log|\boldsymbol{V}| - \frac{1}{2\sigma_e^2}(\boldsymbol{y} - \boldsymbol{W}\boldsymbol{\alpha})'\boldsymbol{V}^{-1}(\boldsymbol{y} - \boldsymbol{W}\boldsymbol{\alpha}), \tag{3}$$

where $\boldsymbol{V} = \delta\boldsymbol{K} + \boldsymbol{I}$, $|\cdot|$ denote the determinant of a matrix.

After estimating the unknown parameters, the test statistics can be produced with different approaches, e.g. the likelihood ratio test and various score tests. Note that we are only interested in the significance of marker effect, namely $\hat{a}$ which is typically the last entry of the vector $\hat{\boldsymbol{\alpha}} = (\hat{\boldsymbol{\beta}}', \hat{a})'$. Taking the Wald-test as an example, the test statistic has the following form

$$T_{Wald} = \frac{\hat{a}^2}{\text{var}(\hat{a})}. \tag{4}$$

Under the null hypothesis, the test statistic asymptotically follows a $\chi^2$-distribution with one-degree of freedom.

1.3. **Solving the linear mixed model.** For the log-likelihood function (Eq. 3) to reach its maximum, it is necessary that the partial derivatives with respect to $\boldsymbol{\alpha}$ and $\sigma_e^2$ are zero. This gives us the following results:

$$\hat{\boldsymbol{\alpha}} = (\boldsymbol{W}'\boldsymbol{V}^{-1}\boldsymbol{W})^{-1}\boldsymbol{W}'\boldsymbol{V}^{-1}\boldsymbol{y}, \tag{5}$$

$$\hat{\sigma}_e^2 = \frac{1}{n}(\boldsymbol{y} - \boldsymbol{W}\boldsymbol{\alpha})'\boldsymbol{V}^{-1}(\boldsymbol{y} - \boldsymbol{W}\boldsymbol{\alpha}) = \frac{1}{n}\boldsymbol{y}'\boldsymbol{H}\boldsymbol{y}, \tag{6}$$

where $\boldsymbol{H} = \boldsymbol{V}^{-1} - \boldsymbol{V}^{-1}\boldsymbol{W}(\boldsymbol{W}'\boldsymbol{V}^{-1}\boldsymbol{W})^{-1}\boldsymbol{W}'\boldsymbol{V}^{-1}$.

Now, replacing $\boldsymbol{\alpha}$ and $\sigma_e^2$ in Eq. (3) by $\hat{\boldsymbol{\alpha}}$ and $\hat{\sigma}_e^2$, the log-likelihood function is as follows:

$$\begin{aligned} LL(\delta) &= -\frac{n}{2}[1+\log(\frac{2\pi}{n})+\log((\boldsymbol{y}-\boldsymbol{W}\hat{\boldsymbol{\alpha}})'\boldsymbol{V}^{-1}(\boldsymbol{y}-\boldsymbol{W}\hat{\boldsymbol{\alpha}}))]-\frac{1}{2}\log|\boldsymbol{V}| \\ &= -\frac{n}{2}[1+\log(\frac{2\pi}{n})+\log(\boldsymbol{y}'\boldsymbol{H}\boldsymbol{y})]-\frac{1}{2}\log|\boldsymbol{V}| \end{aligned} \tag{7}$$

For $LL(\delta)$ to reach its maximum, it is necessary that its first derivative with respect to $\delta$ is zero. The first derivative of $LL(\delta)$ is calculated as the following:

$$\frac{dLL(\delta)}{d\delta} = -\frac{1}{2}\operatorname{tr}(\boldsymbol{V}^{-1}\boldsymbol{K})+\frac{n}{2}\cdot\frac{\boldsymbol{y}'\boldsymbol{HKHy}}{\boldsymbol{y}'\boldsymbol{Hy}}, \tag{8}$$

where tr denote the trace of a matrix (i.e. the sum of all diagonal elements).

Now, it remains to solve $\delta$ by equating Eq. (9) to 0. Unfortunately, it is difficult to find the analytic solutions except in special cases. Thus, numerical methods such as interval bisection, secant method and Newton-Raphson are used, which means a certain number of iterations is unavoidable. During this procedure, the most time-consuming part is calculating $\boldsymbol{V}^{-1}$, $|\boldsymbol{V}|$ and multiplications of matrices with size $n \times n$, whose time complexity is $\mathcal{O}(n^3)$. Moreover, these calculations have to be performed for each iteration and for each marker. Therefore, the total time complexity is $\mathcal{O}(ptn^3)$, where $t$ is the average number of iterations.

1.4. **Producing the test statistics.** To calculate the test statistics (Eq. (4)), we need $\hat{a}$ and $\operatorname{var}(\hat{a})$. Note that Eq. (5) can be reformulated as

$$\begin{pmatrix}\hat{\boldsymbol{\beta}}\\ \hat{a}\end{pmatrix} = \begin{pmatrix}\boldsymbol{X}'\boldsymbol{V}^{-1}\boldsymbol{X} & \boldsymbol{X}'\boldsymbol{V}^{-1}\boldsymbol{m}\\ \boldsymbol{m}'\boldsymbol{V}^{-1}\boldsymbol{X} & \boldsymbol{m}'\boldsymbol{V}^{-1}\boldsymbol{m}\end{pmatrix}^{-1}\begin{pmatrix}\boldsymbol{X}'\boldsymbol{V}^{-1}\boldsymbol{y}\\ \boldsymbol{m}'\boldsymbol{V}^{-1}\boldsymbol{y}\end{pmatrix} \tag{9}$$

from which we can solve for $\hat{a}$:

$$\hat{a} = \frac{\boldsymbol{m}'\boldsymbol{V}^{-1}\boldsymbol{y}-\boldsymbol{m}'\boldsymbol{V}^{-1}\boldsymbol{X}(\boldsymbol{X}'\boldsymbol{V}^{-1}\boldsymbol{X})^{-1}\boldsymbol{X}'\boldsymbol{V}^{-1}\boldsymbol{y}}{\boldsymbol{m}'\boldsymbol{V}^{-1}\boldsymbol{m}-\boldsymbol{m}'\boldsymbol{V}^{-1}\boldsymbol{X}(\boldsymbol{X}'\boldsymbol{V}^{-1}\boldsymbol{X})^{-1}\boldsymbol{X}'\boldsymbol{V}^{-1}\boldsymbol{m}} = \frac{\boldsymbol{m}'\boldsymbol{Py}}{\boldsymbol{m}'\boldsymbol{Pm}} \tag{10}$$

where $P = \boldsymbol{V}^{-1}-\boldsymbol{V}^{-1}\boldsymbol{X}(\boldsymbol{X}'\boldsymbol{V}^{-1}\boldsymbol{X})^{-1}\boldsymbol{X}'\boldsymbol{V}^{-1}$.

From standard theory of LMMs (e.g. Henderson 1975), we know that

$$\operatorname{cov}\begin{pmatrix}\hat{\boldsymbol{\beta}}\\ \hat{a}\end{pmatrix} = \sigma_e^2\begin{pmatrix}\boldsymbol{X}'\boldsymbol{V}^{-1}\boldsymbol{X} & \boldsymbol{X}'\boldsymbol{V}^{-1}\boldsymbol{m}\\ \boldsymbol{m}'\boldsymbol{V}^{-1}\boldsymbol{X} & \boldsymbol{m}'\boldsymbol{V}^{-1}\boldsymbol{m}\end{pmatrix}^{-1} \tag{11}$$

from which we can deduce the following (replacing $\sigma_e^2$ by $\hat{\sigma}_e^2$)

$$\operatorname{var}(\hat{a}) = \frac{\hat{\sigma}_e^2}{\boldsymbol{m}'\boldsymbol{Pm}} = \frac{1}{n}\cdot\frac{\boldsymbol{y}'\boldsymbol{Py}}{\boldsymbol{m}'\boldsymbol{Pm}}. \tag{12}$$

Now, using Eq. (10) and (12), we can calculate the Wald-test statistic (Eq. (4)) as following:

$$T_{Wald} = \frac{1}{\hat{\sigma}_e^2}\cdot\frac{(\boldsymbol{m}'\boldsymbol{Py})^2}{\boldsymbol{m}'\boldsymbol{Pm}} = \frac{n(\boldsymbol{m}'\boldsymbol{Py})^2}{(\boldsymbol{m}'\boldsymbol{Pm})(\boldsymbol{y}'\boldsymbol{Py})}. \tag{13}$$

Since the matrix $\boldsymbol{V}^{-1}$ has been calculated when solving the LMM, so here we only need to consider the matrix-vector multiplications, whose complexity is $\mathcal{O}(n^2)$. Taking all markers into consideration, the time complexity is $\mathcal{O}(pn^2)$. Therefore, if we only consider the most time-consuming part, the time complexity for the entire GWAS procedure is roughly $\mathcal{O}(ptn^3)+\mathcal{O}(pn^2)$ with a standard algorithm.

#### 2. A detailed review of fast GWAS algorithms

2.1. **FaST-LMM.** FaST-LMM is one of the earliest fast GWAS algorithms. Although the its original algorithm was not evaluated in our benchmarking analysis, its variant FaST-LMM-select was. Thus, in the following, we first introduce the original algorithm and then FaST-LMM-select.

2.1.1. *The original algorithm of FaST-LMM.* The key idea of FaST-LMM [8] is to avoid repeating the time-consuming computation of $\boldsymbol{V}^{-1}$, $|\boldsymbol{V}|$ and matrix multiplications of size $n \times n$ for each round of iteration and for each marker. This goal is achieved by applying spectral decomposition to the matrix $\boldsymbol{K}$. Namely we have $\boldsymbol{K} = \boldsymbol{U}\boldsymbol{\Lambda}\boldsymbol{U}'$, where $\boldsymbol{U}$ is an $n \times n$ orthogonal matrix consisting of eigenvectors for $\boldsymbol{K}$, $\boldsymbol{\Lambda}$ is a diagonal matrix whose diagonal entries are the corresponding eigenvalues $\xi_1, \xi_2, \ldots, \xi_n$. Note that, the columns of $\boldsymbol{U}$ are also eigenvectors of $\boldsymbol{V} = \delta\boldsymbol{K} + \boldsymbol{I}$ with the corresponding eigenvalues $\delta\xi_1 + 1$, $\delta\xi_2 + 1$, ..., $\delta\xi_n + 1$. Thus, we also obtain the spectral decomposition for $\boldsymbol{V}$. That is,

$$\boldsymbol{V} = \boldsymbol{U}(\delta\boldsymbol{\Lambda} + \boldsymbol{I})\boldsymbol{U}' = \boldsymbol{U}\,\mathrm{diag}(\delta\xi_1 + 1, \delta\xi_2 + 1, \ldots, \delta\xi_n + 1)\boldsymbol{U}' \tag{14}$$

Now we make linear transformations to the data. Let $\tilde{\boldsymbol{y}} = \boldsymbol{U}'\boldsymbol{y}$, $\tilde{\boldsymbol{W}} = \boldsymbol{U}'\boldsymbol{W}$. Eq. (5) can be simplified as following:

$$\begin{aligned} \hat{\boldsymbol{\alpha}} &= (\tilde{\boldsymbol{W}}'(\delta\boldsymbol{\Lambda} + \boldsymbol{I})^{-1}\tilde{\boldsymbol{W}})^{-1}\tilde{\boldsymbol{W}}'(\delta\boldsymbol{\Lambda} + \boldsymbol{I})^{-1}\tilde{\boldsymbol{y}} \\ &= \left(\sum_{i=1}^{n} \frac{\tilde{\boldsymbol{W}}_i'\tilde{\boldsymbol{W}}_i}{\delta\xi_i + 1}\right)^{-1} \sum_{i=1}^{n} \frac{\tilde{\boldsymbol{W}}_i'\tilde{\boldsymbol{y}}_i}{\delta\xi_i + 1} \end{aligned} \tag{15}$$

where $\tilde{\boldsymbol{W}}_i$ denotes the $i$-th row of $\tilde{\boldsymbol{W}}$ (hence an $k+1$-dimensional vector) and $\tilde{\boldsymbol{y}}_i$ is the $i$-th entry of $\tilde{\boldsymbol{y}}$. Performing the linear transformation $\boldsymbol{U}'\boldsymbol{y}$ and $\boldsymbol{U}'\boldsymbol{W}$ requires $\mathcal{O}(n^2)$ time. Once the transformation is done, Eq. (15) can be evaluated in $\mathcal{O}(kn) + \mathcal{O}(k^2n) + \mathcal{O}(k^3) \sim \mathcal{O}(n)$ time (Since $k$ is a constant and typically much smaller than $n$).

Furthermore, Eq. (7) can be simplified as:

$$\begin{aligned} LL(\delta) &= -\frac{n}{2}[1 + \log(\frac{2\pi}{n}) + \log((\boldsymbol{y} - \boldsymbol{W}\hat{\boldsymbol{\alpha}})'\boldsymbol{V}^{-1}(\boldsymbol{y} - \boldsymbol{W}\hat{\boldsymbol{\alpha}}))] - \frac{1}{2}\log|\boldsymbol{V}| \\ &= -\frac{n}{2}[1 + \log(\frac{2\pi}{n}) + \log(\sum_{i=1}^{n} \frac{(\tilde{\boldsymbol{y}}_i - \tilde{\boldsymbol{W}}_i\hat{\boldsymbol{\alpha}})^2}{\delta\xi_i + 1})] - \frac{1}{2}\sum_{i=1}^{n}\log(\delta\xi_i + 1) \end{aligned} \tag{16}$$

Again, we see that evaluating the above equation only requires $\mathcal{O}(n)$ time. Finally, $\hat{\delta} = \mathrm{argmax}(LL(\delta))$ is solved by using a one-dimensional numerical optimization approach based on the Brent's method.

To summarize, after applying spectral decomposition for the matrix $\boldsymbol{K}$ and making linear transformations $\boldsymbol{U}'\boldsymbol{y}$ and $\boldsymbol{U}'\boldsymbol{W}$, FaST-LMM is able to evaluate the log-likelihood function in each round of iteration for each marker in $\mathcal{O}(n)$ time. The time complexity for the spectral decomposition is $\mathcal{O}(n^3)$ and the linear transformations $\boldsymbol{U}'\boldsymbol{y}$ and $\boldsymbol{U}'\boldsymbol{W} = \boldsymbol{U}'(\boldsymbol{X}, \boldsymbol{m})$ requires $\mathcal{O}(kn^2) \sim \mathcal{O}(n^2)$ time. For different markers, the vector $\boldsymbol{m}$ changes. So, taking all markers into consideration, the linear transformations take $O(pn^2)$ time. Thus, we see that in the step of solving the LMM, FaST-LMM reduces the time complexity from $\mathcal{O}(ptn^3)$ to $\mathcal{O}(n^3) + \mathcal{O}(pn^2) + \mathcal{O}(ptn)$.

After the linear transformation is done, producing the test statistics (Eq. (13)) only takes $\mathcal{O}(n)$ time for each marker. This is because

$$\begin{aligned} \boldsymbol{m}'\boldsymbol{P}\boldsymbol{m} &= \boldsymbol{m}'\boldsymbol{V}^{-1}\boldsymbol{m} - \boldsymbol{m}'\boldsymbol{V}^{-1}\boldsymbol{X}(\boldsymbol{X}'\boldsymbol{V}^{-1}\boldsymbol{X})^{-1}\boldsymbol{X}'\boldsymbol{V}^{-1}\boldsymbol{m} \\ &= \tilde{\boldsymbol{m}}'(\delta\boldsymbol{\Lambda} + \boldsymbol{I})^{-1}\tilde{\boldsymbol{m}} - \tilde{\boldsymbol{m}}'(\delta\boldsymbol{\Lambda} + \boldsymbol{I})^{-1}\tilde{\boldsymbol{X}}(\tilde{\boldsymbol{X}}'(\delta\boldsymbol{\Lambda} + \boldsymbol{I})^{-1}\tilde{\boldsymbol{X}})^{-1}\tilde{\boldsymbol{X}}'(\delta\boldsymbol{\Lambda} + \boldsymbol{I})^{-1}\tilde{\boldsymbol{m}} \\ &= \left(\sum_{i=1}^{n} \frac{\tilde{\boldsymbol{m}}_i^2}{\delta\xi_i + 1}\right) - \left(\sum_{i=1}^{n} \frac{\tilde{\boldsymbol{m}}_i\tilde{\boldsymbol{X}}_i}{\delta\xi_i + 1}\right)\left(\sum_{i=1}^{n} \frac{\tilde{\boldsymbol{X}}_i'\tilde{\boldsymbol{X}}_i}{\delta\xi_i + 1}\right)^{-1}\left(\sum_{i=1}^{n} \frac{\tilde{\boldsymbol{X}}_i'\tilde{\boldsymbol{m}}_i}{\delta\xi_i + 1}\right) \end{aligned} \tag{17}$$

where $\tilde{\boldsymbol{m}}_i$ is the $i$-th entry of the transformed vector $\tilde{\boldsymbol{m}} = \boldsymbol{U}'\boldsymbol{m}$, $\tilde{\boldsymbol{X}}_i$ is the $i$-th row of the transformed matrix $\tilde{\boldsymbol{X}} = \boldsymbol{U}'\boldsymbol{X}$. From the above derivation, we can see that evaluating the term $\boldsymbol{m}'\boldsymbol{P}\boldsymbol{m}$ requires $\mathcal{O}(kn) + \mathcal{O}(k^2n) + \mathcal{O}(k^3) \sim \mathcal{O}(n)$ time. Similar arguments indicated that evaluating the term $\boldsymbol{m}'\boldsymbol{P}\boldsymbol{y}$ also requires $\mathcal{O}(n)$ time. Hence, producing the test statistics for all markers takes $\mathcal{O}(pn)$ time.

Therefore, the overall time complexity for FaST-LMM is dominated by the complexity of solving the LMM, which is roughly $\mathcal{O}(n^3) + \mathcal{O}(pn^2) + \mathcal{O}(ptn)$.

2.1.2. *FaST-LMM-select.* This is an improved version of the original FaST-LMM algorithm which aims at increasing the power of detection [9] [16]. The idea is to select a subset of markers to generate the kinship matrix $\boldsymbol{K}$ instead of using all markers. Namely, let $s$ be the number of selected markers and $\boldsymbol{M}_s$ be the submatrix of $\boldsymbol{M}$ consisting of the $s$ columns of selected markers. Then, FaST-LMM-select replaces $\boldsymbol{K} = \boldsymbol{M}\boldsymbol{M}'$ by $\boldsymbol{K}_s = \boldsymbol{M}_s\boldsymbol{M}'_s$. All other settings are the same as the original FaST-LMM.

The procedure of selecting markers is as follows: 1) Dividing the data samples (individuals) randomly into training and test sets. 2) Fitting the markers one-by-one in a univariate linear regression model and obtaining their $p$-values for the training set. 3) Sorting the markers by an ascending order of $p$-values. 4) For $s_c \in \{0, 1, 2, 4, \ldots, 1024, p\}$ ($p$ is the number of all markers), using the first $s$ markers to generate the matrix $\boldsymbol{K}$, solving the LMM of the null Q+K model (27) in the training set and evaluating the log-likelihood of function for the test set. 5) Repeating the procedures from 1) to 4) for several different partitions of training and test sets, choosing the number $s$ which maximizes the sum of log-likelihood across all test sets.

There is a further improved version FaST-LMM-all+select [16] as it was shown that in some cases the type I error rate of FaST-LMM-Select is significantly higher than the original FaST-LMM. FaST-LMM-all+select uses a linear combination of two matrices, one generated from all markers and the other from selected ones, as the kinship matrix. More precisely, FaST-LMM-all+select assumes that $\boldsymbol{g} \sim \mathcal{N}(0, \boldsymbol{K}_c\sigma_g^2)$, where $\boldsymbol{K}_c = \pi\boldsymbol{K} + (1-\pi)\boldsymbol{K}_s$, $0 \leq \pi \leq 1$. Thus, there are two parameters to be determined, namely the number of selected markers $s$ and the mixture weight $\pi$. The procedure of selecting markers is similar to that of FaST-LMM-Select, but differs in three aspects. First, the markers are tested one-by-one in the Q+K model with $\boldsymbol{K}$ as the kinship matrix instead of a univariate linear regression model. Second, the pool of marker numbers to be selected is smaller, by default $\{0, 1, 2, 4, \ldots, 128\}$, because there is a component of $\boldsymbol{K}$ generated by all markers in the mixture. Last but not least, the mixture weight $\pi$ is to be optimized in the evaluation of log-likelihood of the LMM for the training set.

For the computational load, it is clear that FaST-LMM-Select and FaST-LMM-all+select takes additional time in the procedure of selecting markers, which involves repeatedly solving the LMM for different training sets to optimize the number of selected markers. However, in FaST-LMM-Select one can avoid the explicit formation of the matrix $\boldsymbol{K}_s$, if $s$ is smaller than $n$. Namely, to replace the spectral decomposition of $\boldsymbol{K}_s$ by the singular value decomposition of $\boldsymbol{M}_s$ [8]. Thus, for any $s$ selected markers, the time complexity of solving the LMM is reduced from $\mathcal{O}(n^3) + \mathcal{O}(pn^2) + \mathcal{O}(tn)$ to $\mathcal{O}(s^2n) + \mathcal{O}(tn)$ where $t$ is the average number of iterations. This advantage cannot be exploited in FaST-LMM-all+select as one has to first combine the two kinship matrices and then perform the spectral decomposition. Thus, the total time required for FaST-LMM-Select relative to FaST-LMM depends on several parameters (e.g. the number of samplings for the training-test sets and the number of selected markers), while FaST-LMM-all+select is expected to be slower than FaST-LMM.

2.2. **GEMMA.** The general idea of GEMMA [20] is similar to FaST-LMM, namely performing the spectral decomposition to the matrix $\boldsymbol{K}$ once and then reducing the time complexity of evaluating the likelihood function to $\mathcal{O}(n)$ in each iteration for each marker. But the way of implementation is different from FaST-LMM. Below we sketch the essentials.

In addition to the log-likelihood function (Eq. (7)) and its first derivative (Eq. (8)), GEMMA also evaluates the second derivative, which is of the following form:

$$\frac{d^2LL(\delta)}{d\delta^2} = \frac{1}{2}\,\mathrm{tr}(\boldsymbol{V}^{-1}\boldsymbol{K}\boldsymbol{V}^{-1}\boldsymbol{K}) - \frac{n}{2}\cdot\frac{2(\boldsymbol{y}'\boldsymbol{HKHKHy})(\boldsymbol{y}'\boldsymbol{Hy}) - (\boldsymbol{y}'\boldsymbol{HKHy})^2}{\boldsymbol{y}'\boldsymbol{Hy}^2}. \tag{18}$$

To evaluate Eq. (7), (8) and (18), we need to efficiently evaluate $|\boldsymbol{V}|$, $\mathrm{tr}(\boldsymbol{V}^{-1}\boldsymbol{K})$, $\mathrm{tr}(\boldsymbol{V}^{-1}\boldsymbol{K}\boldsymbol{V}^{-1}\boldsymbol{K})$, $\boldsymbol{y}'\boldsymbol{Hy}$, $\boldsymbol{y}'\boldsymbol{HKHy}$ and $\boldsymbol{y}'\boldsymbol{HKHKHy}$. After some mathematical derivations, it turns out that we only need to efficiently evaluate $|\boldsymbol{V}|$, $\mathrm{tr}(\boldsymbol{V}^{-1})$, $\mathrm{tr}(\boldsymbol{V}^{-1}\boldsymbol{V}^{-1})$, $\boldsymbol{y}'\boldsymbol{Hy}$, $\boldsymbol{y}'\boldsymbol{HHy}$ and $\boldsymbol{y}'\boldsymbol{HHHy}$.

For $|\boldsymbol{V}|$, $\text{tr}(\boldsymbol{V}^{-1})$ and $\text{tr}(\boldsymbol{V}^{-1}\boldsymbol{V}^{-1})$, it suffices to apply the spectral decomposition (Eq. (14)). We have

$$|\boldsymbol{V}| = \prod_{i=1}^{n}(\delta\xi_i + 1) \tag{19}$$

$$\text{tr}(\boldsymbol{V}^{-1}) = \sum_{i=1}^{n} \frac{1}{\delta\xi_i + 1} \tag{20}$$

$$\text{tr}(\boldsymbol{V}^{-1}\boldsymbol{V}^{-1}) = \sum_{i=1}^{n} \frac{1}{(\delta\xi_i + 1)^2} \tag{21}$$

which can all be evaluated in $\mathcal{O}(n)$ time.

For $\boldsymbol{y}'\boldsymbol{H}\boldsymbol{y}$, $\boldsymbol{y}'\boldsymbol{H}\boldsymbol{H}\boldsymbol{y}$ and $\boldsymbol{y}'\boldsymbol{H}\boldsymbol{H}\boldsymbol{H}\boldsymbol{y}$, recursive formulas are derived such that eventually we only need to evaluate the following form of quadratic forms: $\boldsymbol{a}'\boldsymbol{V}^{-1}\boldsymbol{b}$, $\boldsymbol{a}'\boldsymbol{V}^{-1}\boldsymbol{V}^{-1}\boldsymbol{b}$ and $\boldsymbol{a}'\boldsymbol{V}^{-1}\boldsymbol{V}^{-1}\boldsymbol{V}^{-1}\boldsymbol{b}$, where the vectors $\boldsymbol{a}$ and $\boldsymbol{b}$ are either $\boldsymbol{y}$ or a column of the matrix $\boldsymbol{W}$ (recall Eq. (2)). For this we again apply the spectral decomposition (Eq. (14) and make the linear transformations: $\tilde{\boldsymbol{a}} = \boldsymbol{U}'\boldsymbol{a}$, $\tilde{\boldsymbol{b}} = \boldsymbol{U}'\boldsymbol{b}$. Then we have

$$\boldsymbol{a}'\boldsymbol{V}^{-1}\boldsymbol{b} = \sum_{i=1}^{n} \frac{\tilde{\boldsymbol{a}}_i\tilde{\boldsymbol{b}}_i}{\delta\xi_i + 1} \tag{22}$$

$$\boldsymbol{a}'\boldsymbol{V}^{-1}\boldsymbol{V}^{-1}\boldsymbol{b} = \sum_{i=1}^{n} \frac{\tilde{\boldsymbol{a}}_i\tilde{\boldsymbol{b}}_i}{(\delta\xi_i + 1)^2} \tag{23}$$

$$\boldsymbol{a}'\boldsymbol{V}^{-1}\boldsymbol{V}^{-1}\boldsymbol{V}^{-1}\boldsymbol{b} = \sum_{i=1}^{n} \frac{\tilde{\boldsymbol{a}}_i\tilde{\boldsymbol{b}}_i}{(\delta\xi_i + 1)^3} \tag{24}$$

which can again be evaluated in $\mathcal{O}(n)$ time.

The purpose of including the second derivative of the likelihood function $LL(\delta)$ in the algorithm is that GEMMA uses a combination of Brent's method, which is more stable and Newton-Raphson algorithm which is more efficient for numerical optimization.

The overall time complexity for GEMMA is similar to FaST-LMM, namely $\mathcal{O}(n^3)+\mathcal{O}(pn^2)+\mathcal{O}(ptn)$.

2.3. **Grid-LMM.** The algorithm Grid-LMM [13] solves the LMM in a slightly different way from FaST-LMM and GEMMA. Instead of directly finding solutions for the unknown parameters by numerical optimization methods, it defines a grid spanning all valid values for the unknown parameters and evaluates the log-likelihood function at each grid location. Then, the optimized value for the unknown parameter is determined as the one corresponding to the largest value of the log-likelihood function across the grid.

The settings of the LMM (Eq. (2)) are slightly modified as follows: Let $\sigma^2 = \sigma_g^2 + \sigma_e^2$, $h^2 = \sigma_g^2/\sigma^2$, then we have $\boldsymbol{g} \sim \mathcal{N}(0, h^2\sigma^2\boldsymbol{K})$, $\boldsymbol{e} \sim \mathcal{N}(0, (1-h^2)\sigma^2\boldsymbol{I})$. We define $\boldsymbol{V} = h^2\boldsymbol{K} + (1-h^2)\boldsymbol{I}$ (Note that this $\boldsymbol{V}$ is not the same as the one defined in subsection 1.2. But they only differ by a scalar, namely $\sigma_e^2/\sigma^2$).

If the value of $h^2$ is predetermined, $\boldsymbol{V}$ is a positive-definite symmetric matrix. Hence we can apply Cholesky decomposition to it. Namely, there exists a lower-triangular matrix $\boldsymbol{L}$ such that $\boldsymbol{V} = \boldsymbol{L}\boldsymbol{L}'$. Making linear transformations $\boldsymbol{y}^* = \boldsymbol{L}^{-1}\boldsymbol{y}$ and $\boldsymbol{W}^* = \boldsymbol{L}^{-1}\boldsymbol{W}$. we can reformulate the log-likelihood function Eq. (3) as following:

$$LL(\alpha, \sigma^2|h^2) = -\frac{n}{2}\log(2\pi\sigma^2) - \log|\boldsymbol{L}| - \frac{1}{2\sigma^2}(\boldsymbol{y}^* - \boldsymbol{W}^*\boldsymbol{\alpha})'(\boldsymbol{y}^* - \boldsymbol{W}^*\boldsymbol{\alpha}) \tag{25}$$

Maximizing the function requires that its partial derivatives with respect to both $\alpha$ and $\sigma^2$ are zero. Thus, similar to Eq. (7), we obtain

$$LL(h^2) = -\frac{n}{2}\left(1 + \log(\frac{n}{2\pi}) + \log(\boldsymbol{y}^{*\prime}(\boldsymbol{I} - \boldsymbol{W}^*(\boldsymbol{W}^{*\prime}\boldsymbol{W}^*)^{-1}\boldsymbol{W}^{*\prime})\boldsymbol{y}^*)\right) - \log|\boldsymbol{L}| \tag{26}$$

So, at each grid vertex, the Cholesky decomposition of $\boldsymbol{V}$ has to be calculated once with time complexity $\frac{1}{6}\mathcal{O}(n^3) \sim \mathcal{O}(n^3)$. But the decomposition applies to all markers. The linear transformation with $\boldsymbol{L}^{-1}$ takes $\mathcal{O}(n^2)$ time for each marker. Finally, evaluating the log-likelihood function (Eq. (26)) takes only $\mathcal{O}(n)$ time. Therefore, considering the dominating part, the overall time complexity for Grid-LMM is approximately $\mathcal{O}(gn^3) + \mathcal{O}(gpn^2)$, where $g$ is the number of grid vertices.

Note that algorithms based on direct numerical optimization method such as FaST-LMM and GEMMA only requires to perform the spectral decomposition to an $n \times n$ matrix once, but Grid-LMM needs to perform the Chelosky decomposition $g$ times. Although calculating the Chelosky decomposition is much faster than calculating the spectral decomposition (but still takes $\mathcal{O}(n^3)$ time), if the number of grid vertices is too large, it may hamper the speed of Grid-LMM. However, the Grid-LMM algorithm can be easily generalized to solve LMM with multiple random terms with different covariance matrices, while algorithms based on spectral decomposition do not work because different matrices are usually not simultaneously diagonalizable.

2.4. **GAPIT-MLM.** In this algorithm, it is proposed that we do not maximize the log-likelihood function for each marker. Instead, we only maximize it for the null model, i.e. the full model (Eq. (1)) excluding the term of marker being tested:

$$\boldsymbol{y} = \boldsymbol{X\beta} + \boldsymbol{g} + \boldsymbol{e}, \tag{27}$$

Using the spectral decomposition as in FaST-LMM or GEMMA, this model can be solved with $\mathcal{O}(n^3) + \mathcal{O}(tn)$ time. Now, we get the estimates of the unknown variance components $\hat{\sigma}_g^2$ and $\hat{\sigma}_e^2$. The key idea is to fix them throughout the testing procedure, i.e. to use them as approximated values of the full model (Eq. (1)) for all markers. Thus, it is called "population parameters previously determined", abbreviated as"P3D" [19].

Then, we only need to produce the test statistics (Eq. (13)), which can be completed in $\mathcal{O}(pn)$ time provided that the linear transformation $\boldsymbol{U'm}$ has been done for all markers. Note that in FaST-LMM and GEMMA, it is done in the first step when evaluating the log-likelihood function for each marker. But in the case of P3D, it has not been done because the only evaluated log-likelihood function is for the null model. Taking this into account, the time complexity for calculating the test statistics for all markers is $\mathcal{O}(pn^2) + \mathcal{O}(pn) \sim \mathcal{O}(pn^2)$, and the total complexity of GAPIT-MLM is $\mathcal{O}(n^3) + \mathcal{O}(tn) + \mathcal{O}(pn^2)$.

Note that P3D has been implemented in many algorithms such as FaST-LMM (as an option in the old C++ version, and mandatory in the new Python version), EMMAX [6] and GCTA-MLMA [17]. These algorithms are very similar in terms of underlying mathematics. Thus, here we choose GAPIT-MLM as a representative. BOLT-LMM [11] and fastGWA [5] is also based on P3D, but they have implemented other techniques which will be introduced later.

2.5. **MLMA-LOCO.** The algorithm MLMA is implemented in the package GCTA. Compared with GAPIT-MLM, it implements an additional technique called "leave-one-chromosome-out" (LOCO). That is, when a marker is tested, all markers on the same chromosome are excluded in the calculation of kinship matrix. The purpose of LOCO is to avoid double fitting the candidate marker both as fixed effect and as random effect in the kinship matrix, as it was reported in the literature that including the candidate marker in the kinship matrix reduced the statistical power. This is the so-called "proximal contamination" phenomenon [9, 18].

Let $h$ be the number of chromosomes. For each $i \in \{1, 2, \ldots, h\}$, there is a specific null model:

$$\boldsymbol{y} = \boldsymbol{X\beta} + \boldsymbol{g}_i + \boldsymbol{e}_i, \tag{28}$$

where $\boldsymbol{g}_i \sim \mathcal{N}(0, \boldsymbol{K}_i\sigma_{g_i}^2)$, $\boldsymbol{e}_i \sim \mathcal{N}(0, \boldsymbol{I}\sigma_{e_i}^2)$, $\boldsymbol{K}_i$ is a modified kinship matrix calculated as $\boldsymbol{M}_i\boldsymbol{M}_i'$ (up to a scalar), and $\boldsymbol{M}_i$ is the matrix of all markers except those on the $i$-th chromosome. Other parameters are the same as in Eq. (27).

Then, the $h$ null models were solved and unknown parameters are estimated. For markers on the $i$-th chromosome, the test statistics are calculated by Eq. (13) using the estimates of variance components $\hat{\sigma}_{g_i}^2$, $\hat{\sigma}_{e_i}^2$ and the modified kinship matrix $\boldsymbol{K}_i$.

2.6. **BOLT-LMM.** The main contribution of this algorithm in efficiency is that it applies a series of mathematical techniques to reduce the overall time complexity to $\mathcal{O}(pn)$, up to a certain number of iterations and Monte Carlo samplings, which is a great improvement as the time complexity is linear with respect to the population size $n$. Another important feature is that in addition to the standard assumption of the Q+K model, it provides an option to assume non-infinitesimal genetic architecture via Bayesian methods. Some details are discussed below.

2.6.1. *REML.* BOLT-LMM uses REML instead of ML to estimate the variance components of the model and it adopted the "P3D" method. So the variance components are only estimated once for the null model (Eq. (27)).

First, the null model is re-written as the following

$$\boldsymbol{y} = \boldsymbol{X}\boldsymbol{\beta} + \boldsymbol{M}\boldsymbol{u} + \boldsymbol{e}. \tag{29}$$

That is, $\boldsymbol{g}$ is replaced by $\boldsymbol{M}\boldsymbol{u}$. Recall that $\boldsymbol{M}$ is the $n \times p$ matrix of all markers. Here, $\boldsymbol{u}$ is a random vector of all marker effects, $\boldsymbol{u} \sim \mathcal{N}(0, \boldsymbol{I}\sigma_u^2)$. This means that the polygenic effects $\boldsymbol{g}$ are assumed to be a linear combination of all marker effects. Assuming $\sigma_u^2 = \sigma_g^2/c$, we see that Eq. (29) is equivalent to Eq. (27), since $\boldsymbol{K} = \boldsymbol{M}\boldsymbol{M}'/c$ (see Section 1.1).

Next, the fixed effects term are annihilated by left-multiplying a projection matrix $\boldsymbol{R}$ to both sides of (29) such that $\boldsymbol{R}\boldsymbol{X} = 0$. Then, Eq. (29) becomes

$$\tilde{\boldsymbol{y}} = \tilde{\boldsymbol{M}}\boldsymbol{u} + \tilde{\boldsymbol{e}}, \tag{30}$$

where $\tilde{\boldsymbol{y}} = \boldsymbol{R}\boldsymbol{y}$, $\tilde{\boldsymbol{M}} = \boldsymbol{R}\boldsymbol{M}$ and $\tilde{\boldsymbol{e}} = \boldsymbol{R}\boldsymbol{e}$. Note that $\boldsymbol{R}^2 = \boldsymbol{R}$ since it is a projection matrix. Hence $\tilde{\boldsymbol{e}} \sim \mathcal{N}(0, \boldsymbol{R}\sigma_e^2)$. The assumption for $\boldsymbol{u}$ does not change. Let $\tilde{\boldsymbol{V}}_0 = \mathrm{var}(\tilde{\boldsymbol{y}}) = \tilde{\boldsymbol{M}}\tilde{\boldsymbol{M}}'\sigma_u^2 + \boldsymbol{R}\sigma_e^2$.

Now, the log-likelihood function has the following form:

$$LL_{re}(\sigma_u^2, \sigma_e^2) = -\frac{n}{2}\log(2\pi) - \frac{1}{2}|\tilde{\boldsymbol{V}}_0| - \frac{1}{2}\tilde{\boldsymbol{y}}'\tilde{\boldsymbol{V}}_0^{-1}\tilde{\boldsymbol{y}}. \tag{31}$$

It seems that the above expression depends on the choice of the projection matrix $\boldsymbol{R}$, which is not unique. But it can be proved that actually it is independent of the choice of $\boldsymbol{R}$ [12].

As before, we need to find $\hat{\sigma}_u^2$ and $\hat{\sigma}_e^2$ which maximize the log-likelihood function. For this, it is required that the partial derivatives of the log-likelihood function with respect to $\sigma_u$ and $\sigma_e$ are zero. This yields the following

$$\mathrm{tr}(\tilde{\boldsymbol{V}}_0^{-1}\tilde{\boldsymbol{M}}\tilde{\boldsymbol{M}}') = \tilde{\boldsymbol{y}}'\tilde{\boldsymbol{V}}_0^{-1}\tilde{\boldsymbol{M}}\tilde{\boldsymbol{M}}'\tilde{\boldsymbol{V}}_0^{-1}\tilde{\boldsymbol{y}} \tag{32}$$

$$\mathrm{tr}(\tilde{\boldsymbol{V}}_0^{-1}\boldsymbol{R}) = \tilde{\boldsymbol{y}}'\tilde{\boldsymbol{V}}_0^{-1}\boldsymbol{R}\tilde{\boldsymbol{V}}_0^{-1}\tilde{\boldsymbol{y}} \tag{33}$$

2.6.2. *Efficiently solving the LMM.* The method used by BOLT-LMM to solve $\hat{\sigma}_u^2$ and $\hat{\sigma}_e^2$ is different from previous algorithms. It is based on an observation that Eq. (32) and (33) are equivalent to

$$E(\hat{\boldsymbol{u}}'_{\mathrm{rand}}\hat{\boldsymbol{u}}_{\mathrm{rand}}) = \hat{\boldsymbol{u}}'\hat{\boldsymbol{u}} \tag{34}$$

$$E(\hat{\boldsymbol{e}}'_{\mathrm{rand}}\hat{\boldsymbol{e}}_{\mathrm{rand}}) = \hat{\boldsymbol{e}}'\hat{\boldsymbol{e}} \tag{35}$$

where $\hat{\boldsymbol{u}}$ and $\hat{\boldsymbol{e}}$ are the best linear unbiased predictions (BLUPs) for $\boldsymbol{u}$ and $\boldsymbol{e}$ based on the observed (and transformed) phenotypic data $\tilde{\boldsymbol{y}}$, whereas $\hat{\boldsymbol{u}}_{\mathrm{rand}}$ and $\hat{\boldsymbol{e}}_{\mathrm{rand}}$ are BLUPs for $\boldsymbol{u}$ and $\boldsymbol{e}$ based on randomly generated phenotypic data $\tilde{\boldsymbol{y}}_{\mathrm{rand}}$ according to Eq. (30), i.e. $\tilde{\boldsymbol{y}}_{\mathrm{rand}} \sim \mathcal{N}(0, \tilde{\boldsymbol{V}}_0)$. The proof can be found in [12].

This result enables us to solve $\hat{\sigma}_u^2$ and $\hat{\sigma}_e^2$ by Monte-Carlo sampling. Namely, given $\sigma_u^2$ and $\sigma_e^2$, we can randomly sample phenotypic data $\tilde{\boldsymbol{y}}_{\mathrm{rand}}$, calculate the expectations $E(\hat{\boldsymbol{u}}'_{\mathrm{rand}}\hat{\boldsymbol{u}}_{\mathrm{rand}})$, $E(\hat{\boldsymbol{e}}'_{\mathrm{rand}}\hat{\boldsymbol{e}}_{\mathrm{rand}})$ and compare them with the BLUPs $\hat{\boldsymbol{u}}$, $\hat{\boldsymbol{e}}$ based on the observed data. Based on this result, BOLT-LMM implemented an efficient algorithm to find out $\hat{\sigma}_u^2$ and $\hat{\sigma}_e^2$. We briefly sketch the essential steps as follows:

Recall that $\sigma_u^2 = \sigma_g^2/c$. Let $\delta = \sigma_g^2/\sigma_e^2$, then $\tilde{\boldsymbol{V}}_0 = (\delta\tilde{\boldsymbol{M}}\tilde{\boldsymbol{M}}'/c + \boldsymbol{R})\sigma_e^2$. For convenience, we define $\tilde{\boldsymbol{V}} = \delta\tilde{\boldsymbol{M}}\tilde{\boldsymbol{M}}'/c + \boldsymbol{R}$, i.e. $\tilde{\boldsymbol{V}}_0 = \tilde{\boldsymbol{V}}\sigma_e^2$ (Note that this notation is consistent with $\boldsymbol{V}$ defined in Section

1.2). From [3], we know that the BLUPs for $\boldsymbol{u}$ and $\boldsymbol{e}$ are:

$$\hat{\boldsymbol{u}} = \frac{\delta}{c}\tilde{\boldsymbol{M}}'\tilde{\boldsymbol{V}}^{-1}\tilde{\boldsymbol{y}} \tag{36}$$

$$\hat{\boldsymbol{e}} = \tilde{\boldsymbol{V}}^{-1}\tilde{\boldsymbol{y}} \tag{37}$$

The above equations indicate that $\hat{\boldsymbol{u}}$ and $\hat{\boldsymbol{e}}$ only depend on the variance ratio $\delta$, not on the scale of $\hat{\sigma}_u^2$ and $\hat{\sigma}_e^2$. Therefore, finding $\hat{\sigma}_u^2$ and $\hat{\sigma}_e^2$ satisfying Eq. (34) and (35) is equivalent to finding $\hat{\delta}$ satisfying

$$\frac{E(\hat{\boldsymbol{u}}_{\text{rand}}'\hat{\boldsymbol{u}}_{\text{rand}})}{E(\hat{\boldsymbol{e}}_{\text{rand}}'\hat{\boldsymbol{e}}_{\text{rand}})} = \frac{\hat{\boldsymbol{u}}'\hat{\boldsymbol{u}}}{\hat{\boldsymbol{e}}'\hat{\boldsymbol{e}}} \tag{38}$$

and choosing $\hat{\sigma}_u^2$ to scale the expectations on the left side to match the values on the right side of Eq. (34) or (35).

Then, we can define the following function

$$f(\delta) = \log\left(\frac{E(\hat{\boldsymbol{u}}_{\text{rand}}'\hat{\boldsymbol{u}}_{\text{rand}})}{E(\hat{\boldsymbol{e}}_{\text{rand}}'\hat{\boldsymbol{e}}_{\text{rand}})} \bigg/ \frac{\hat{\boldsymbol{u}}'\hat{\boldsymbol{u}}}{\hat{\boldsymbol{e}}'\hat{\boldsymbol{e}}}\right). \tag{39}$$

And we can see that requiring $\hat{\delta}$ to satisfy Eq. (38) is equivalent to saying that $\hat{\delta}$ is a zero of $f(\delta)$. Thus, we could apply classic numerical methods such as the secant method to find out $\hat{\delta}$.

Now, we discuss the time complexity of the above procedure. To evaluate the function $f(\delta)$, we need to first generate a set of random phenotypes $\tilde{\boldsymbol{y}}$. This can be done by first randomly generating the following components:

$$\boldsymbol{u}_{\text{rand}} \sim \mathcal{N}(0, 1/c) \tag{40}$$

$$\boldsymbol{e}_{\text{rand,unscaled}} \sim \mathcal{N}(0, 1) \tag{41}$$

$$\boldsymbol{e}_{\text{rand,unscaled,proj}} = \boldsymbol{R}\boldsymbol{e}_{\text{rand,unscaled}} \tag{42}$$

Then, the random phenotypes are generated as follows:

$$\tilde{\boldsymbol{y}}_{\text{rand}} = \tilde{\boldsymbol{M}}\boldsymbol{u}_{\text{rand}} + \boldsymbol{e}_{\text{rand,unscaled,proj}}/\sqrt{\delta} \tag{43}$$

We can check that $\text{var}(\tilde{\boldsymbol{y}}_{\text{rand}}) = \tilde{\boldsymbol{V}}$. This procedure involves mainly matrix-vector multiplications, in which the dominant part is the product of $n \times p$ matrix $\tilde{\boldsymbol{M}}$ and the $p$-dimensional vector $\boldsymbol{u}_{\text{rand}}$. So the time complexity is $\mathcal{O}(spn)$, where $s$ is the total number of sampling.

Seconly, we need to calculate the BLUPs $\hat{\boldsymbol{u}}_{\text{rand}}$, $\hat{\boldsymbol{e}}_{\text{rand}}$, $\hat{\boldsymbol{u}}$ and $\hat{\boldsymbol{e}}$ using Eq. (36) and (37). If we calculate directly, the most time consuming part is $\tilde{\boldsymbol{V}}^{-1}$ because it requires $\mathcal{O}(n^2p)$ time to calculate the product $\tilde{\boldsymbol{M}}\tilde{\boldsymbol{M}}'$ and $\mathcal{O}(n^3)$ time to take the matrix inverse. However, all we need is to calculate the matrix-vector product $\tilde{\boldsymbol{V}}^{-1}\tilde{\boldsymbol{y}}$. This is equivalent to the problem of solving systems of linear equations $\tilde{\boldsymbol{V}}\boldsymbol{x} = \tilde{\boldsymbol{y}}$ and hence equivalent to the problem of finding $\boldsymbol{x}$ maximizing the function $\phi(\boldsymbol{x}) = -\boldsymbol{x}'\tilde{\boldsymbol{V}}\boldsymbol{x} + 2\boldsymbol{x}'\tilde{\boldsymbol{y}}$. Then, BOLT-LMM applied the conjugate gradient method [2], which is a well-known numerical optimization method. In each iteration (say, the $k$-th one), we only need to calculate a single product of the matrix $\tilde{\boldsymbol{V}}$ and a certain vector $\boldsymbol{s}_k$, as well as several inner products of vectors. In particular, we do not explicitly calculate the matrix product $\tilde{\boldsymbol{M}}\tilde{\boldsymbol{M}}'$, but only need to calculate $\boldsymbol{R}\boldsymbol{s}_k$, $\tilde{\boldsymbol{M}}'\boldsymbol{s}_k$ and $\tilde{\boldsymbol{M}}(\tilde{\boldsymbol{M}}'\boldsymbol{s}_k)$, each being a matrix-vector product. Thus, the time complexity is $\mathcal{O}(stn^2) + \mathcal{O}(stpn)$, where $t$ is the average number of iterations. In addition, the projection matrix $\boldsymbol{R}$ can be replaced by the identity matrix $\boldsymbol{I}$ [11], which further reduces the time complexity to $\mathcal{O}(stpn)$.

2.6.3. *Producing the test statistics.* In order to calculate the test statistics efficiently, BOLT-LMM introduced certain approximations to the standard test statistic. Recall that the full model including the marker effect is the following:

$$\boldsymbol{y} = \boldsymbol{X}\boldsymbol{\beta} + \boldsymbol{m}a + \boldsymbol{M}\boldsymbol{u} + \boldsymbol{e}. \tag{44}$$

Note that this model is equivalent to Eq. (1) because the corresponding null models (27) and (29) are equivalent. Thus the test statistic is the same as in Eq. (13).

Then, the model is transformed by the projection matrix $\boldsymbol{R}$, which yields the following

$$\tilde{\boldsymbol{y}} = \tilde{\boldsymbol{m}}\boldsymbol{a} + \tilde{\boldsymbol{M}}\boldsymbol{u} + \tilde{\boldsymbol{e}}, \tag{45}$$

where $\tilde{\boldsymbol{m}} = \boldsymbol{R}\boldsymbol{m}$ the transformed vector of marker profiles (Note that it is very important that the transformation is also applied to $\boldsymbol{m}$), all other notations are the same as in Eq. (30). Following the arguments in Section 1.4, the corresponding Wald-test statistic is

$$T_{Wald,re} = \frac{1}{\hat{\sigma}_e^2} \cdot \frac{(\tilde{\boldsymbol{m}}'\tilde{\boldsymbol{V}}^{-1}\tilde{\boldsymbol{y}})^2}{\tilde{\boldsymbol{m}}'\tilde{\boldsymbol{V}}^{-1}\tilde{\boldsymbol{m}}} = \frac{(\tilde{\boldsymbol{m}}'\tilde{\boldsymbol{V}}_0^{-1}\tilde{\boldsymbol{y}})^2}{\tilde{\boldsymbol{m}}'\tilde{\boldsymbol{V}}_0^{-1}\tilde{\boldsymbol{m}}} \tag{46}$$

An important result is that the two test statistics $T_{wald}$ and $T_{wald,re}$ are the same, which means that the test statistic is invariant after making the linear transformation by the projection matrix.

The BOLT-LMM-inf statistic is obtained by replacing the term $\tilde{\boldsymbol{m}}'\tilde{\boldsymbol{V}}_0^{-1}\tilde{\boldsymbol{m}}$ in Eq. (46) by a constant, namely

$$T_{\text{BOLT-LMM-inf}} = \frac{(\tilde{\boldsymbol{m}}'\tilde{\boldsymbol{V}}_0^{-1}\tilde{\boldsymbol{y}})^2}{c_{\text{inf}}}, \tag{47}$$

where

$$c_{\text{inf}} = \frac{\text{mean}(\tilde{\boldsymbol{m}}'\tilde{\boldsymbol{V}}_0^{-1}\tilde{\boldsymbol{y}})^2}{\text{mean}(T_{\text{BOLT-LMM-inf}})}. \tag{48}$$

And practically, the two means in the above equation is calculated by randomly taking 30 markers not significantly associated with the phenotype [11]. Thus, the calculation of $c_{\text{inf}}$ is efficient.

In this way, the BOLT-LMM-inf test statistic avoids calculating $\tilde{\boldsymbol{m}}'\tilde{\boldsymbol{V}}_0^{-1}\tilde{\boldsymbol{m}}$ for each marker, which would take $\mathcal{O}(pn^2)$ time. Evaluating $\tilde{\boldsymbol{m}}'\tilde{\boldsymbol{V}}_0^{-1}\tilde{\boldsymbol{y}}$ for each marker only takes $\mathcal{O}(pn)$ time, since one can first calculate the vector $\tilde{\boldsymbol{V}}_0^{-1}\tilde{\boldsymbol{y}}$ and then what remains are just inner products of vectors.

2.6.4. *Incorporating Gaussian mixture priors to the model.* Similar to Bayesian genomic prediction models, BOLT-LMM incorporates Gaussian mixture priors for the marker effects $\boldsymbol{u} = (u_1, u_2, \ldots, u_p)'$ in the null model (30) in order to take non-infinitesimal genetic architecture into account. More precisely, the basic equation of the model is the same, but we assume that for any $i \in \{1, 2, \ldots, p\}$,

$$u_i \sim \begin{cases} \mathcal{N}(0, \sigma_{u,1}^2), & \text{with probability } \pi \\ \mathcal{N}(0, \sigma_{u,2}^2), & \text{with probability } 1-\pi \end{cases} \tag{49}$$

and the total variance of the mixture for all markers was set to equal $\hat{\sigma}_u^2$ estimated in the infinitesimal model (Eq. (30)). BOLT-LMM defines a small set of values for the unknown parameters (namely $\sigma_{u,1}^2$, $\sigma_{u,1}^2$ and $\pi$), and performs cross-validation to determine the parameters which optimize the mean-squared prediction ability. The Bayesian linear model is fitted by a variational approximation approach. For details, see the supplementary notes in [11]. The resulting best Gaussian mixture model was compared with the infinitesimal model. If its prediction ability is not significantly higher than the infinitesimal model, the procedure stops and the BOLT-LMM-inf test statistics are the final results. Otherwise, new test statistics are calculated as follows.

$$T_{\text{BOLT-LMM-mix}} = \frac{(\tilde{\boldsymbol{m}}'\hat{\boldsymbol{e}}_{\text{Gaussian}})^2}{c_{\text{LD}}}, \tag{50}$$

where $\hat{\boldsymbol{e}}_{\text{Gaussian}}$ is the residual vector of the non-infinitesimal null model and $c_{\text{LD}}$ is a constant calculated via LD-score regression [1], for which the details are beyond the scope of this note.

2.7. **fastGWA.** The algorithm fastGWA [5] implemented in the software package GCTA [17] provides an option to put a threshold to the entries in the relationship matrix $\boldsymbol{K}$ (the default value is 0.05). All entries below the threshold are set to zero. In this way, $\boldsymbol{K}$ may become a sparse matrix, and then $\boldsymbol{V} = \delta\boldsymbol{K} + \boldsymbol{I}$ is also sparse. Many efficient algorithms are available for sparse matrices and have been implemented as libraries in various programming environments. In particular, the matrix inverse and the Chelosky decomposition of sparse matrices can be much more efficiently calculated than dense matrices. Together with a grid search strategy similar to Grid-LMM [13], the LMM is solved efficiently. Note that fastGWA also follows P3D, so the variance components are estimated only once for the null model. When the matrix $\boldsymbol{K}$ is still dense, fastGWA implemented the conjugate gradient method (see Section 2.6.2) to avoid explicitly calculating the inverse of $\boldsymbol{V}$.

Similar to BOLT-LMM, fastGWA also implements an approximated approach to reduce the computational complexity in calculating test statistics, namely the GRAMMAR-Gamma approximation [14]. Recall that the Wald-test statistic is as Eq. (46), which can be re-written as

$$T_{\text{Wald,re}} = \frac{(\tilde{\boldsymbol{m}}'\tilde{\boldsymbol{V}}_0^{-1}\tilde{\boldsymbol{y}})^2}{\tilde{\boldsymbol{m}}'\tilde{\boldsymbol{V}}_0^{-1}\tilde{\boldsymbol{m}}} = \frac{(\tilde{\boldsymbol{m}}'\tilde{\boldsymbol{V}}_0^{-1}\tilde{\boldsymbol{y}})^2}{\tilde{\boldsymbol{m}}'\tilde{\boldsymbol{m}}} \Bigg/ \frac{\tilde{\boldsymbol{m}}'\tilde{\boldsymbol{V}}_0^{-1}\tilde{\boldsymbol{m}}}{\tilde{\boldsymbol{m}}'\tilde{\boldsymbol{m}}} \tag{51}$$

GRAMMAR-Gamma proposed to replace the denominator in Eq. (51) by its mean value over all markers, i.e.

$$T_{\text{GRAMMAR-Gamma}} = \frac{(\tilde{\boldsymbol{m}}'\tilde{\boldsymbol{V}}_0^{-1}\tilde{\boldsymbol{y}})^2}{\tilde{\boldsymbol{m}}'\tilde{\boldsymbol{m}}} \Bigg/ \frac{1}{p}\sum_{i=1}^{p}\frac{\tilde{\boldsymbol{m}}_i'\tilde{\boldsymbol{V}}_0^{-1}\tilde{\boldsymbol{m}}_i}{\tilde{\boldsymbol{m}}_i'\tilde{\boldsymbol{m}}_i} \tag{52}$$

As explained in Section 2.6.3, calculating $\tilde{\boldsymbol{m}}'\tilde{\boldsymbol{V}}_0^{-1}\tilde{\boldsymbol{y}}$ for all markers requires only $\mathcal{O}(np)$ time, and the same for evaluating $\tilde{\boldsymbol{m}}'\tilde{\boldsymbol{m}}$. It was shown that $\gamma = \frac{1}{p}\sum_{i=1}^{p}\frac{\tilde{\boldsymbol{m}}_i'\tilde{\boldsymbol{V}}_0^{-1}\tilde{\boldsymbol{m}}_i}{\tilde{\boldsymbol{m}}_i'\tilde{\boldsymbol{m}}_i}$ can be directly calculated from the trace of $b\tilde{V}_0^{-1}$ and the estimated variance components [14]. Hence, the total time complexity for calculating $T_{\text{GRAMMAR-Gamma}}$ for all markers is only $\mathcal{O}(pn)$.

In fastGWA, the value $\gamma$ is approximated by $\hat{\gamma}$ which is the mean value of $\frac{\tilde{\boldsymbol{m}}_i'\tilde{\boldsymbol{V}}_0^{-1}\tilde{\boldsymbol{m}}_i}{\tilde{\boldsymbol{m}}_i'\tilde{\boldsymbol{m}}_i}$ over $1,000$ randomly chosen markers not associated with the phenotype.

2.8. **GAPIT-CMLM.** The idea of CMLM [19] is to reduce the dimension of the term $\boldsymbol{g}$ in the Q+K model such that the computational load of decomposing the matrix $\boldsymbol{K}$ decreases. More precisely, it applies a clustering algorithm to group the $n$ individuals into $l$ groups and replace the $n \times n$ kinship matrix $\boldsymbol{K}$ by a compressed kinship matrix $\boldsymbol{K}_c$ of size $l \times l$. The model is of the following form.

$$\boldsymbol{y} = \boldsymbol{X}\boldsymbol{\beta} + \boldsymbol{m}a + \boldsymbol{Z}\boldsymbol{r} + \boldsymbol{e}, \tag{53}$$

where $\boldsymbol{r}$ is an $l$-dimensional vector of the genetic effects for the $l$ groups, $\boldsymbol{Z}$ is the $n \times l$ incidence matrix allocating the $n$ individuals into the corresponding groups. All other notations are the same as in (1). $\boldsymbol{r}$ is assumed to be a random vector, $\boldsymbol{r} \sim \mathcal{N}(0, \boldsymbol{K}_c\sigma_r^2)$.

Assuming that the $l$ groups are denoted by $G_1, G_2, \ldots, G_l$. For any $i, j \in \{1, 2, \ldots, l\}$, the $(i, j)$-entry in $\boldsymbol{K}_c$ is determined as the mean of individual-based kinship coefficients between the group $G_i$ and $G_j$. That is

$$(\boldsymbol{K}_c)_{G_iG_j} = \frac{1}{|G_i||G_j|}\sum_{x\in G_i}\sum_{y\in G_j} k_{xy} \tag{54}$$

where $|G_i|$ ($|G_j|$) is the number of individuals in group $G_i$ ($G_j$), $k_{xy}$ is the $(x, y)$-entry in the original kinship matrix $\boldsymbol{K}$.

There is also an improved version ECMLM [7] aiming at increasing the statistical power but at the cost of increase the computational load. Briefly, ECMLM implemented three different ways of compressing the kinship coefficients (taking the mean as in (54), and replacing the mean by the median or the maximum) and eight different approaches of clustering. The combination which gives the best model fit is finally used to producing the test statistic. Note that P3D is implemented in

CMLM and ECMLM, thus after the clustering and compressing procedure, the LMM is solved only once for the null model.

2.9. **FarmCPU.** With the aim of increasing the statistical power, keeing control of false-positive rate, and meanwhile reducing the computational load, the framework of FarmCPU [10] is slightly different from the original Q+K LMM (Eq. (1)). Similar to the idea of FaST-LMM-select, it does not use all markers to generate the kinship matrix $\boldsymbol{K}$, but implements an algorithm to select marker covariates which were termed pseudo QTNs for constructing $\boldsymbol{K}$. Once the covariates are selected, all markers are tested one-by-one in a multiple linear regression model including the selected covariates. The procedure is iterated until the set of covariates is stable. Some details are sketched below.

The multiple linear regression model used for testing marker effects is as follows:

$$\boldsymbol{y} = \boldsymbol{X}\boldsymbol{\beta} + \boldsymbol{M}_S\boldsymbol{b} + m_i\boldsymbol{a}_i + \boldsymbol{e}. \tag{55}$$

All notations are the same as in Eq. (1) except for the term $\boldsymbol{M}_s\boldsymbol{b}$, in which $\boldsymbol{b} = (b_{i_1}, b_{i_2}, \ldots, b_{i_s})'$ is the vector of marker covariate effects and $\boldsymbol{M}_s$ is the $n \times s$ matrix of the corresponding marker profiles. Here $S = \{i_1, i_2, \ldots, i_s\} \subset \{1, 2, \ldots, p\}$ is the subset of marker indices chosen as covariates. This term replaces the random vector $\boldsymbol{g}$ in (1) and serves as a control of population structure.

The LMM used for selecting marker covariates is the following:

$$\boldsymbol{y} = \boldsymbol{X}\boldsymbol{\beta} + \boldsymbol{g} + \boldsymbol{e}, \tag{56}$$

where $\boldsymbol{y}$, $\boldsymbol{e}$ is the same as before, and similar to Eq. (1), $\boldsymbol{g} \sim \mathcal{N}(0, \boldsymbol{K}_s\sigma_g^2)$, $\boldsymbol{e} \sim \mathcal{N}(0, \boldsymbol{I}\sigma_e^2)$. Here $\boldsymbol{K}_s$ is a kinship matrix derived from $s$ selected markers.

The full procedure of FarmCPU can be described as follows: 1) For each $i \in \{1, 2, \ldots, p\} \setminus S$, testing the effect $\boldsymbol{a}_i$ using model (55). Thus, in total $p - s$ markers are tested and the model 55 is fitted $p - s$ times. 2) Meanwhile, each of the $s$ marker covariates get $p-s$ p-values and the final p-value of a marker covariate is determined as the lowest one among the $p - s$ values. This step is called substitution. Note that in the first iteration, $S = \emptyset$ as covariates have not been selected and the substitution step is skipped. 3) Selecting marker covariates based on model (56). More precisely, the genome is divided into bins each represented by the marker with lowest p-values in the bin. Then, the $s$ bins whose representative markers have the lowest p values are selected as covariates. The size of each bin and the number of selected bins $s$ is determined by maximizing the log-likelihood of (56). This step is adopted from the algorithm SUPER [15]. 4) Updating $S$ and go to 1). If $S$ does not change (which means that no new covariates can be identified), stop the iteration.

2.10. **BLINK.** The algorithm BLINK [4] can be considered as an improved version of FarmCPU. It has a similar iterative framework of selecting marker covariates and testing marker effects. The model for testing marker effects is the same as in FarmCPU, i.e. model (55). But BLINK replace the LMM model for selecting marker covariates by another multiple linear regression model. In this way, it improves the computational efficiency further as it does not use LMM at all.

The model used in BLINK for selecting marker covariates is the following.

$$\boldsymbol{y} = \boldsymbol{X}\boldsymbol{\beta} + \boldsymbol{M}_S\boldsymbol{b} + \boldsymbol{e}, \tag{57}$$

where the notations are the same as in (55). In fact, it is the "null model" of (55).

The procedure of BLINK is also similar to FarmCPU. In step 3) of FarmCPU, the bin size and the number of bins whose representative maker are selected as covariates are determined by maximizing the likelihood function of (56). In BLINK, the two parameters are determined by maximizing the Bayesian Information Content (BIC) of model (57).

#### References

1. Brendan K Bulik-Sullivan, Po-Ru Loh, Hilary K Finucane, Stephan Ripke, Jian Yang, Nick Patterson, Mark J Daly, Alkes L Price, and Benjamin M Neale, *Ld score regression distinguishes confounding from polygenicity in genome-wide association studies*, Nature genetics **47** (2015), no. 3, 291–295.
2. Michael T Heath, *Scientific computing: an introductory survey, revised second edition*, SIAM, 2018.

3. Charles R Henderson, *Best linear unbiased estimation and prediction under a selection model*, Biometrics (1975), 423–447.
4. Meng Huang, Xiaolei Liu, Yao Zhou, Ryan M Summers, and Zhiwu Zhang, *Blink: a package for the next level of genome-wide association studies with both individuals and markers in the millions*, Gigascience **8** (2019), no. 2, giy154.
5. Longda Jiang, Zhili Zheng, Ting Qi, Kathryn E Kemper, Naomi R Wray, Peter M Visscher, and Jian Yang, *A resource-efficient tool for mixed model association analysis of large-scale data*, Nature genetics **51** (2019), no. 12, 1749–1755.
6. Hyun Min Kang, Jae Hoon Sul, Susan K Service, Noah A Zaitlen, Sit-yee Kong, Nelson B Freimer, Chiara Sabatti, and Eleazar Eskin, *Variance component model to account for sample structure in genome-wide association studies*, Nature genetics **42** (2010), no. 4, 348–354.
7. Meng Li, Xiaolei Liu, Peter Bradbury, Jianming Yu, Yuan-Ming Zhang, Rory J Todhunter, Edward S Buckler, and Zhiwu Zhang, *Enrichment of statistical power for genome-wide association studies*, BMC biology **12** (2014), no. 1, 1–10.
8. Christoph Lippert, Jennifer Listgarten, Ying Liu, Carl M Kadie, Robert I Davidson, and David Heckerman, *Fast linear mixed models for genome-wide association studies*, Nature methods **8** (2011), no. 10, 833–835.
9. Jennifer Listgarten, Christoph Lippert, Carl M Kadie, Robert I Davidson, Eleazar Eskin, and David Heckerman, *Improved linear mixed models for genome-wide association studies*, Nature methods **9** (2012), no. 6, 525–526.
10. Xiaolei Liu, Meng Huang, Bin Fan, Edward S Buckler, and Zhiwu Zhang, *Iterative usage of fixed and random effect models for powerful and efficient genome-wide association studies*, PLoS genetics **12** (2016), no. 2, e1005767.
11. Po-Ru Loh, George Tucker, Brendan K Bulik-Sullivan, Bjarni J Vilhjalmsson, Hilary K Finucane, Rany M Salem, Daniel I Chasman, Paul M Ridker, Benjamin M Neale, Bonnie Berger, et al., *Efficient bayesian mixed-model analysis increases association power in large cohorts*, Nature genetics **47** (2015), no. 3, 284–290.
12. Charles E McCulloch and Shayle R Searle, *Generalized, linear, and mixed models*, John Wiley & Sons, 2004.
13. Daniel E Runcie and Lorin Crawford, *Fast and flexible linear mixed models for genome-wide genetics*, PLoS genetics **15** (2019), no. 2, e1007978.
14. Gulnara R Svishcheva, Tatiana I Axenovich, Nadezhda M Belonogova, Cornelia M Van Duijn, and Yurii S Aulchenko, *Rapid variance components–based method for whole-genome association analysis*, Nature genetics **44** (2012), no. 10, 1166–1170.
15. Qishan Wang, Feng Tian, Yuchun Pan, Edward S Buckler, and Zhiwu Zhang, *A super powerful method for genome wide association study*, PloS one **9** (2014), no. 9, e107684.
16. Christian Widmer, Christoph Lippert, Omer Weissbrod, Nicolo Fusi, Carl Kadie, Robert Davidson, Jennifer Listgarten, and David Heckerman, *Further improvements to linear mixed models for genome-wide association studies*, Scientific reports **4** (2014), no. 1, 1–13.
17. Jian Yang, S Hong Lee, Michael E Goddard, and Peter M Visscher, *Gcta: a tool for genome-wide complex trait analysis*, The American Journal of Human Genetics **88** (2011), no. 1, 76–82.
18. Jian Yang, Noah A Zaitlen, Michael E Goddard, Peter M Visscher, and Alkes L Price, *Advantages and pitfalls in the application of mixed-model association methods*, Nature genetics **46** (2014), no. 2, 100–106.
19. Zhiwu Zhang, Elhan Ersoz, Chao-Qiang Lai, Rory J Todhunter, Hemant K Tiwari, Michael A Gore, Peter J Bradbury, Jianming Yu, Donna K Arnett, Jose M Ordovas, et al., *Mixed linear model approach adapted for genome-wide association studies*, Nature genetics **42** (2010), no. 4, 355–360.
20. Xiang Zhou and Matthew Stephens, *Genome-wide efficient mixed-model analysis for association studies*, Nature genetics **44** (2012), no. 7, 821–824.

#### Supplementary Note B

In this supplementary note, we discussed some details on fastGWA-sp and fastGWA-GG. Recall that fastGWA-sp is based on the sparse kinship matrix but GRAMMAR-Gamma approximation is not applied, whereas fastGWA-GG implemented the original kinship matrix as well as the GRAMMAR-Gamma approximation. For comparison, it is necessary to consider the original algorithm of fastGWA, i.e. the algorithm based on the original kinship matrix and without the GRAMMAR-Gamma approximation, denoted by fastGWA-ori in this note.

**B.1 On the performance of fastGWA-GG**

In the main text, we reported that fastGWA-GG produced the lowest statistical power among the 12 selected algorithms in the benchmarking analysis. We would like to find out the reason. In particular, we may ask whether the reduced power was due to the implementation of the GRAMMAR-Gamma approximation.

To answer this question, we only need to compare the performance of fastGWA-GG with fastGWA-ori because they differ only in whether the GRAMMAR-Gamma approximation is implemented or not. However, fastGWA-ori was not evaluated in our benchmarking analysis, because the underlying model and key techniques implemented in this algorithm were already represented by other algorithms. More precisely, fastGWA-ori was based on the standard Q+K linear mixed model (represented by GEMMA), it implemented the P3D approximation (represented by GAPIT-MLM) and a grid-search approach similar to Grid-LMM. Since the performance of the three algorithms (GEMMA, Grid-LMM and GAPIT-MLM) were quite similar in the benchmarking analysis (see the main text), it seems that we could expect fastGWA-ori to produce similar power and false-positive rate as GEMMA/Grid-LMM/GAPIT-MLM. If this is true, we can conclude that the low power of fastGWA-GG was indeed due to the GRAMMAR-Gamma approximation.

However, we noticed that the underlying model of fastGWA-ori was slightly different from GEMMA/Grid-LMM/GAPIT-MLM, and their default methods of calculating the kinship matrix were also different. In the following we first provided a detail description on the differences. Then, we compared the performance of fastGWA-ori and GEMMA using simulated data sets. Finally, we drew conclusions on the possible source of the low power for fastGWA-GG.

***Comparing the underlying model of fastGWA-ori with GEMMA.*** In this subsection, we compare the underlying model and test statistics of fastGWA-ori with the standard Q+K linear mixed model (LMM) implemented by GEMMA/Grid-LMM/GAPIT-MLM.

We first consider the standard Q+K model. In our benchmarking analysis, the Q part was not implemented. Thus, the model is as follows:

$$\boldsymbol{y} = \boldsymbol{1}_n\mu + \boldsymbol{m}a + \boldsymbol{g} + \boldsymbol{e}, \qquad (B1)$$

where $\boldsymbol{y}$ is the n-dimensional vector of phenotypic values (n is the number of individuals), $\boldsymbol{1}_n = (1,1,\cdots,1)'$ is the n-dimensional vector of one's, $\mu$ is the intercept term, $\boldsymbol{m}$ is the n-dimensional vector of marker profile, $a$ is the effect of the marker being tested, $\boldsymbol{g}$ is the n-dimensional vector of random polygenic effects, $\boldsymbol{e}$ is the n-dimensional vector of residuals. In the model, $\mu$ and $a$ are fixed unknown parameters, $\boldsymbol{g} \sim \boldsymbol{N}(\boldsymbol{0}, \boldsymbol{K}\sigma_g^2)$, $\boldsymbol{e} \sim \boldsymbol{N}(\boldsymbol{0}, \boldsymbol{I}\sigma_e^2)$, $\boldsymbol{K}$ is the marker-derived kinship matrix, $\boldsymbol{I}$ is the n × n identity matrix.

Let $\boldsymbol{V} = \boldsymbol{K}\sigma_g^2 + \boldsymbol{I}\sigma_e^2$. The solution of $\mu$ and $a$ for Eq. (B3) was given by the following system of linear equations:

$$\begin{pmatrix} \mathbf{1}_n{}'\boldsymbol{V}^{-1}\mathbf{1}_n & \mathbf{1}_n{}'\boldsymbol{V}^{-1}\boldsymbol{m} \\ \boldsymbol{m}'\boldsymbol{V}^{-1}\mathbf{1}_n & \boldsymbol{m}'\boldsymbol{V}^{-1}\boldsymbol{m} \end{pmatrix} \begin{pmatrix} \hat{\mu} \\ \hat{a} \end{pmatrix} = \begin{pmatrix} \mathbf{1}_n{}'\boldsymbol{V}^{-1}\boldsymbol{y} \\ \boldsymbol{m}'\boldsymbol{V}^{-1}\boldsymbol{y} \end{pmatrix}. \tag{B2}$$

And the variances of the corresponding estimates were given by:

$$\operatorname{var}\begin{pmatrix} \hat{\mu} \\ \hat{a} \end{pmatrix} = \begin{pmatrix} \mathbf{1}_n{}'\boldsymbol{V}^{-1}\mathbf{1}_n & \mathbf{1}_n{}'\boldsymbol{V}^{-1}\boldsymbol{m} \\ \boldsymbol{m}'\boldsymbol{V}^{-1}\mathbf{1}_n & \boldsymbol{m}'\boldsymbol{V}^{-1}\boldsymbol{m} \end{pmatrix}^{-1}. \tag{B3}$$

From Eq. (B2) and (B3), we can deduce $\hat{a}$ and $\operatorname{var}(\hat{a})$:

$$\hat{a} = \frac{(\mathbf{1}_n'\boldsymbol{V}^{-1}\mathbf{1}_n)(\boldsymbol{m}'\boldsymbol{V}^{-1}\boldsymbol{y}) - (\boldsymbol{m}'\boldsymbol{V}^{-1}\mathbf{1}_n)(\mathbf{1}_n'\boldsymbol{V}^{-1}\boldsymbol{y})}{(\mathbf{1}_n'\boldsymbol{V}^{-1}\mathbf{1}_n)(\boldsymbol{m}'\boldsymbol{V}^{-1}\boldsymbol{m}) - (\boldsymbol{m}'\boldsymbol{V}^{-1}\mathbf{1}_n)^2}, \tag{B4}$$

$$\operatorname{var}(\hat{a}) = \frac{\mathbf{1}_n'\boldsymbol{V}^{-1}\mathbf{1}_n}{(\mathbf{1}_n'\boldsymbol{V}^{-1}\mathbf{1}_n)(\boldsymbol{m}'\boldsymbol{V}^{-1}\boldsymbol{m}) - (\boldsymbol{m}'\boldsymbol{V}^{-1}\mathbf{1}_n)^2}. \tag{B5}$$

Thus, the Wald-test statistic resulting from the model (B1) was the following:

$$W_{\text{GEMMA}} = \frac{\hat{a}^2}{\operatorname{var}(\hat{a})} = \frac{\left[(\mathbf{1}_n'\boldsymbol{V}^{-1}\mathbf{1}_n)(\boldsymbol{m}'\boldsymbol{V}^{-1}\boldsymbol{y}) - (\boldsymbol{m}'\boldsymbol{V}^{-1}\mathbf{1}_n)(\mathbf{1}_n'\boldsymbol{V}^{-1}\boldsymbol{y})\right]^2}{(\mathbf{1}_n'\boldsymbol{V}^{-1}\mathbf{1}_n)\left[(\mathbf{1}_n'\boldsymbol{V}^{-1}\mathbf{1}_n)(\boldsymbol{m}'\boldsymbol{V}^{-1}\boldsymbol{m}) - (\boldsymbol{m}'\boldsymbol{V}^{-1}\mathbf{1}_n)^2\right]}. \tag{B6}$$

The algorithm fastGWA-ori was based on a slightly different model:

$$\tilde{\boldsymbol{y}} = \tilde{\boldsymbol{m}}a + \boldsymbol{g} + \boldsymbol{e}, \tag{B7}$$

where $\tilde{\boldsymbol{y}}$ is the centered vector of phenotypic values, $\tilde{\boldsymbol{m}}$ is the centered vector of marker profiles, the other notations and assumptions are the same as in Eq. (B1). With this model, we have

$$\hat{a} = \tilde{\boldsymbol{m}}'\boldsymbol{V}^{-1}\tilde{\boldsymbol{y}}, \qquad \operatorname{var}(\hat{a}) = \frac{1}{\tilde{\boldsymbol{m}}'\boldsymbol{V}^{-1}\tilde{\boldsymbol{m}}}. \tag{B8}$$

Hence, the Wald-test statistic is given by the following:

$$W_{\text{fastGWA-ori}} = \frac{\hat{a}^2}{\operatorname{var}(\hat{a})} = \frac{(\tilde{\boldsymbol{m}}'\boldsymbol{V}^{-1}\tilde{\boldsymbol{y}})^2}{\tilde{\boldsymbol{m}}'\boldsymbol{V}^{-1}\tilde{\boldsymbol{m}}}. \tag{B9}$$

Note that in Eq. (B6) and (B9), there are two unknown parameters $\sigma_g^2$ and $\sigma_e^2$ (hidden in the matrix $\boldsymbol{V}$). To calculate the test statistics, they are replaced by the corresponding maximum likelihood estimates $\hat{\sigma}_g^2$ and $\hat{\sigma}_e^2$. In general, the likelihood functions of model (B1) and (B7) are different. Thus, the two models would provide different estimates for $\hat{\sigma}_g^2$ and $\hat{\sigma}_e^2$. However, we have the following interesting result:

**Proposition B1.** Suppose that the estimated $\hat{\sigma}_g^2$ and $\hat{\sigma}_e^2$ are the same from the two models (B1) and (B7), then the Wald-test statistics calculated by the two models (Eq. B6 and B9) are also the same, provided that the marker-derived kinship matrix has the form $\boldsymbol{K} = \boldsymbol{M}\boldsymbol{M}'/d$, where $\boldsymbol{M}$ is a matrix of centered marker profiles (i.e. the sum of entries in each column of $\boldsymbol{M}$ is zero) and $d$ is a constant.

A strict mathematical proof is provided at the end of this note. As we can see in the next subsection, the additional assumption on the kinship matrix is fulfilled in all algorithms considered in this note. Proposition B1 indicates that if we run fastGWA-ori and GEMMA with the same data set and based on the same kinship matrix of form $\boldsymbol{M}\boldsymbol{M}'/d$, their difference in the test statistics is solely determined by the estimated variance components $\hat{\sigma}_g^2$ and $\hat{\sigma}_e^2$.

***Comparing the default kinship matrix of fastGWA-ori and GEMMA.*** To make a meaningful comparison between the performance of fastGWA-ori and GEMMA based on simulated data sets, we need to make sure that they are based on the same kinship matrices. In fact, the default methods of calculating the kinship matrix in these two algorithms are different.

Let $\boldsymbol{M} = (m_{ij})_{n\times p}$ be the matrix of marker profiles coded as 0, 1, 2 (p is the number of all markers). With default setting of fastGWA, the kinship matrix was calculated based on the standardized marker vectors, namely as following:

$$\boldsymbol{K}_s = \frac{1}{p}\boldsymbol{N}\boldsymbol{N}', \qquad \boldsymbol{N} = (n_{ij})_{n\times p}, \qquad n_{ij} = \frac{m_{ij} - 2p_j}{\sqrt{2p_j(1-p_j)}}\,, \qquad (B10)$$

where $p_j$ is the frequency of the reference allele for the $j$-th marker.

In GEMMA/Grid-LMM/GAPIT-MLM, the default method for calculating the kinship matrix is based on centered marker vectors without being divided by the expected standard deviation, namely the following:

$$\boldsymbol{K}_c = \frac{1}{c}\boldsymbol{Z}\boldsymbol{Z}', \qquad \boldsymbol{Z} = (z_{ij})_{n\times p}, \qquad z_{ij} = m_{ij} - 2p_j. \qquad (B11)$$

where $c = p$ in GEMMA/Grid-LMM, and $c = \sum_{s=1}^{p} 2p_s(1-p_s)$ in GAPIT-MLM. Since multiplying the kinship matrix by a constant does not affect the resulting test statistics, it is safe to treat the default kinship matrix used by GEMMA, Grid-LMM and GAPIT-MLM as the same.

We noticed that fastGWA provides the option of using $\boldsymbol{K}_c$ (with $c = \sum_{s=1}^{p} 2p_s(1-p_s)$) as the kinship matrix, and GEMMA has the option to change the kinship matrix to $\boldsymbol{K}_s$. Thus, comparing the performance of fastGWA-ori and GEMMA based on the same kinship matrix is possible.

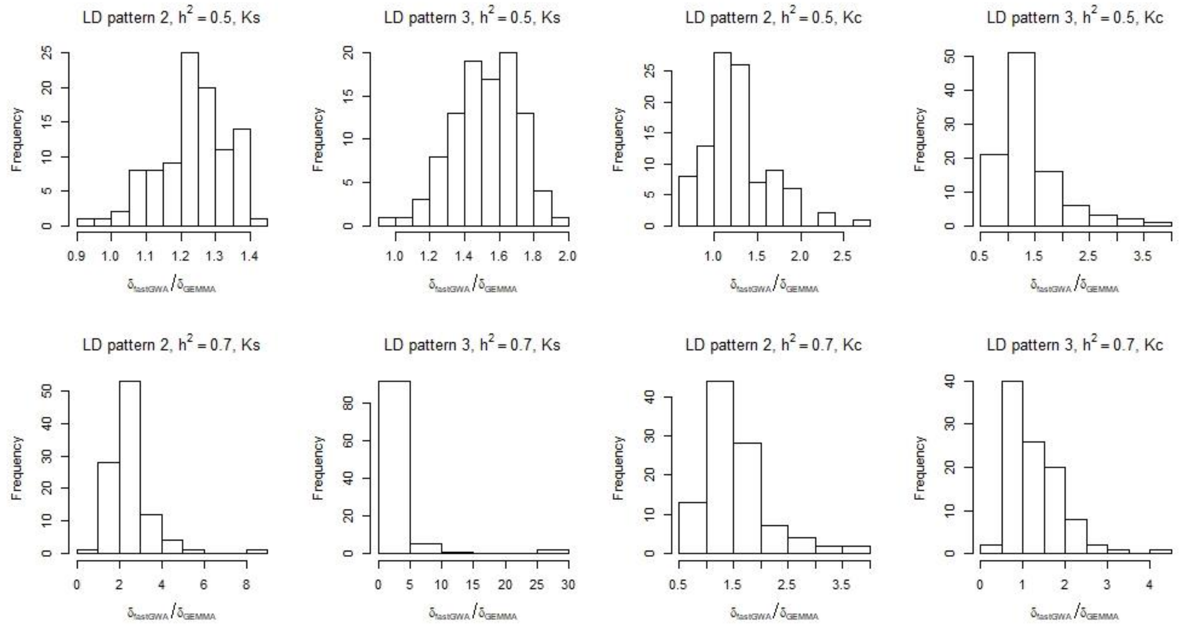

**Figure B1.** The distribution of $\hat{\delta}_{\text{fastGWA}}/\hat{\delta}_{\text{GEMMA}}$ value in each of the 4 scenarios (LD pattern 2 and 3, trait heritability 0.5 and 0.7) based on two different kinship matrices ($\boldsymbol{K}_s$ and $\boldsymbol{K}_c$).

***Comparing the performance of fastGWA-ori and GEMMA***. In this subsection, we compared fastGWA-ori with GEMMA on simulated data sets. To reduce the computational time, we chose 4 out of the 108 scenarios in the benchmarking analysis. Since each scenario consisted of 100 simulated data sets, in total there were 400 data sets. The 4 scenarios represented two different trait heritabilities (0.5 and 0.7) and two different LD patterns (LD pattern 2 and 3). The other parameters were fixed: The population size was 1,000. There were six major QTL explaining from 2% to 12 % (with a step of 2%) proportion of genetic variance (PG2), and 200 markers from each chromosome contributed as minor QTL to the genetic background effects (GB1). In LD pattern 2, there was no LD between any two major QTL, but LD exited between major and minor QTL. In LD pattern 3, there existed LD among the major QTL as well as between major and minor QTL.

We first compared the estimated ratio of variance components $\hat{\delta} = \hat{\sigma}_g^2/\hat{\sigma}_e^2$ with both kinship matrices. Note that the different scaling of the kinship matrix in GEMMA and fastGWA was accounted for in this analysis. We observed substantial difference in the estimated $\hat{\delta}$ values between the two algorithms (Figure B1). In general, fastGWA tended to produce higher $\hat{\delta}$ values than GEMMA, regardless which kinship matrix was used. Hence, by Proposition B1, we would not expect that fastGWA and GEMMA produce similar test statistics. This was confirmed by the observation that the power and FPR of fastGWA-ori were different from that of GEMMA in all 4 scenarios (Tables B1, B2). Compared with GEMMA, the power and FPR of fastGWA-ori was lower with $\boldsymbol{K}_s$, but higher with $\boldsymbol{K}_c$. We also observed that the difference between GEMMA based on the two distinct kinship matrices was very small. In contrast, the difference was much more pronounced for fastGWA-ori.

**Table B1.** The statistical power of three algorithms (GEMMA, fastGWA-ori and fastGWA-GG), each with two different kinship matrices, evaluated in four scenarios. Each scenario consisted of 100 simulated phenotypes.

| Algorithm | Kinship matrix | Heritability 0.5 | | Heritability 0.7 | |
|---|---|---|---|---|---|
| | | LD pattern 2 | LD pattern 3 | LD pattern 2 | LD pattern 3 |
| GEMMA | $K_c$ | 0.493 | 0.628 | 0.588 | 0.682 |
| GEMMA | $K_s$ | 0.500 | 0.623 | 0.592 | 0.690 |
| fastGWA-ori | $K_s$ | 0.365 | 0.455 | 0.445 | 0.442 |
| fastGWA-ori | $K_c$ | 0.597 | 0.717 | 0.718 | 0.798 |
| fastGWA-GG | $K_s$ | 0.395 | 0.465 | 0.385 | 0.320 |
| fastGWA-GG | $K_c$ | 0.582 | 0.657 | 0.628 | 0.667 |

**Table B2.** The false-positive rate (FPR) of three algorithms (GEMMA, fastGWA-ori and fastGWA-GG), each with two different kinship matrices, evaluated in four scenarios. Each scenario consisted of 100 simulated phenotypes.

| Algorithm | Kinship matrix | Heritability 0.5 | | Heritability 0.7 | |
|---|---|---|---|---|---|
| | | LD pattern 2 | LD pattern 3 | LD pattern 2 | LD pattern 3 |
| GEMMA | $K_c$ | $5.73 \times 10^{-5}$ | $1.18 \times 10^{-4}$ | $3.74 \times 10^{-4}$ | $6.29 \times 10^{-4}$ |
| GEMMA | $K_s$ | $5.77 \times 10^{-5}$ | $9.92 \times 10^{-5}$ | $3.58 \times 10^{-4}$ | $5.24 \times 10^{-4}$ |
| fastGWA-ori | $K_s$ | $2.90 \times 10^{-5}$ | $1.87 \times 10^{-5}$ | $1.16 \times 10^{-4}$ | $9.40 \times 10^{-5}$ |
| fastGWA-ori | $K_c$ | $1.97 \times 10^{-4}$ | $3.39 \times 10^{-4}$ | $1.22 \times 10^{-3}$ | $1.84 \times 10^{-3}$ |
| fastGWA-GG | $K_s$ | $2.09 \times 10^{-5}$ | $1.58 \times 10^{-5}$ | $7.60 \times 10^{-5}$ | $3.90 \times 10^{-5}$ |
| fastGWA-GG | $K_c$ | $1.84 \times 10^{-4}$ | $2.84 \times 10^{-4}$ | $1.04 \times 10^{-3}$ | $1.56 \times 10^{-3}$ |

***Comparing fastGWA-ori and fastGWA-GG.*** Now, we can discuss the performance of fastGWA-GG. Note that the underlying model and the procedure of solving the LMM in fastGWA-GG was the same as in fastGWA-ori, hence they produced the same estimates for $\hat{\sigma}_g^2$ and $\hat{\sigma}_e^2$. From the results of previous subsection, we know that we cannot draw any conclusion on the influence of the GRAMMAR-Gamma approximation by simply comparing fastGWA-GG with GEMMA because there is an additional difference between the two algorithms, namely their underlying model (which lead to different values for $\hat{\sigma}_g^2$ and $\hat{\sigma}_e^2$).

Therefore, it is necessary to use fastGWA-ori as a bridge. On one hand, the influence of the GRAMMAR-Gamma approximation can be inferred by comparing fastGWA-GG and fastGWA-ori. On the other hand, the influence of the slight difference in the underlying model (or equivalently, the different $\hat{\sigma}_g^2$ and $\hat{\sigma}_e^2$ values, by Proposition B1) can be investigated by comparing fastGWA-ori with GEMMA. In this way, we could find out which of the two differences played a more important role for the low power of fastGWA-GG.

With the default kinship matrix $\boldsymbol{K}_s$, fastGWA-ori already produced much lower power than GEMMA. Compared with fastGWA-ori, the power of fastGWA-GG with $\boldsymbol{K}_s$ was slightly higher in the two scenarios with heritability 0.5, but lower in scenarios with heritability 0.7 (Table B1). With the alternative kinship matrix $\boldsymbol{K}_c$, the power of fastGWA-GG was consistently lower than fastGWA-ori, both were higher than GEMMA. However, their FPR were also inflated, especially in the case of heritability 0.7, which exceeded 0.1% (Table B2). In our benchmarking analysis, fastGWA-GG was evaluated with the default kinship matrix $\boldsymbol{K}_s$. Thus, we may conclude that the observed low power for fastGWA-GG was likely due to the low power of fastGWA-ori, i.e. the slightly different underlying model (Eq. B7) instead of the GRAMMAR-Gamma approximation. Of course, since we only performed a small-scale simulation study to compare fastGWA-ori and GEMMA, further analysis is needed to verify this conclusion.

##### B.2 On the performance of fastGWA-sp

In the main text, we observed that fastGWA-sp had small FPR in some scenarios and high FPR in other scenarios. Here, we took one scenario in which the FPR was very high and did further analyses to find out the reason. For the 100 simulated data sets in this scenario, we extracted the FPR for each data set and found that the FPR of fastGWA-sp was higher than 0.001 in 27 data sets (Table B3). In contrast, the FPR of GEMMA/Grid-LMM/GAPIT-MLM was less than 0.001 in all 100 data sets. Surprisingly, there were even 5 data sets in which the FPR of fastGWA-sp was 100%.

**Table B3.** The number of simulated data sets in six different intervals of false positive rate for four algorithms fastGWA-sp, GEMMA, Grid-LMM and GAPIT-MLM. The total number of data sets is 100 and they are from a single simulation scenario. In this scenario, there existed LD among the major QTL as well as between major and minor QTL (LD pattern 3). The simulated trait heritability was 0.7 with a population size of 3,000. There were six major QTL each explaining 2% of the genetic variance (PG1), and 200 markers from each chromosome contributed as genetic background (GB2).

| FPR interval | GEMMA | Grid-LMM | GAPIT-MLM | fastGWA-sp |
|---|---|---|---|---|
| FPR < 0.0001 | 61 | 61 | 64 | 23 |
| 0.0001 ≤ FPR < 0.001 | 39 | 39 | 36 | 50 |
| 0.001 ≤ FPR < 0.01 | 0 | 0 | 0 | 16 |
| 0.01 ≤ FPR < 0.1 | 0 | 0 | 0 | 1 |
| 0.1 ≤ FPR < 1 | 0 | 0 | 0 | 5 |
| FPR=1 | 0 | 0 | 0 | 5 |

Then, we chose three data sets for a detailed case study: Rep_5 in which FPR = 1, Rep_78 in which FPR = 0.24 and Rep_50 in which FPR = 0.00026. We found that the output *p*-values of fastGWA-sp for Rep_5 were zero for all markers, which is clearly erroneous. To investigate whether the error was caused by the sparse kinship matrix, we extracted the sparse kinship matrix generated by fastGWA-sp and plugged it in GEMMA to re-calculate the test statistics. This was done for all three data sets. For convenience, this approach was denoted by GEMMA-sp.

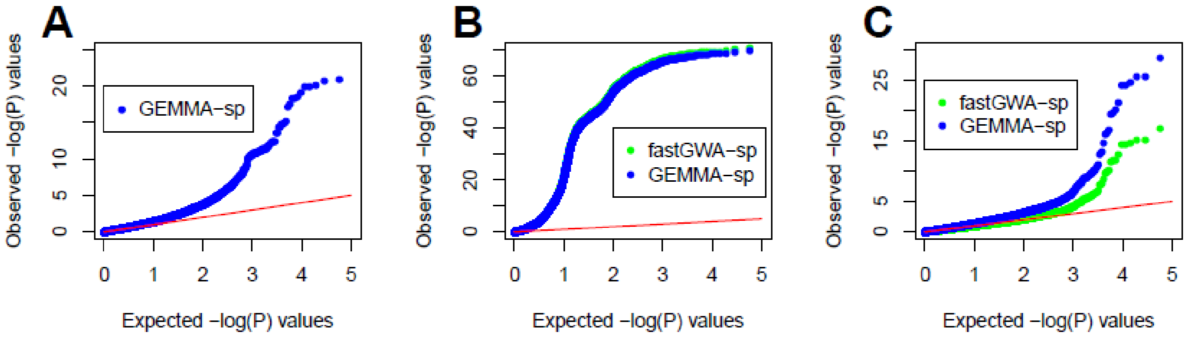

**Figure B2.** Quantile-quantile plots of -log$_{10}$(*p*) values produced by fastGWA-sp and GEMMA-sp in three data sets in a scenario with LD pattern 3, trait heritability 0.7, population size 3,000, GB1 and PG1. **A)** data set Rep_5. Only the -log$_{10}$(*p*) values of GEMMA-sp were plotted, because the *p*-values of fastGWA-sp were all zero. **B)** data set Rep_78. **C)** data set Rep_50.

We observed that GEMMA-sp produced normal test statistics in Rep_5 (Figure B2, Panel A), indicating that the sparse kinship matrix was not responsible for the 100% FPR produced by fastGWA. We further checked that the $\hat{\delta} = \hat{\sigma}_g^2/\hat{\sigma}_e^2$ values estimated by fastGWA-sp and GEMMA-sp were also similar

($\hat{\delta}_{\text{fastGWA}-\text{sp}}$ = 0.33, $\hat{\delta}_{\text{GEMMA}-\text{sp}}$ = 0.38). Note that although the original kinship matrix $\boldsymbol{K}_S$ satisfies the assumptions in Proposition B1, the sparse kinship matrix (obtained from $\boldsymbol{K}_S$ by setting entries below 0.05 to 0) may not. Thus, we cannot apply Proposition B1 to conclude that the test statistics provided by fastGWA-sp and GEMMA-sp are also similar. Nevertheless, the estimated $\hat{\delta}$ value looks quite normal and is not likely the source of any error. We anticipated that the erroneous *p*-values might be a result of unexpected numerical errors in the computation of test statistics. Since GEMMA-sp seemed not affected by the issue, we may guess that the sparse kinship matrix is ill-conditioned and the underlying model of GEMMA (Eq. B1) is numerically more stable than that of fastGWA (Eq. B7).

Then, we looked at the results of data set Rep_78 in which the FPR of fastGWA-sp was clearly inflated but not erroneous as in Rep_5. In this case, the *p*-values provided by fastGWA-sp and GEMMA-sp were almost the same (Figure B2, Panel B). Hence, unlike Rep_5, in this case the sparse kinship matrix was indeed responsible for the inflated FPR. Since the sparse kinship matrix was obtained from the original kinship matrix by setting entries below 0.05 to 0, it simplified the structure of the matrix and this might lead to an insufficient control of the population stratification and genetic relatedness. In the data set Rep_50 in which the FPR of fastGWA-sp was well-controlled, GEMMA-sp seemed to have higher power than fastGWA-sp (Figure B2, Panel C). This result was consistent to what we observed for fastGWA-ori and GEMMA with the original kinship matrix $\boldsymbol{K}_S$ (Table B1).

In view of the above results, our conclusions are: 1) The 100% FPR observed in a small proportion of data sets might be due to numerical errors in the computation of test statistics of fastGWA-sp. 2) The sparse kinship technique introduced by fastGWA-sp may lead to insufficient control of the population structure, which in turn results in inflated FPR, as observed in a certain proportion of data sets. Further analyses are out of the scope of this study, but are needed to confirm these conclusions.

**B.3 The proof of Proposition B1.**

We shall use the key assumption in the proposition, i.e., the kinship matrix is of the form $\boldsymbol{K} = \boldsymbol{M}\boldsymbol{M}'/d$, and $\boldsymbol{M}$ is column-centered.

First, when $\boldsymbol{K} = \boldsymbol{M}\boldsymbol{M}'/d$, we can use results from basic linear algebra to calculate the inverse of $\boldsymbol{V} = \boldsymbol{K}\sigma_g^2 + \boldsymbol{I}\sigma_e^2$, namely we have ($\delta = \sigma_g^2/\sigma_e^2$):

$$\boldsymbol{V}^{-1} = [\sigma_e^2(\boldsymbol{I} + \boldsymbol{K}\delta)]^{-1} = \sigma_e^{-2}(\boldsymbol{I} + \boldsymbol{M}\boldsymbol{M}'\delta/d)^{-1}$$

$$= \sigma_e^{-2}\left[\boldsymbol{I} - \boldsymbol{M}\left(\frac{\delta}{d}\boldsymbol{I} + \boldsymbol{M}'\boldsymbol{M}\right)^{-1}\boldsymbol{M}'\right]. \qquad (B12)$$

Then, because $\boldsymbol{M}$ is column-centered, we know that $\boldsymbol{1}_n'\boldsymbol{M} = \boldsymbol{0}_{1\times n}$, $\boldsymbol{M}'\boldsymbol{1}_n = \boldsymbol{0}_{n\times 1}$. Thus, we have

$$\boldsymbol{1}_n'\boldsymbol{V}^{-1}\boldsymbol{1}_n = \sigma_e^{-2}n, \qquad \boldsymbol{m}'\boldsymbol{V}^{-1}\boldsymbol{1}_n = \sigma_e^{-2}n\bar{m}, \qquad \boldsymbol{1}_n'\boldsymbol{V}^{-1}\boldsymbol{y} = \sigma_e^{-2}n\bar{y}, \qquad (B13)$$

Using Eq. (B13), we can rewrite Eq. (B6) as following:

$$W_{\text{GEMMA}} = \frac{\left[(\boldsymbol{1}_n'\boldsymbol{V}^{-1}\boldsymbol{1}_n)(\boldsymbol{m}'\boldsymbol{V}^{-1}\boldsymbol{y}) - (\boldsymbol{m}'\boldsymbol{V}^{-1}\boldsymbol{1}_n)(\boldsymbol{1}_n'\boldsymbol{V}^{-1}\boldsymbol{y})\right]^2}{(\boldsymbol{1}_n'\boldsymbol{V}^{-1}\boldsymbol{1}_n)[(\boldsymbol{1}_n'\boldsymbol{V}^{-1}\boldsymbol{1}_n)(\boldsymbol{m}'\boldsymbol{V}^{-1}\boldsymbol{m}) - (\boldsymbol{m}'\boldsymbol{V}^{-1}\boldsymbol{1}_n)^2]}$$

$$= \frac{\left[\sigma_e^{-2}n(\boldsymbol{m}'\boldsymbol{V}^{-1}\boldsymbol{y}) - \sigma_e^{-4}n^2\bar{m}\bar{y}\right]^2}{\sigma_e^{-2}n[\sigma_e^{-2}n(\boldsymbol{m}'\boldsymbol{V}^{-1}\boldsymbol{m}) - \sigma_e^{-4}n^2\bar{m}^2]} = \frac{\boldsymbol{m}'\boldsymbol{V}^{-1}\boldsymbol{y} - \sigma_e^{-2}n\bar{m}\bar{y}}{\boldsymbol{m}'\boldsymbol{V}^{-1}\boldsymbol{m} - \sigma_e^{-2}n\bar{m}^2}. \qquad (B14)$$

Next, using $\tilde{\boldsymbol{y}} = \boldsymbol{y} - \boldsymbol{1}_n\bar{y}$, $\tilde{\boldsymbol{m}} = \boldsymbol{m} - \boldsymbol{1}_n\bar{m}$, we can rewrite Eq. (B9) as following:

$$W_{\text{fastGWA-ori}} = \frac{(\tilde{\boldsymbol{m}}'\boldsymbol{V}^{-1}\tilde{\boldsymbol{y}})^2}{\tilde{\boldsymbol{m}}'\boldsymbol{V}^{-1}\tilde{\boldsymbol{m}}} = \frac{\left[(\boldsymbol{m}' - \boldsymbol{1}_n'\bar{m})\boldsymbol{V}^{-1}(\boldsymbol{y} - \boldsymbol{1}_n\bar{y})\right]^2}{(\boldsymbol{m}' - \boldsymbol{1}_n'\bar{m})\boldsymbol{V}^{-1}(\boldsymbol{m} - \boldsymbol{1}_n\bar{m})}$$

$$= \frac{\boldsymbol{m}'\boldsymbol{V}^{-1}\boldsymbol{y} - \bar{m}\boldsymbol{1}_n'\boldsymbol{V}^{-1}\boldsymbol{y} - \bar{y}\boldsymbol{m}'\boldsymbol{V}^{-1}\boldsymbol{1}_n + \bar{m}\bar{y}\boldsymbol{1}_n'\boldsymbol{V}^{-1}\boldsymbol{1}_n}{\boldsymbol{m}'\boldsymbol{V}^{-1}\boldsymbol{m} - 2\bar{m}\boldsymbol{m}'\boldsymbol{V}^{-1}\boldsymbol{1}_n + \bar{m}^2\boldsymbol{1}_n'\boldsymbol{V}^{-1}\boldsymbol{1}_n}$$

$$= \frac{\boldsymbol{m}'\boldsymbol{V}^{-1}\boldsymbol{y} - \bar{m}\cdot\sigma_e^{-2}n\bar{y} - \bar{y}\cdot\sigma_e^{-2}n\bar{m} + \bar{m}\bar{y}\cdot\sigma_e^{-2}n}{\boldsymbol{m}'\boldsymbol{V}^{-1}\boldsymbol{m} - 2\bar{m}\cdot\sigma_e^{-2}n\bar{m} + \bar{m}^2\cdot\sigma_e^{-2}n} = \frac{\boldsymbol{m}'\boldsymbol{V}^{-1}\boldsymbol{y} - \sigma_e^{-2}n\bar{m}\bar{y}}{\boldsymbol{m}'\boldsymbol{V}^{-1}\boldsymbol{m} - \sigma_e^{-2}n\bar{m}^2}. \qquad (B15)$$

Now, it is clear that Eq. (B14) and (B15) are the same, which means that Eq. (B6) and (B9) are equal. Hence, the proof is completed.

#### Supplementary Note C

In this supplementary note, we briefly discussed the performance of two recently published algorithms MM4LMM and REGENIE in 400 simulated data sets. The two algorithms were published after we had started the benchmarking analysis. Since they implemented different techniques from the 12 selected algorithms, we decided to evaluate them with a sub-collection of the 10,800 simulated data sets used in our benchmarking analysis.

More precisely, we chose 4 scenarios each consisting of 100 simulated data sets, resulting in 400 data sets. The 4 scenarios represented two different population size (PS300 and PS1000) and two different LD patterns (LD pattern 2 and 3). The other parameters were fixed: The trait heritability was 0.7. There were six major QTL explaining from 2% to 12 % (with a step of 2%) proportion of genetic variance (PG2), and 1,200 markers contributed as minor QTL to the genetic background effects (GB1). Note that in LD pattern 2, there was no LD between any two major QTL, but LD exited between major and minor QTL. In LD pattern 3, there existed LD among the major QTL as well as between major and minor QTL.

The statistical power and false positive rate (FPR) of the two algorithms were shown in Table C1 and C2, respectively. Since MM4LMM is an exact algorithm, the results of GEMMA were included for comparison. Similarly, the results of BOLT-LMM-inf was included to be compared with REGENIE, as both implemented the leave-one-chromosome-out (LOCO) technique.

**Table C1.** The statistical power of four algorithms (GEMMA, BOLT-LMM-inf, REGENIE, MM4LMM) evaluated in four scenarios, each consisting of 100 simulated phenotypes

| Algorithm | PS300 | | PS1000 | |
|---|---|---|---|---|
| | LD pattern 2 | LD pattern 3 | LD pattern 2 | LD pattern 3 |
| GEMMA | 0.172 | 0.318 | 0.628 | 0.681 |
| BOLT-LMM-inf | 0.260 | 0.555 | 0.752 | 0.867 |
| REGENIE | 0.123 | 0.630 | 0.732 | 0.927 |
| MM4LMM | 0.200 | 0.333 | 0.628 | 0.688 |

**Table C2.** The false positive rate (FPR) of four algorithms (GEMMA, BOLT-LMM-inf, REGENIE, MM4LMM) evaluated in four scenarios, each consisting of 100 simulated phenotypes.

| Algorithm | PS300 | | PS1000 | |
|---|---|---|---|---|
| | LD pattern 2 | LD pattern 3 | LD pattern 2 | LD pattern 3 |
| GEMMA | $2.06 \times 10^{-5}$ | $2.05 \times 10^{-4}$ | $1.18 \times 10^{-4}$ | $6.29 \times 10^{-4}$ |
| BOLT-LMM-inf | $1.63 \times 10^{-3}$ | $1.86 \times 10^{-2}$ | $1.17 \times 10^{-2}$ | $3.65 \times 10^{-2}$ |
| REGENIE | $5.41 \times 10^{-3}$ | $4.37 \times 10^{-2}$ | $3.56 \times 10^{-2}$ | $7.66 \times 10^{-2}$ |
| MM4LMM | $3.84 \times 10^{-5}$ | $2.26 \times 10^{-4}$ | $1.22 \times 10^{-4}$ | $6.42 \times 10^{-4}$ |

We observed that the power and FPR of MM4LMM were comparable to that of GEMMA, which was expected as both are exact algorithms. The performance of the two algorithms was quite similar in the data sets with 1,000 individuals, while MM4LMM produced slightly higher power than GEMMA when the population size was 300. Further analyses are needed to verify whether the higher power is a consistent phenomenon.

The power of REGENIE was similar to or higher than BOLT-LMM-inf in three scenarios, and was lower than BOLT-LMM-inf in LD pattern 2 when the population size was 300. The FPR of REGENIE was comparable to that of BOLT-LMM-inf, which was substantially higher than that of GEMMA and MM4LMM. This was likely due to the implementation of LOCO, as we observed inflated FPR for all algorithms implementing LOCO (BOLT-LMM-inf, BOLT-LMM-mix, MLMA-LOCO) in the benchmarking analysis (see the main text).

scenarios, each consisting of 100 simulated phenotypes. In all scenarios, there were six major QTL explaining from 2% to 12% (with a step of 2%) proportion of genetic variance (PG2), and 1,200 markers contributed as minor QTL to the genetic background effects (GB1). The trait heritability was 0.7. The population size was 300 for two scenarios (PS300), and 1000 for the other two (PS1000). In LD pattern 2, there is no LD between any two major QTL, but LD exits between major and minor QTL. In LD pattern 3, there exists LD among the major QTL as well as between major and minor QTL. BOLT-LMM-inf-GLOCO and BOLT-LMM-mix-GLOCO are variants of BOLT-LMM-inf and BOLT-LMM-mix, where a genuine leave-one-chromosome-out (LOCO) procedure was forced.

| Algorithm | PS300 | | PS1000 | |
|---|---|---|---|---|
| | LD pattern 2 | LD pattern 3 | LD pattern 2 | LD pattern 3 |
| GEMMA | $2.06 \times 10^{-5}$ | $2.05 \times 10^{-4}$ | $1.18 \times 10^{-4}$ | $6.29 \times 10^{-4}$ |
| GAPIT-MLM | $4.19 \times 10^{-6}$ | $1.12 \times 10^{-4}$ | $9.00 \times 10^{-5}$ | $5.32 \times 10^{-4}$ |
| BOLT-LMM-inf | $1.63 \times 10^{-3}$ | $1.86 \times 10^{-2}$ | $1.17 \times 10^{-2}$ | $3.65 \times 10^{-2}$ |
| BOLT-LMM-mix | $2.14 \times 10^{-3}$ | $2.03 \times 10^{-2}$ | $1.74 \times 10^{-2}$ | $3.80 \times 10^{-2}$ |
| BOLT-LMM-inf-GLOCO | $1.64 \times 10^{-3}$ | $1.85 \times 10^{-2}$ | $1.15 \times 10^{-2}$ | $3.59 \times 10^{-2}$ |
| BOLT-LMM-mix-GLOCO | $1.91 \times 10^{-3}$ | $2.05 \times 10^{-2}$ | $1.60 \times 10^{-2}$ | $1.48 \times 10^{-2}$ |

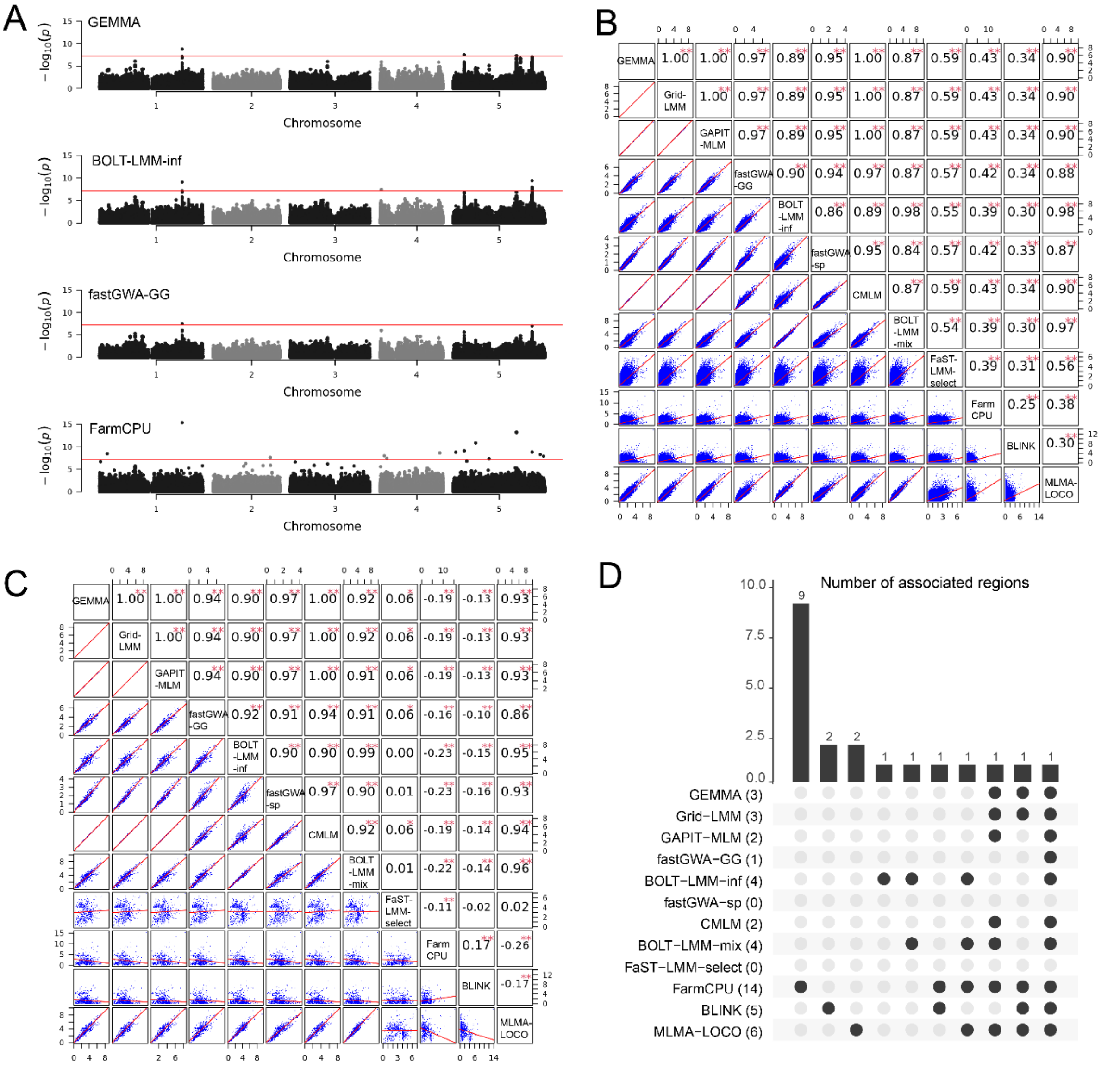

**Supplementary Figure 1.** A comparison of the results of 12 GWAS algorithms for flowering time in an Arabidopsis data set consisting of 1,003 individuals and 749,722 markers. **A.** The Manhattan plots of 4 selected algorithms. The threshold was $p < 0.05$ after Bonferroni correction. **B.** Correlations between the -log$_{10}$($p$) values of all markers obtained by each of the indicated pairs of algorithms. The names of the algorithms were indicated in the diagonal blocks. **C.** Pairwise correlations between the -log$_{10}$($p$) values of markers which were significant under a liberal threshold (-log$_{10}$($p$) > 4) in at least one algorithm. **D.** A comparison of significant regions identified by the 12 algorithms. The bar plot showed the number of regions commonly identified by the algorithms indicated by the black dots. The number of regions identified by each algorithm was presented in the parentheses next to the names of the algorithms.

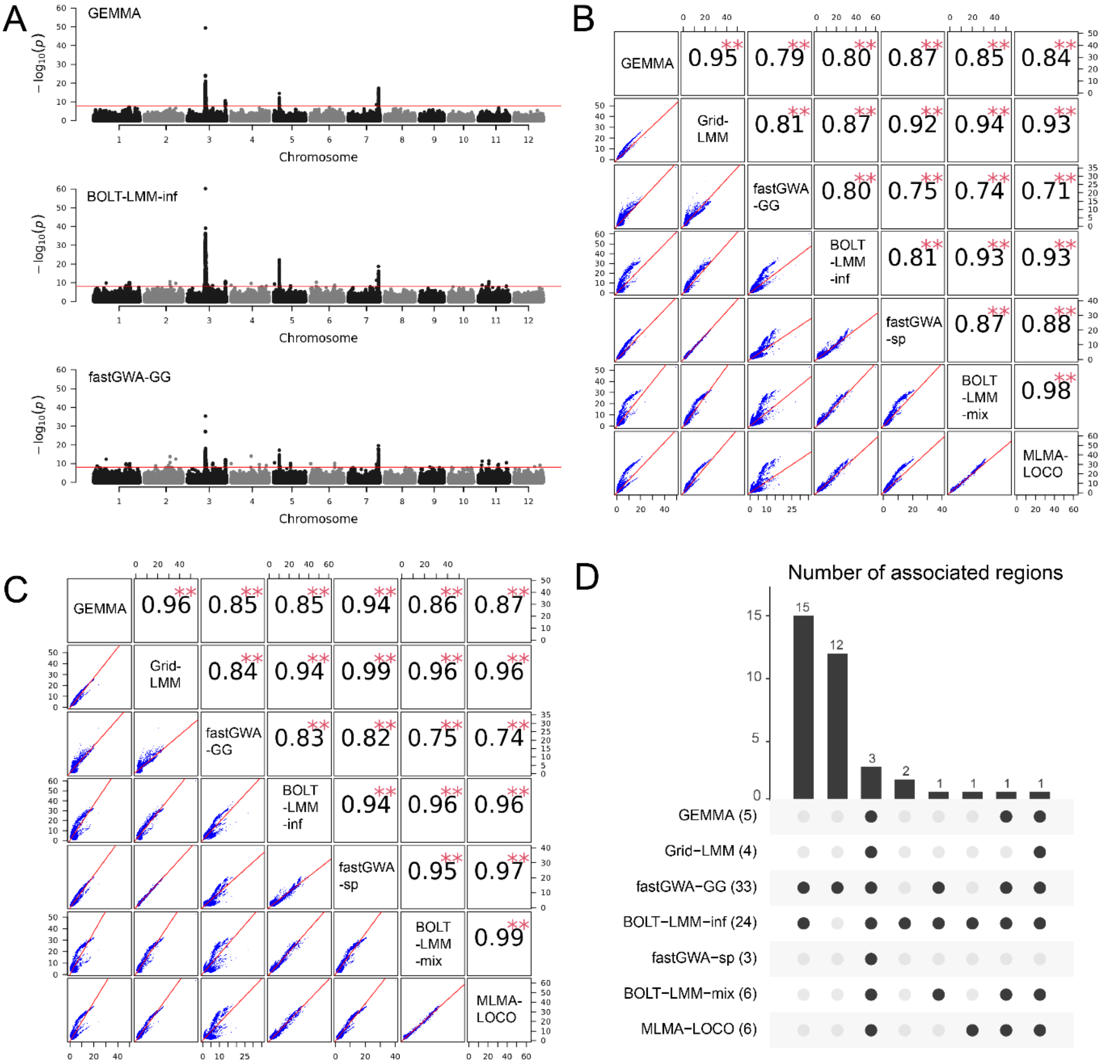

**Supplementary Figure 2.** A comparison of the results of 7 GWAS algorithms for grain length in a rice data set consisting of 3,024 individuals and 4,817,964 markers. **A.** The Manhattan plots of 4 selected algorithms. The threshold was $p < 0.05$ after Bonferroni correction. **B.** Correlations between the $-\log_{10}(p)$ values of all markers obtained by each of the indicated pairs of algorithms. The names of the algorithms were indicated in the diagonal blocks. **C.** Pairwise correlations between the $-\log_{10}(p)$ values of markers which were significant under a liberal threshold ($-\log_{10}(p) > 4$) in at least one algorithm. **D.** A comparison of significant regions identified by the 7 algorithms. The bar plot showed the number of regions commonly identified by the algorithms indicated by the black dots. The number of regions identified by each algorithm was presented in the parentheses next to the names of the algorithms. The algorithms GAPIT-MLM, CMLM, FarmCPU, BLINK and FaST-LMM-select were not evaluated due to the computational load.

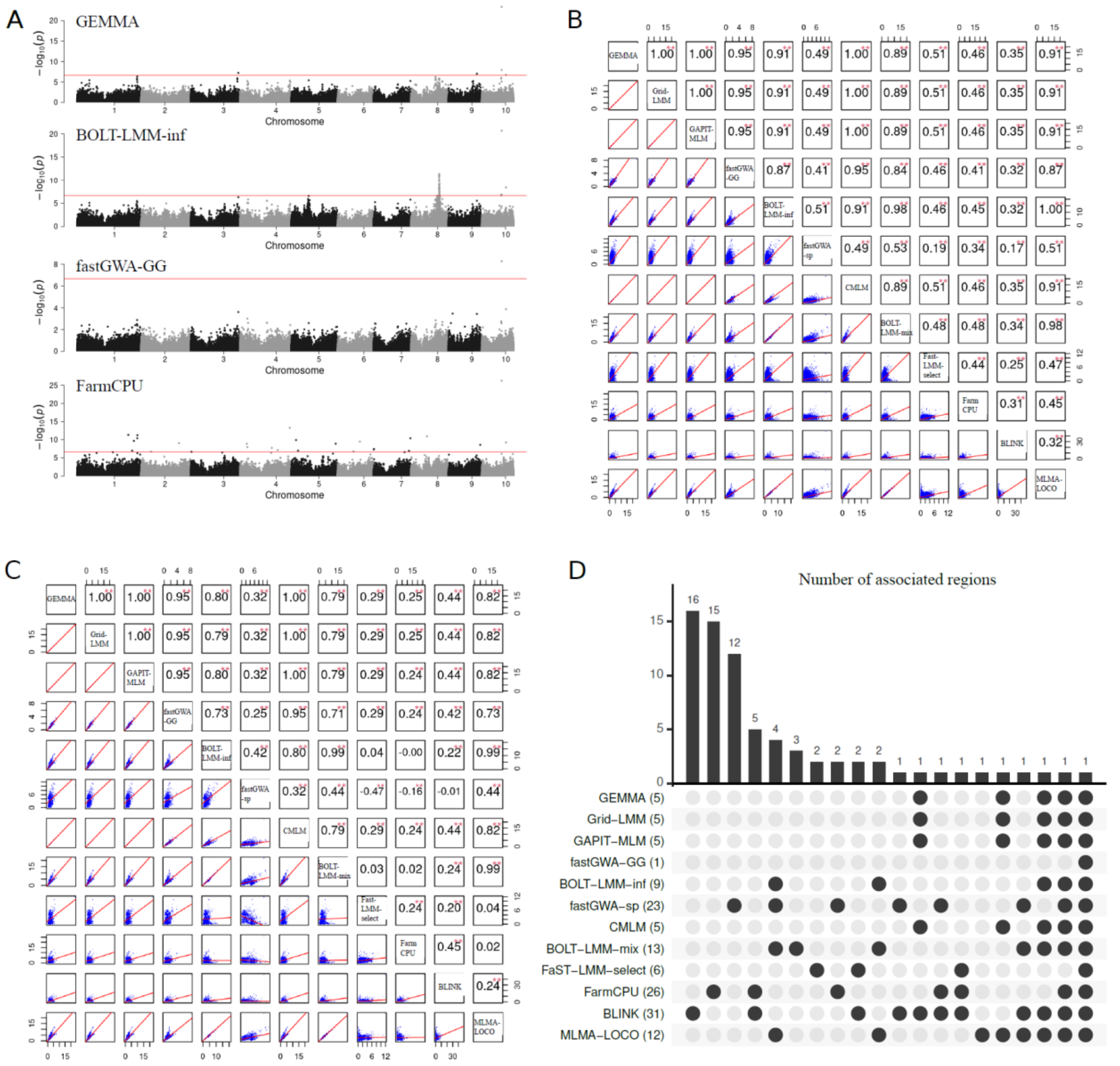

**Supplementary Figure 3.** A comparison of the results of 12 GWAS algorithms for days to silking in a maize data set consisting of 2,279 individuals and 225,563 markers. **A.** The Manhattan plots of 4 selected algorithms. The threshold was $p < 0.05$ after Bonferroni correction. **B.** Correlations between the $-\log_{10}(p)$ values of all markers obtained by each of the indicated pairs of algorithms. The names of the algorithms were indicated in the diagonal blocks. **C.** Pairwise correlations between the $-\log_{10}(p)$ values of markers which were significant under a liberal threshold ($-\log_{10}(p) > 4$) in at least one algorithm. **D.** A comparison of significant regions identified by the 12 algorithms. The bar plot showed the number of regions commonly identified by the algorithms indicated by the black dots. The number of regions identified by each algorithm was presented in the parentheses next to the names of the algorithms.

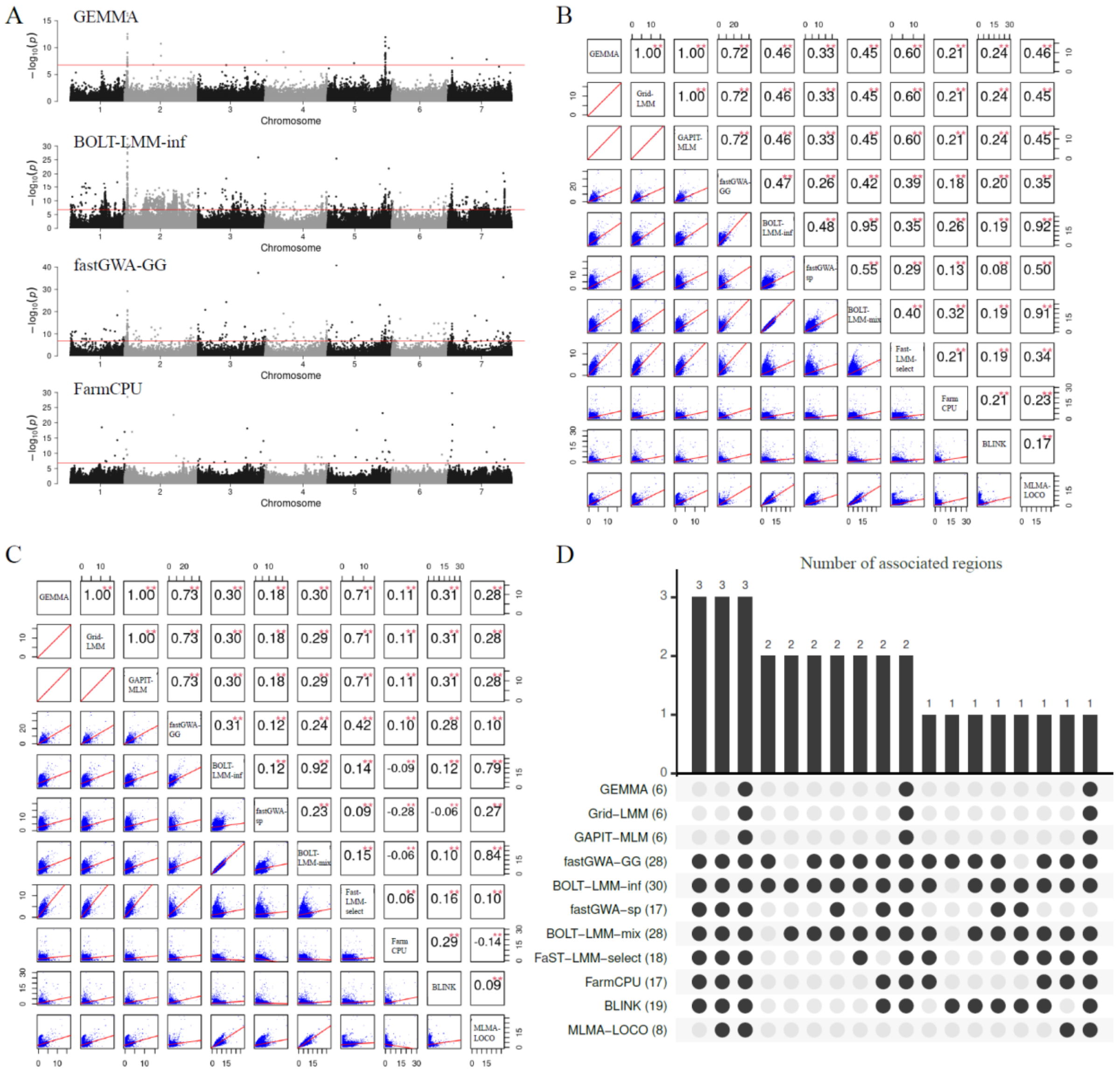

**Supplementary Figure 4.** A comparison of the results of 11 GWAS algorithms for flowering time in a barley data set consisting of 8,825 individuals and 306,049 markers. **A.** The Manhattan plots of 4 selected algorithms. The threshold was $p < 0.05$ after Bonferroni correction. **B.** Correlations between the $-\log_{10}(p)$ values of all markers obtained by each of the indicated pairs of algorithms. The names of the algorithms were indicated in the diagonal blocks. **C.** Pairwise correlations between the $-\log_{10}(p)$ values of markers which were significant under a liberal threshold ($-\log_{10}(p) > 4$) in at least one algorithm. **D.** A comparison of significant regions identified by the 12 algorithms. The bar plot showed the number of regions commonly identified by the algorithms indicated by the black dots. The number of regions identified by each algorithm was presented in the parentheses next to the names of the algorithms. The algorithm CMLM was not evaluated due to the computational load.

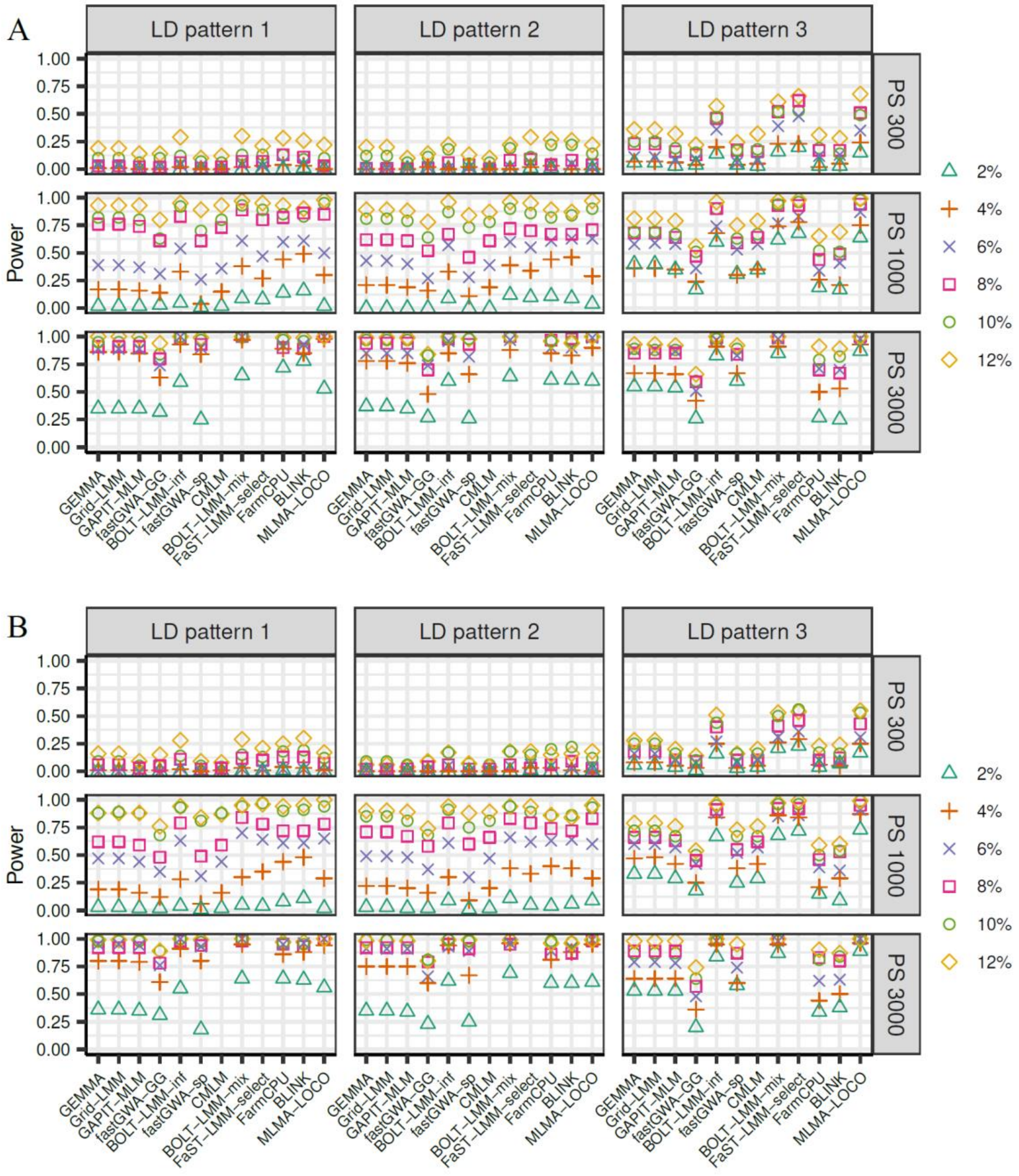

**Supplementary Figure 5.** The statistical power of detecting QTL explaining a specific proportion of genetic variance for 12 GWAS algorithms evaluated in simulated data sets with 18 scenarios for trait heritability 0.5. The 18 scenarios are combinations of three population sizes (PS 300, PS 1000 and PS 3000), three different LD patterns among the QTL (LD patterns 1-3), and two different genetic backgrounds (GB1 and GB2). In LD Pattern 1, there is no LD between any two major QTL or between a major and a minor QTL. In LD Pattern 2, there is no LD between any two major QTL, but LD exits between major and minor QTL. In LD Pattern 3, there exists LD among the major QTL as well as between major and minor QTL. In GB1, there were 1,200 markers as minor QTL. In GB2, all markers on the chromosomes (LD patterns 2 and 3) or on half of the chromosomes (LD pattern 1) contributed as minor QTL. The results for GB1 and GB2 were shown in panel **A** and **B**, respectively. Each panel was further divided into 9 subpanels, each showing the results of a specific combination of population size and data set. Within each subpanel, the results for QTL explaining six different PGs (from 2% to 12% with a step of 2%) were indicated by different symbols. The algorithms CMLM and FaST-LMM-select were not evaluated for PS 3000 because the computational load was too high.

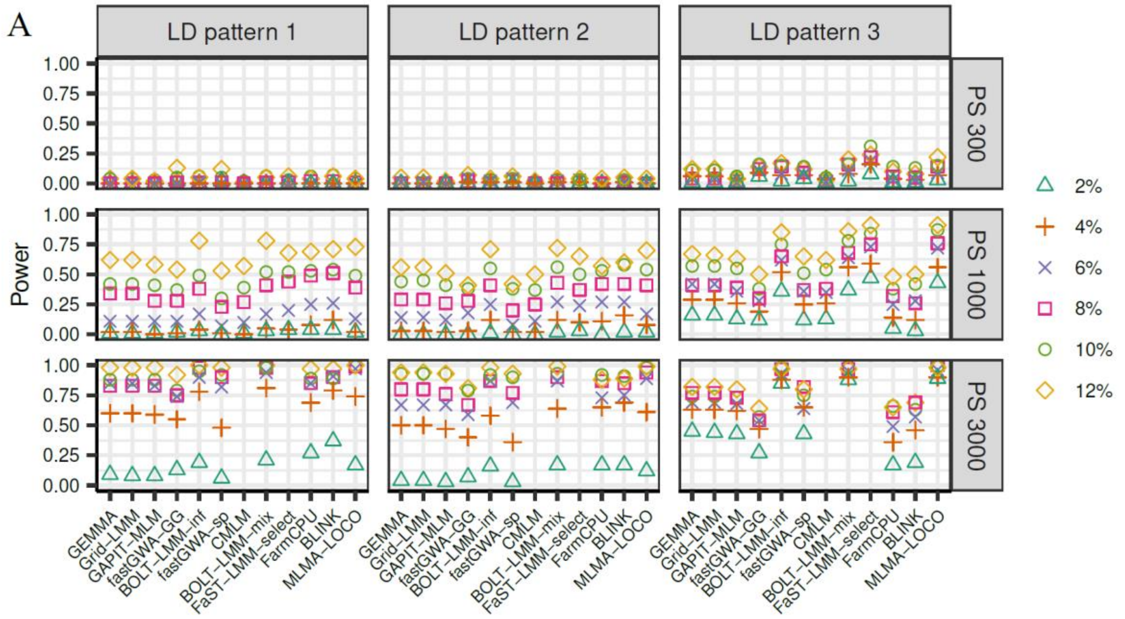

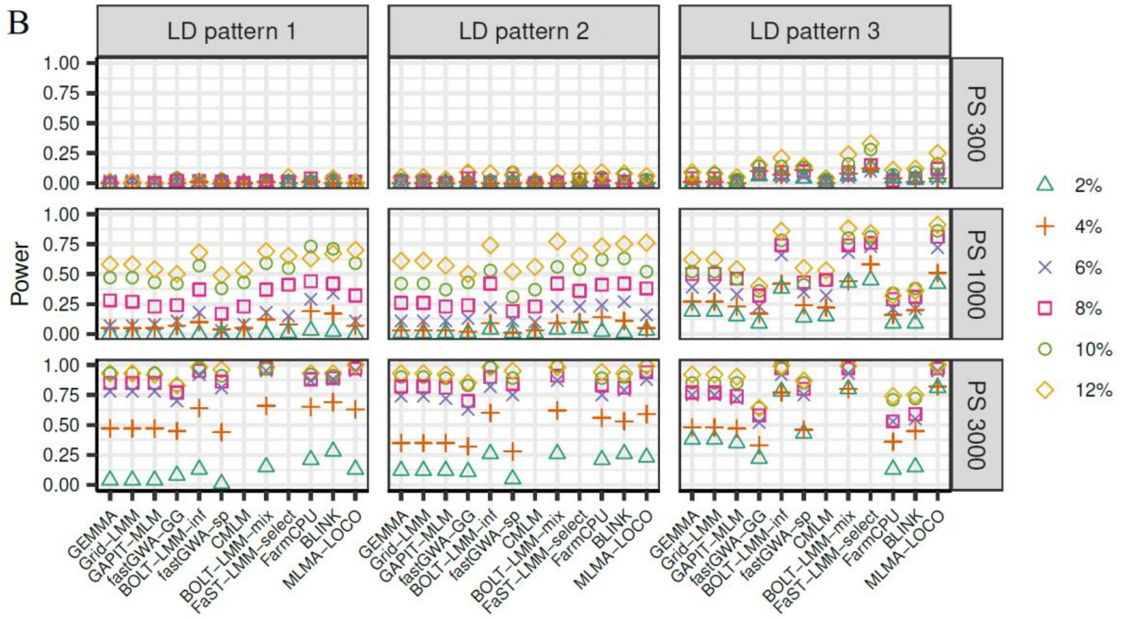

**Supplementary Figure 6.** The statistical power of detecting QTL explaining a specific proportion of genetic variance for 12 GWAS algorithms evaluated in simulated data sets with 18 scenarios for trait heritability 0.3. The 18 scenarios are combinations of three population sizes (PS 300, PS 1000 and PS 3000), three different LD patterns among the QTL (LD patterns 1-3), and two different genetic backgrounds (GB1 and GB2). In LD Pattern 1, there is no LD between any two major QTL or between a major and a minor QTL. In LD Pattern 2, there is no LD between any two major QTL, but LD exits between major and minor QTL. In LD Pattern 3, there exists LD among the major QTL as well as between major and minor QTL. In GB1, there were 1,200 markers as minor QTL. In GB2, all markers on the chromosomes (LD patterns 2 and 3) or on half of the chromosomes (LD pattern 1) contributed as minor QTL. The results for GB1 and GB2 were shown in panel **A** and **B**, respectively. Each panel was further divided into 9 subpanels, each showing the results of a specific combination of population size and data set. Within each subpanel, the results for QTL explaining six different PGs (from 2% to 12% with a step of 2%) were indicated by different symbols. The algorithms CMLM and FaST-LMM-select were not evaluated for PS 3000 because the computational load was too high.

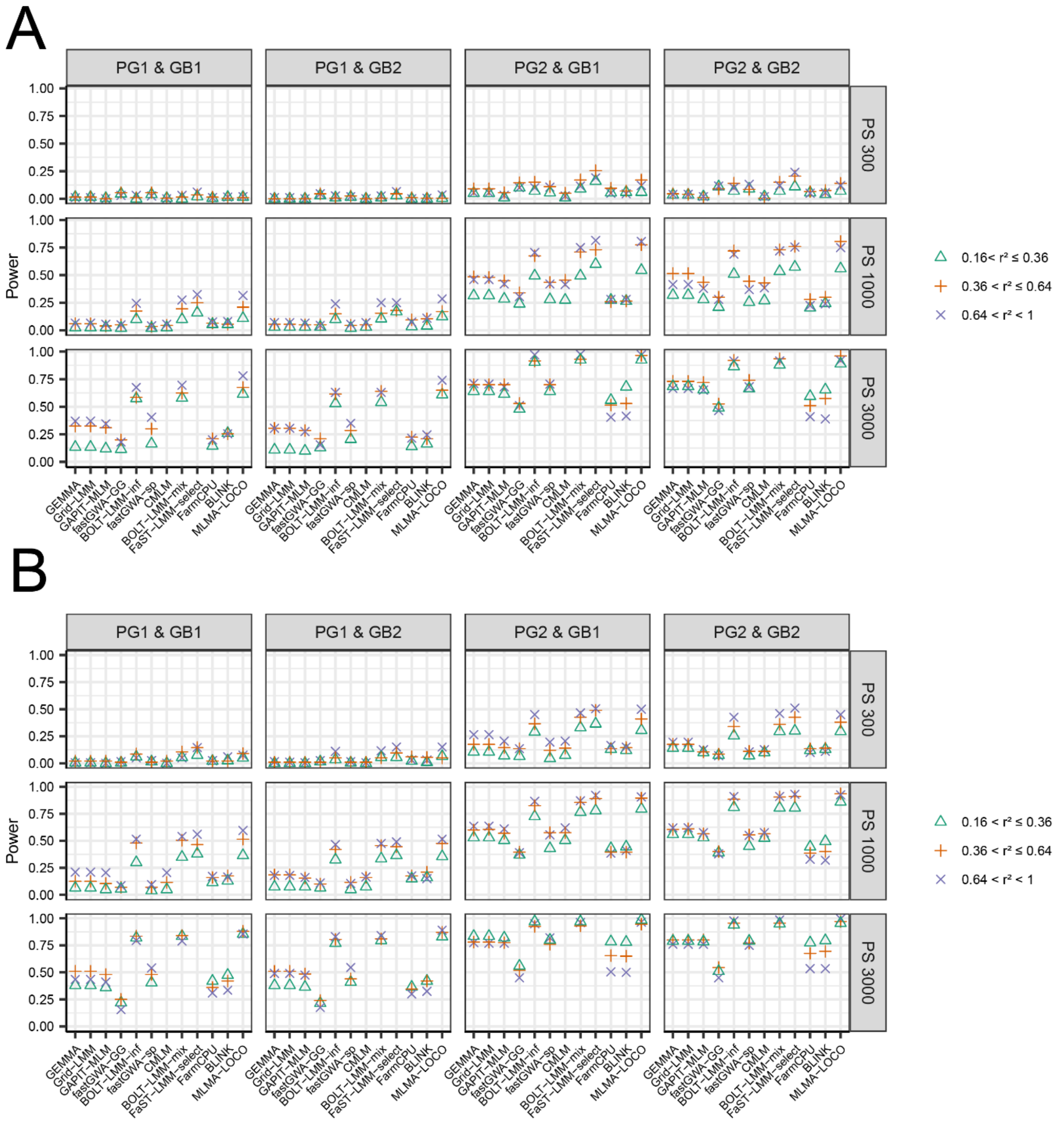

**Supplementary Figure 7.** The statistical power of detecting QTL pairs with a particular range of linkage disequilibrium (LD) for 12 GWAS algorithms evaluated in simulated data sets with 12 scenarios for trait heritability 0.3 (**A**) and 0.5 (**B**). The 12 scenarios are combinations of three population sizes (PS 300, PS 1000 and PS 3000), two patterns of QTL effect sizes (PG1 and PG2), and two different genetic backgrounds (GB1 and GB2). In PG1, each of the 6 major QTL explained 2% of the genetic variance, In PG2, the 6 major QTL were randomly assigned to explain 2%, 4%, 6%, 8%, 10% and 12% of the genetic variance respectively. In GB1, there were 1,200 markers as minor QTL. In GB2, all markers on the chromosomes (LD patterns 2 and 3) or on half of the chromosomes (LD pattern 1) contributed as minor QTL. Each of the 12 subpanels showed the results of a specific combination of population size, PG and GB. Within each subpanel, the results for QTL pairs with three different ranges of LD (measured by $r^2$) were indicated by different symbols. The algorithms CMLM and FaST-LMM-select were not evaluated for PS 3000 because the computational load was too high.

A

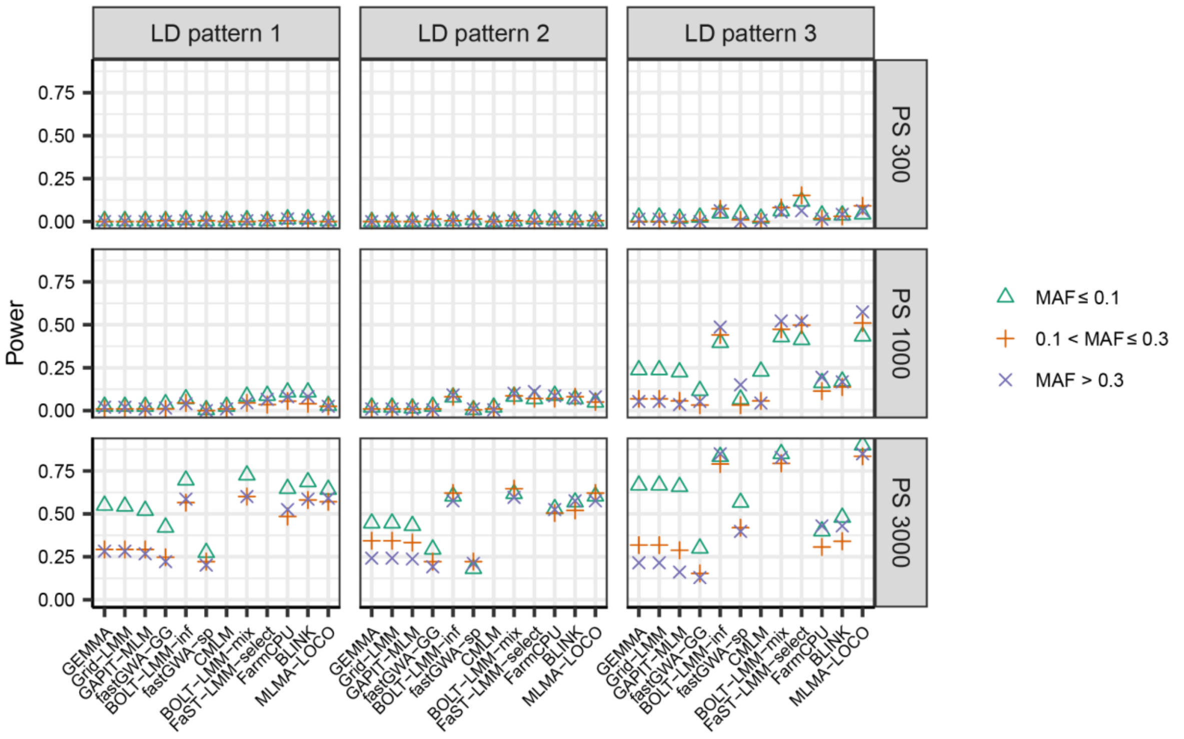

B

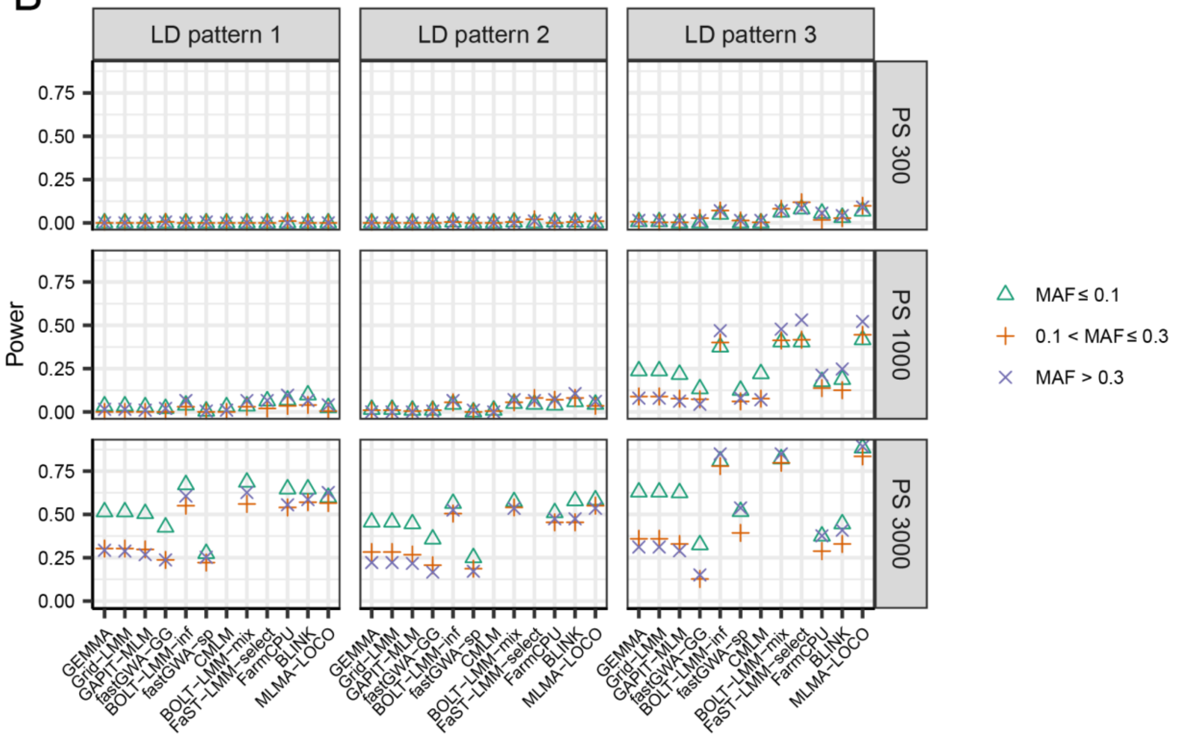

**Supplementary Figure 8.** The statistical power of detecting QTL with a specific range of MAF for 12 GWAS algorithms evaluated in simulated data sets with 18 scenarios for trait heritability 0.5 with PG1 (each of the 6 major QTL explained 2% of the genetic variance). The 18 scenarios are combinations of three population sizes (PS 300, PS 1000 and PS 3000), three different LD patterns among the QTL (LD patterns 1-3), and two different genetic backgrounds (GB1 and GB2). In LD Pattern 1, there is no LD

between any two major QTL or between a major and a minor QTL. In LD Pattern 2, there is no LD between any two major QTL, but LD exits between major and minor QTL. In LD Pattern 3, there exists LD among the major QTL as well as between major and minor QTL. In GB1, there were 1,200 markers as minor QTL. In GB2, all markers on the chromosomes (LD patterns 2 and 3) or on half of the chromosomes (LD pattern 1) contributed as minor QTL. The results for GB1 and GB2 were shown in panel **A** and **B**, respectively. Each panel was further divided into 9 subpanels, each showing the results of a specific combination of population size and LD pattern. Within each subpanel, the results for QTL with three different ranges of MAF were indicated by different symbols. The algorithms CMLM and FaST-LMM-select were not evaluated for PS 3000 because the computational load was too high.

A

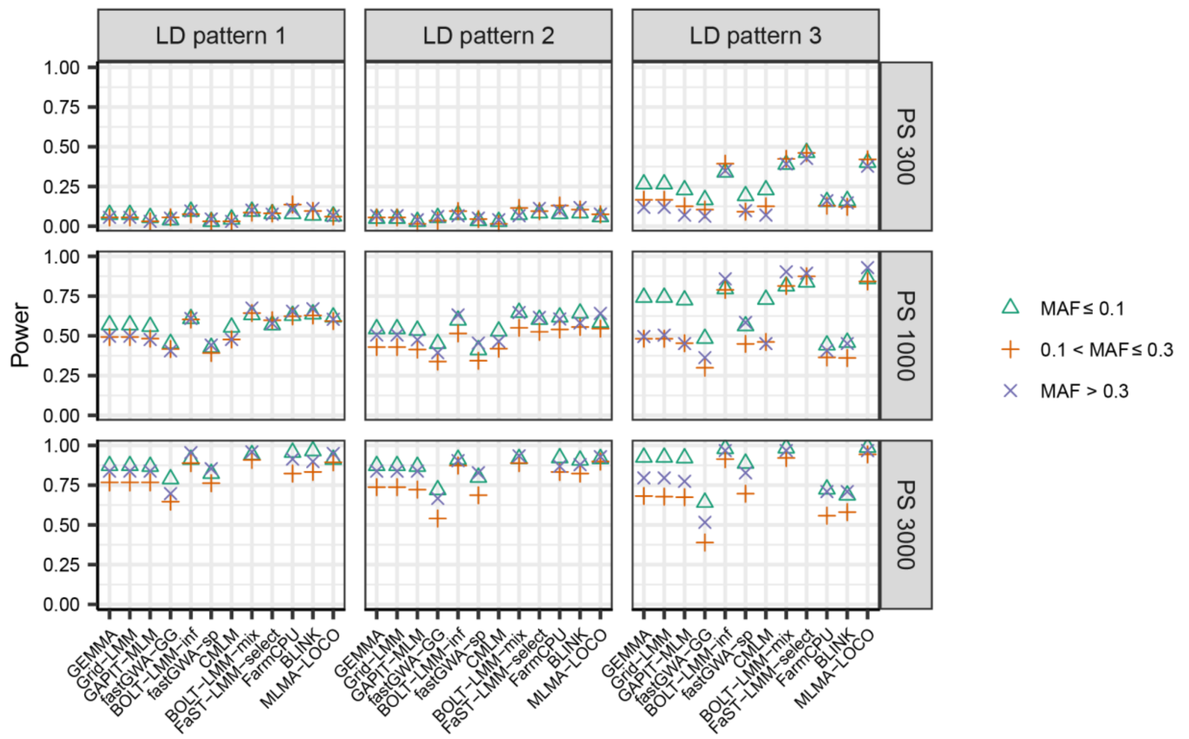

B

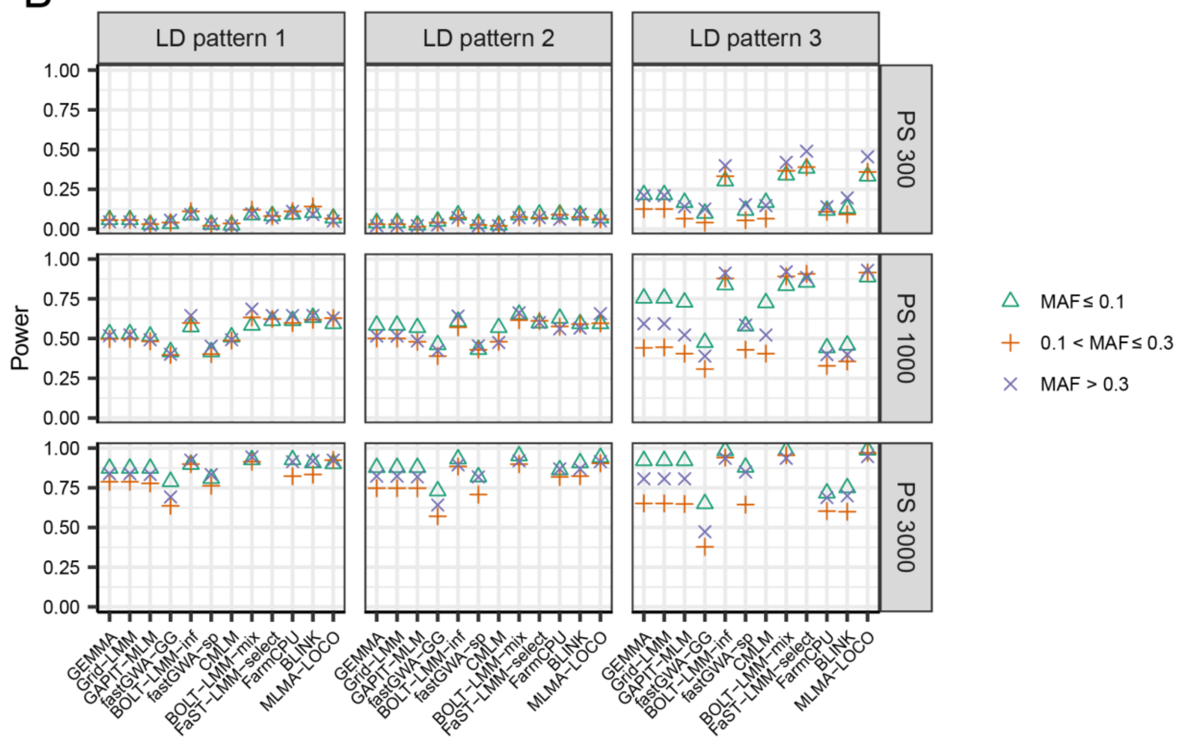

**Supplementary Figure 9.** The statistical power of detecting QTL with a specific range of MAF for 12 GWAS algorithms evaluated in simulated data sets with 18 scenarios for trait heritability 0.5 with PG2 (each of the 6 major QTL explained 2% of the genetic variance). The 18 scenarios are combinations of three population sizes (PS 300, PS 1000 and PS 3000), three different LD patterns among the QTL (LD patterns 1-3), and two different genetic backgrounds (GB1 and GB2). In LD Pattern 1, there is no LD between any two major QTL or between a major and a minor QTL. In LD Pattern 2, there is no LD between any two major QTL, but LD exits between major and minor QTL. In LD Pattern 3, there exists LD among the major QTL as well as between major and minor QTL. In GB1, there were 1,200 markers as minor QTL. In GB2, all markers on the chromosomes (LD patterns 2 and 3) or on half of the chromosomes (LD pattern 1) contributed as minor QTL. The results for GB1 and GB2 were shown in panel **A** and **B**, respectively. Each panel was further divided into 9 subpanels, each showing the results of a specific combination of population size and LD pattern. Within each subpanel, the results for QTL with three different ranges of MAF were indicated by different symbols. The algorithms CMLM and FaST-LMM-select were not evaluated for PS 3000 because the computational load was too high.

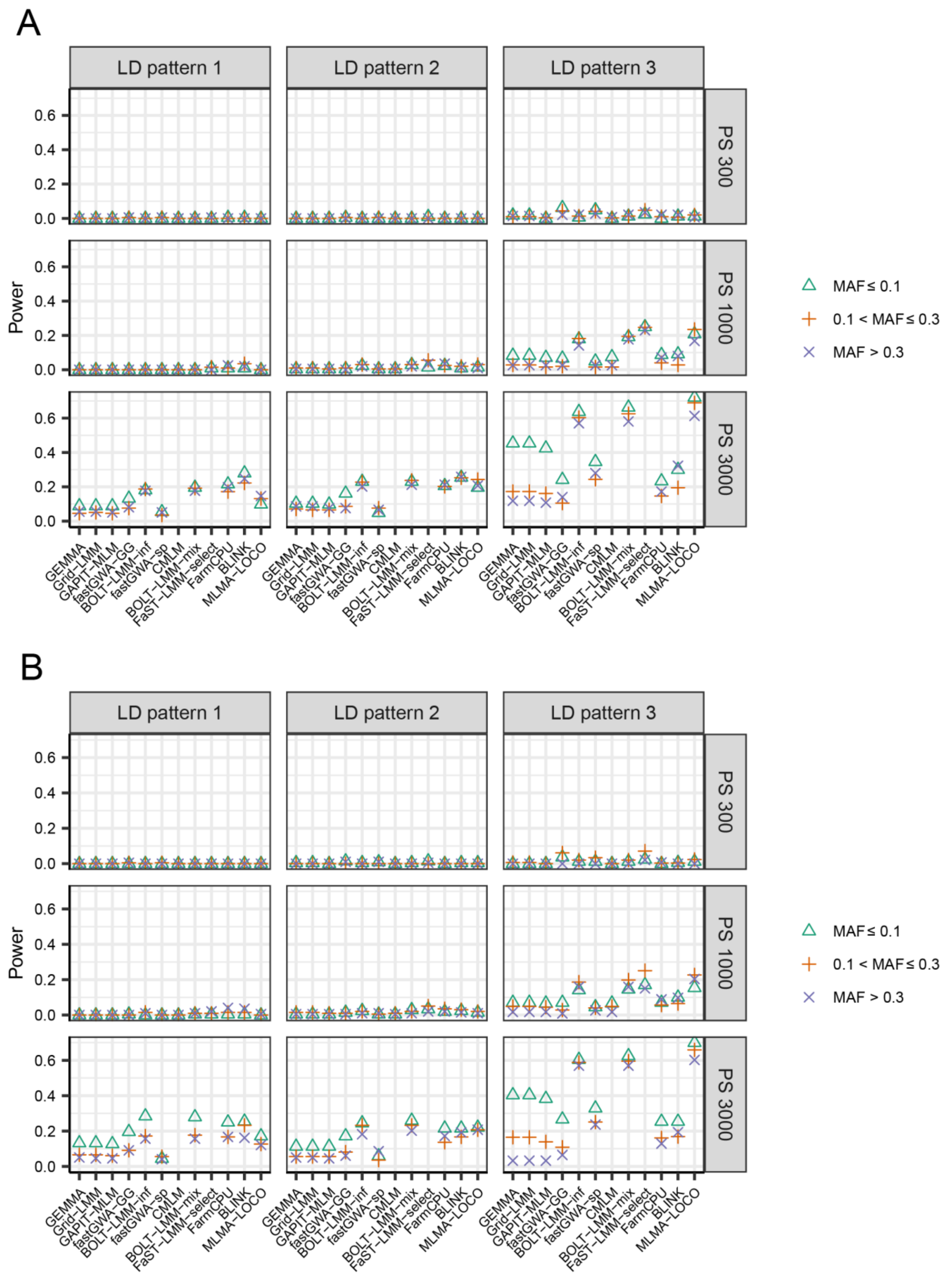

**Supplementary Figure 10.** The statistical power of detecting QTL with a specific range of MAF for 12 GWAS algorithms evaluated in simulated data sets with 18 scenarios for trait heritability 0.3 with PG1 (each of the 6 major QTL explained 2% of the genetic variance). The 18 scenarios are combinations of three population sizes (PS 300, PS 1000 and PS 3000), three different LD patterns among the QTL (LD patterns 1-3), and two different genetic backgrounds (GB1 and GB2). In LD Pattern 1, there is no LD

between any two major QTL or between a major and a minor QTL. In LD Pattern 2, there is no LD between any two major QTL, but LD exits between major and minor QTL. In LD Pattern 3, there exists LD among the major QTL as well as between major and minor QTL. In GB1, there were 1,200 markers as minor QTL. In GB2, all markers on the chromosomes (LD patterns 2 and 3) or on half of the chromosomes (LD pattern 1) contributed as minor QTL. The results for GB1 and GB2 were shown in panel **A** and **B**, respectively. Each panel was further divided into 9 subpanels, each showing the results of a specific combination of population size and LD pattern. Within each subpanel, the results for QTL with three different ranges of MAF were indicated by different symbols. The algorithms CMLM and FaST-LMM-select were not evaluated for PS 3000 because the computational load was too high.

A

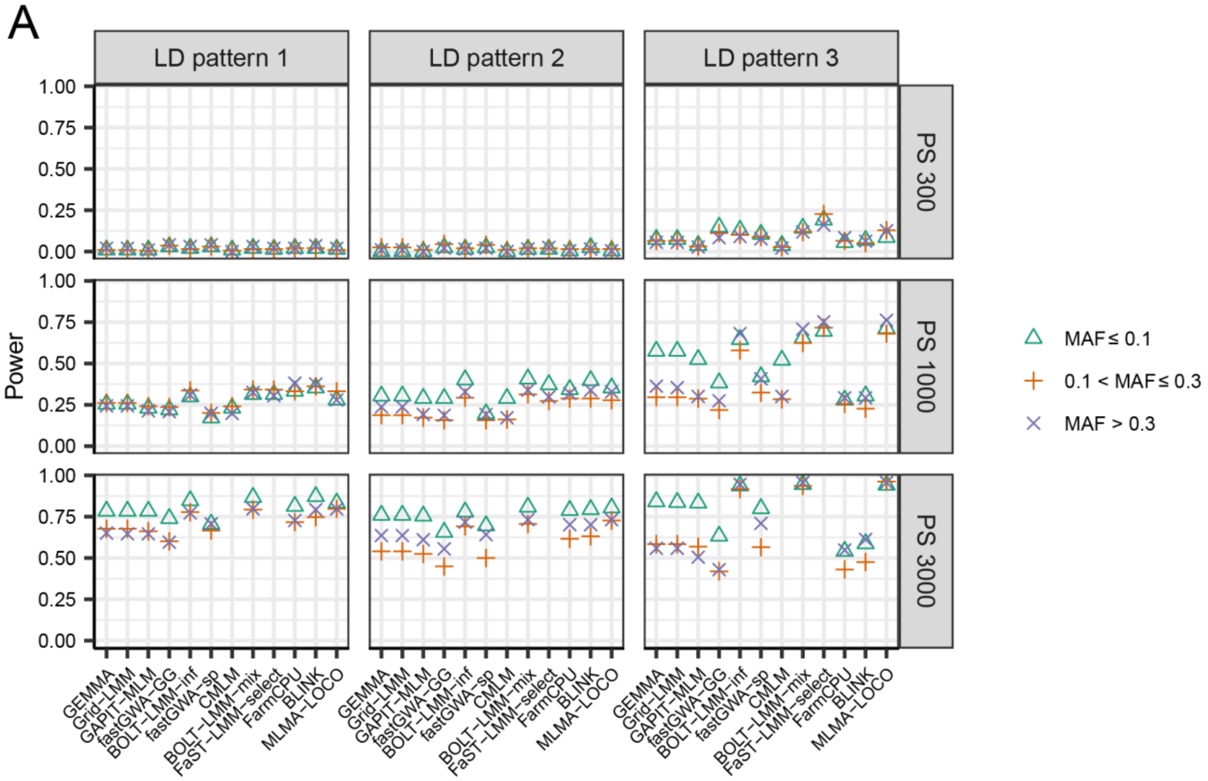

B

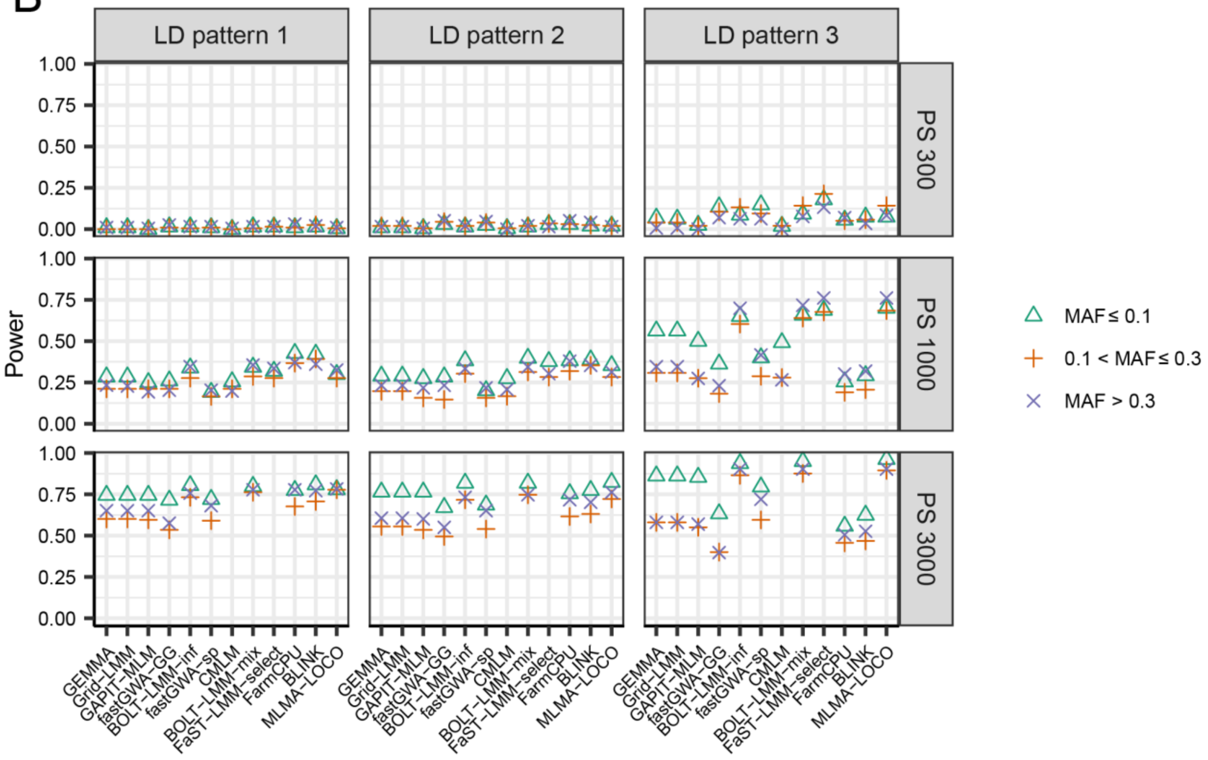

**Supplementary Figure 11.** The statistical power of detecting QTL with a specific range of MAF for 12 GWAS algorithms evaluated in simulated data sets with 18 scenarios for trait heritability 0.3 with PG2 (each of the 6 major QTL explained 2% of the genetic variance). The 18 scenarios are combinations of three population sizes (PS 300, PS 1000 and PS 3000), three different LD patterns among the QTL (LD patterns 1-3), and two different genetic backgrounds (GB1 and GB2). In LD Pattern 1, there is no LD

between any two major QTL or between a major and a minor QTL. In LD Pattern 2, there is no LD between any two major QTL, but LD exits between major and minor QTL. In LD Pattern 3, there exists LD among the major QTL as well as between major and minor QTL. In GB1, there were 1,200 markers as minor QTL. In GB2, all markers on the chromosomes (LD patterns 2 and 3) or on half of the chromosomes (LD pattern 1) contributed as minor QTL. The results for GB1 and GB2 were shown in panel **A** and **B**, respectively. Each panel was further divided into 9 subpanels, each showing the results of a specific combination of population size and LD pattern. Within each subpanel, the results for QTL with three different ranges of MAF were indicated by different symbols. The algorithms CMLM and FaST-LMM-select were not evaluated for PS 3000 because the computational load was too high.

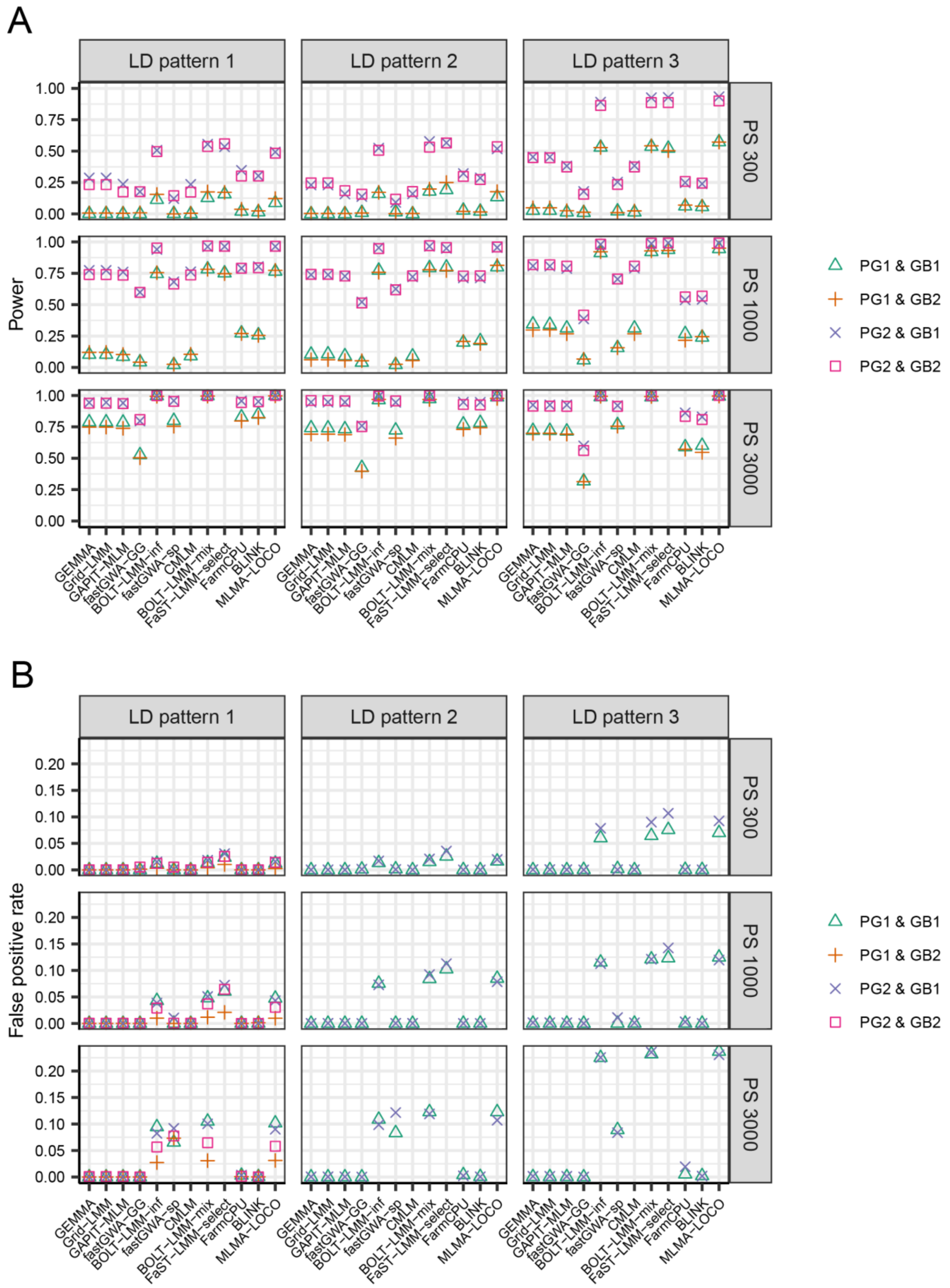

**Supplementary Figure 12.** The statistical power (**A**) and false positive rate (**B**) of 12 GWAS algorithms evaluated in simulated data sets with 36 scenarios for heritability 0.7, under the threshold of $p < 0.05$ after Benjamini-Hochberg correction for multiple testing. The 36 scenarios are combinations of three population sizes (PS 300, PS 1000 and PS 3000), three different LD patterns among the QTL (LD

patterns 1-3), two patterns of QTL effect sizes (PG1 and PG2), and two different genetic backgrounds (GB1 and GB2). In LD pattern 1, there is no LD between any two major QTL or between a major and a minor QTL. In LD pattern 2, there is no LD between any two major QTL, but LD exits between major and minor QTL. In LD pattern 3, there exists LD among the major QTL as well as between major and minor QTL. In PG1, each of the 6 major QTL explained 2% of the genetic variance, In PG2, the 6 major QTL were randomly assigned to explain 2%, 4%, 6%, 8%, 10% and 12% of the genetic variance respectively. In GB1, there were 1,200 markers as minor QTL. In GB2, all markers on the chromosomes (LD patterns 2 and 3) or on half of the chromosomes (LD pattern 1) contributed as minor QTL. Each of the 9 subpanels showed the results of a specific combination of population size and data set. Within each subpanel, the results of four combinations of two PGs and two GBs were indicated by different symbols. The algorithms CMLM and FaST-LMM-select were not evaluated for PS 3000 because the computational load was too high.

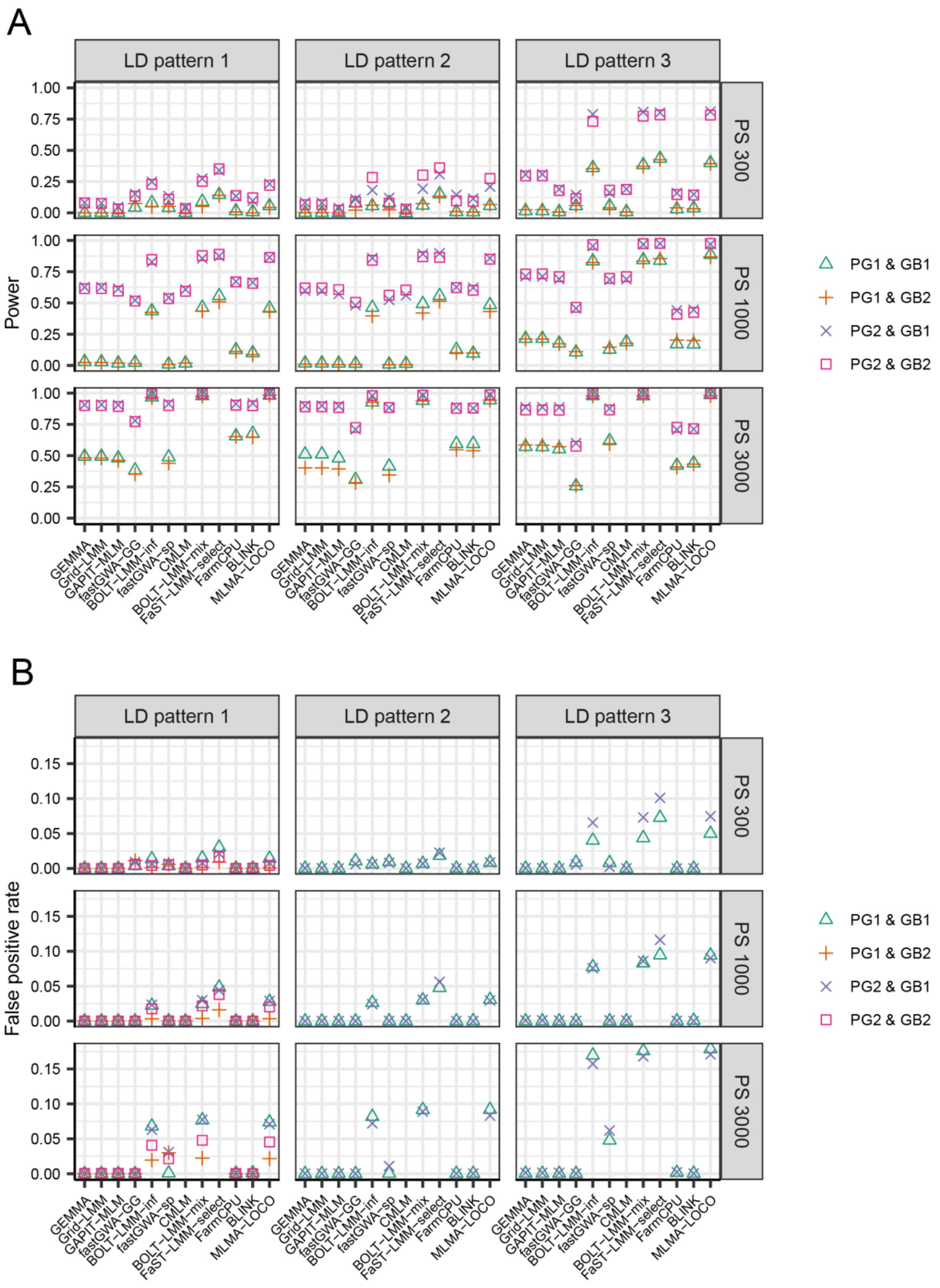

**Supplementary Figure 13.** The statistical power (**A**) and false positive rate (**B**) of 12 GWAS algorithms evaluated in simulated data sets with 36 scenarios for heritability 0.5, under the threshold of $p < 0.05$ after Benjamini-Hochberg correction for multiple testing. The 36 scenarios are combinations of three population sizes (PS 300, PS 1000 and PS 3000), three different LD patterns among the QTL (LD

patterns 1-3), two patterns of QTL effect sizes (PG1 and PG2), and two different genetic backgrounds (GB1 and GB2). In LD pattern 1, there is no LD between any two major QTL or between a major and a minor QTL. In LD pattern 2, there is no LD between any two major QTL, but LD exits between major and minor QTL. In LD pattern 3, there exists LD among the major QTL as well as between major and minor QTL. In PG1, each of the 6 major QTL explained 2% of the genetic variance, In PG2, the 6 major QTL were randomly assigned to explain 2%, 4%, 6%, 8%, 10% and 12% of the genetic variance respectively. In GB1, there were 1,200 markers as minor QTL. In GB2, all markers on the chromosomes (LD patterns 2 and 3) or on half of the chromosomes (LD pattern 1) contributed as minor QTL. Each of the 9 subpanels showed the results of a specific combination of population size and data set. Within each subpanel, the results of four combinations of two PGs and two GBs were indicated by different symbols. The algorithms CMLM and FaST-LMM-select were not evaluated for PS 3000 because the computational load was too high.

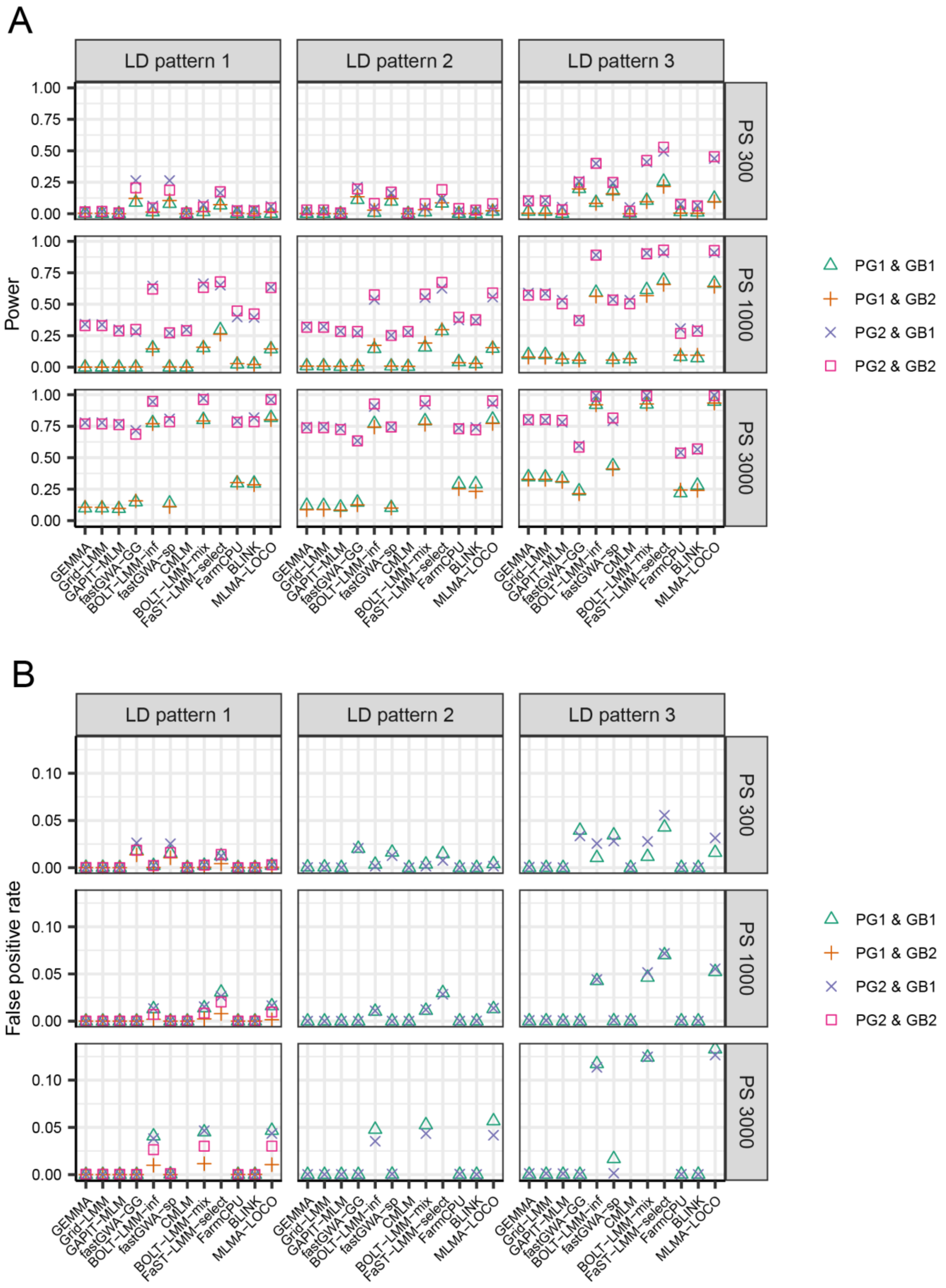

**Supplementary Figure 14.** The statistical power (**A**) and false positive rate (**B**) of 12 GWAS algorithms evaluated in simulated data sets with 36 scenarios for heritability 0.3, under the threshold of $p < 0.05$ after Benjamini-Hochberg correction for multiple testing. The 36 scenarios are combinations of three population sizes (PS 300, PS 1000 and PS 3000), three different LD patterns among the QTL (LD

patterns 1-3), two patterns of QTL effect sizes (PG1 and PG2), and two different genetic backgrounds (GB1 and GB2). In LD pattern 1, there is no LD between any two major QTL or between a major and a minor QTL. In LD pattern 2, there is no LD between any two major QTL, but LD exits between major and minor QTL. In LD pattern 3, there exists LD among the major QTL as well as between major and minor QTL. In PG1, each of the 6 major QTL explained 2% of the genetic variance, In PG2, the 6 major QTL were randomly assigned to explain 2%, 4%, 6%, 8%, 10% and 12% of the genetic variance respectively. In GB1, there were 1,200 markers as minor QTL. In GB2, all markers on the chromosomes (LD patterns 2 and 3) or on half of the chromosomes (LD pattern 1) contributed as minor QTL. Each of the 9 subpanels showed the results of a specific combination of population size and data set. Within each subpanel, the results of four combinations of two PGs and two GBs were indicated by different symbols. The algorithms CMLM and FaST-LMM-select were not evaluated for PS 3000 because the computational load was too high.

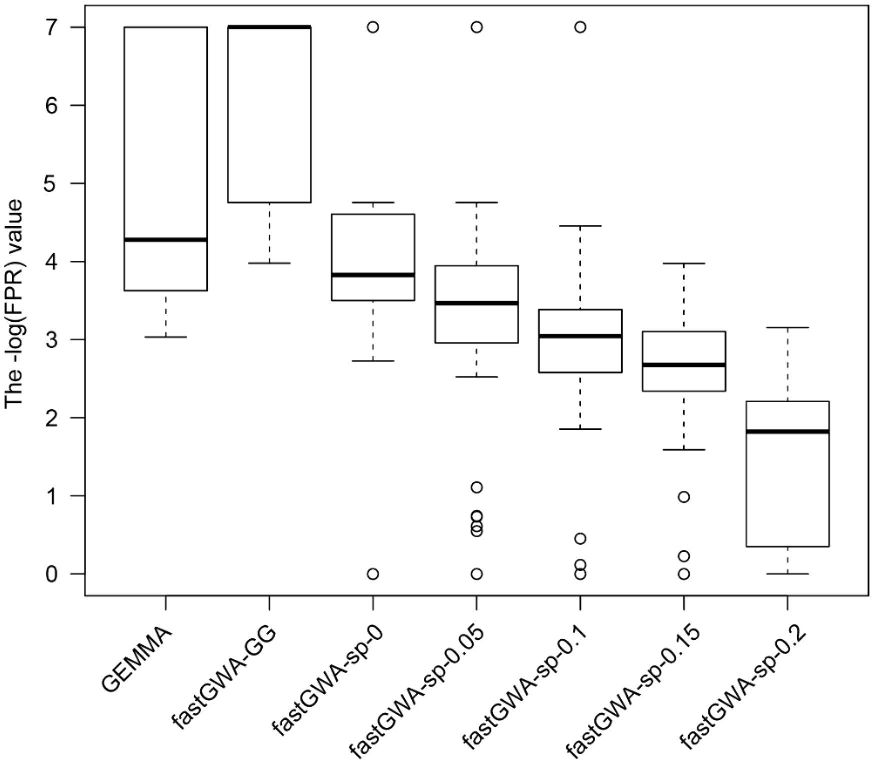

**Supplementary Figure 15.** The distribution of false positive rate (FPR) of fastGWA-sp with different thresholds (from 0 to 0.2 with a step of 0.05) for generating the sparse kinship matrix, evaluated in 100 simulated phenotypes. To make the variation more clearly visible, a very small value ($1 \times 10^{-7}$) was added to the FPR values to allow log-transformation of all FPR values. The population size of the simulated data was 3,000 with a trait heritability of 0.7. There were six major QTL each explaining 2% of the genetic variance. 1,200 markers contributed as minor QTL to the genetic background effects. The FPR of the algorithm GEMMA and fastGWA-GG were presented as a benchmark for comparison. Note that fastGWA-GG implemented the GRAMMAR-Gamma approximation, but with the exact kinship matrix.

**Supplementary Table 2.** The average false positive rate (FPR) of six algorithms evaluated in four scenarios, each consisting of 100 simulated phenotypes. In all scenarios, there were six major QTL explaining from 2% to 12% (with a step of 2%) proportion of genetic variance (PG2), and 1,200 markers contributed as minor QTL to the genetic background effects (GB1). The trait heritability was 0.7. The population size was 300 for two scenarios (PS300), and 1000 for the other two (PS1000). In LD pattern 2, there is no LD between any two major QTL, but LD exits between major and minor QTL. In LD pattern 3, there exists LD among the major QTL as well as between major and minor QTL. BOLT-LMM-inf-GLOCO and BOLT-LMM-mix-GLOCO are variants of BOLT-LMM-inf and BOLT-LMM-mix, where a genuine leave-one-chromosome-out (LOCO) procedure was forced.

| Algorithm | PS300 | | PS1000 | |
|---|---|---|---|---|
| | LD pattern 2 | LD pattern 3 | LD pattern 2 | LD pattern 3 |
| GEMMA | $2.06 \times 10^{-5}$ | $2.05 \times 10^{-4}$ | $1.18 \times 10^{-4}$ | $6.29 \times 10^{-4}$ |
| GAPIT-MLM | $4.19 \times 10^{-6}$ | $1.12 \times 10^{-4}$ | $9.00 \times 10^{-5}$ | $5.32 \times 10^{-4}$ |
| BOLT-LMM-inf | $1.63 \times 10^{-3}$ | $1.86 \times 10^{-2}$ | $1.17 \times 10^{-2}$ | $3.65 \times 10^{-2}$ |
| BOLT-LMM-mix | $2.14 \times 10^{-3}$ | $2.03 \times 10^{-2}$ | $1.74 \times 10^{-2}$ | $3.80 \times 10^{-2}$ |
| BOLT-LMM-inf-GLOCO | $1.64 \times 10^{-3}$ | $1.85 \times 10^{-2}$ | $1.15 \times 10^{-2}$ | $3.59 \times 10^{-2}$ |
| BOLT-LMM-mix-GLOCO | $1.91 \times 10^{-3}$ | $2.05 \times 10^{-2}$ | $1.60 \times 10^{-2}$ | $1.48 \times 10^{-2}$ |

**Supplementary Table 3.** The power and false positive rate (FPR) of three variants of the algorithm CMLM evaluated in two scenarios, each consisting of 100 simulated phenotypes. In both scenarios, there were six major QTL explaining from 2% to 12% (with a step of 2%) proportion of genetic variance (PG2), and 1,200 markers contributed as minor QTL to the genetic background effects (GB1). The trait heritability was 0.7 and the population size was 1000. In LD pattern 2, there is no LD between any two major QTL, but LD exits between major and minor QTL. In LD pattern 3, there exists LD among the major QTL as well as between major and minor QTL. CMLM_op: the default setting of CMLM in which the compression level is optimized. CMLM_5: fixing the rate of compression to 5, namely clustering the individuals into groups of 5 individuals. CMLM_10: fixing the rate of compression to 10.

| Algorithm | LD pattern 2 | | LD pattern 3 | |
|---|---|---|---|---|
| | Power | FPR | Power | FPR |
| CMLM_op | 0.623 | $8.96 \times 10^{-5}$ | 0.668 | $5.33 \times 10^{-4}$ |
| CMLM_5 | 0.742 | $2.31 \times 10^{-2}$ | 0.867 | $2.98 \times 10^{-2}$ |
| CMLM_10 | 0.733 | $1.17 \times 10^{-2}$ | 0.847 | $1.57 \times 10^{-2}$ |
